# mRNA lipid nanoparticle-incorporated nanofiber-hydrogel composite generates a local immunostimulatory niche for cancer immunotherapy

**DOI:** 10.1101/2025.01.27.633179

**Authors:** Yining Zhu, Zhi-Cheng Yao, Shuyi Li, Jingyao Ma, Christine Wei, Di Yu, Jessica L. Stelzel, Bobby Y.X. Ni, Yang Miao, Kyra Van Batavia, Xiaoya Lu, Jinghan Lin, Yifan Dai, Jiayuan Kong, Ruochen Shen, Kailei D. Goodier, Xiang Liu, Leonardo Cheng, Ivan Vuong, Gregory P. Howard, Natalie K. Livingston, Joseph Choy, Jonathan P. Schneck, Joshua C. Doloff, Sashank K. Reddy, John W. Hickey, Hai-Quan Mao

**Affiliations:** Department of Biomedical Engineering, Johns Hopkins University School of Medicine, Baltimore, MD, USA; Institute for NanoBioTechnology, Johns Hopkins University, Baltimore, MD, USA; Translational Tissue Engineering Center, Johns Hopkins University School of Medicine, Baltimore, MD, USA; Department of Materials Science and Engineering, Johns Hopkins University, Baltimore, MD, USA; Department of Biomedical Engineering, Duke University, Durham, NC, USA; Department of Biostatistics, Gillings School of Global Public Health, University of North Carolina at Chapel Hill, Chapel Hill, NC, USA; Institute for Cell Engineering, Johns Hopkins University School of Medicine, Baltimore, MD, USA; Department of Pathology, Johns Hopkins University School of Medicine, Baltimore, MD, USA; Department of Medicine, Johns Hopkins University School of Medicine, Baltimore, MD, USA; Department of Oncology, Sidney Kimmel Comprehensive Cancer Center and the Bloomberg-Kimmel Institute for Cancer Immunotherapy, Johns Hopkins University School of Medicine, Baltimore, MD, USA; Department of Plastic and Reconstructive Surgery, Johns Hopkins University School of Medicine, Baltimore, MD, USA

## Abstract

Hydrogel materials have emerged as versatile platforms for various biomedical applications. Notably, the engineered nanofiber-hydrogel composite (NHC) has proven effective in mimicking the soft tissue extracellular matrix, facilitating substantial recruitment of host immune cells and the formation of a local immunostimulatory microenvironment. Leveraging this feature, here we report an mRNA lipid nanoparticle (LNP)-incorporated NHC microgel matrix, termed LiNx, by incorporating LNPs loaded with mRNA encoding tumour antigens. Harnessing the potent transfection efficiency of LNPs in antigen-presenting cells (APCs), LiNx demonstrates remarkable immune cell recruitment, antigen expression and presentation, and cellular interaction. These attributes collectively create an immunostimulating milieu and yield a potent immune response achievable with a single dose, comparable to the conventional three-dose LNP immunization regimen. Further investigations reveal that the LiNx not only generates heightened Th1 and Th2 responses but also elicits a distinctive Type 17 T helper cell-mediated response pivotal for bolstering antitumour efficacy. Our findings elucidate the mechanism underlying LiNx’s role in potentiating antigen-specific immune responses, presenting a new strategy for cancer immunotherapy.

## INTRODUCTION

Hydrogels have been increasingly proposed as delivery carriers for vaccines through various modes of action, such as sustained release of antigens and/or adjuvants, recruitment of host cells to the site of injection, enhancement of the interaction of antigens and immune cells, etc.^1,2^ The hydrogel-based vaccine delivery platform is capable of orchestrating an immunostimulatory niche if the key cellular components are present in the local microenvironment.^3,4^ Such a delivery system has a strong potential in augmenting immune responses.^5–7^ We have recently developed a biostimulatory nanofiber-hydrogel composite (NHC) that mimics the microarchitecture and mechanical properties of the extracellular matrix of soft tissues.^8,9^ The NHC has an integrated composite structure where electrospun poly(ε-caprolactone) (PCL) nanofiber fragments are covalently crosslinked to a hyaluronic acid (HA) network through covalent interfacial bonding. The NHC offers tunable stiffness, pore size, and degradation duration by adjusting HA structure and concentration, crosslinker concentration, and nanofiber concentration.^10^ A highly porous NHC with a low shear storage modulus (G’: 150–350 Pa) and 1–3 w/v% PCL nanofibers showed a high level of biostimulatory, proangiogenic, and pro-regenerative properties without using any additional exogenous biochemical cues.^11^ Using a simple mechanical fragmentation method, a particulated microgel form of the NHC can be generated for ease of injection. These NHC microgels promoted host cell recruitment, immunomodulation, angiogenesis, soft tissue remodeling, and tissue repair. Moreover, the unique property of recruiting a significant number of immune cells to form a local immunostimulatory microenvironment after subcutaneous (*s.c.*) or intramuscular (*i.m.*) injection opens up exciting possibilities. With the advent of newly developed mRNA vaccines,^12^ leveraging this property can potentiate efficacy by modulating the transfection of the local cell types and coordinating the recruited host immune cells.

The success of two mRNA vaccines, Spikevax^®^ (Moderna) and Comirnaty^®^ (BioNTech/Pfizer),^13^ in combating SARS-CoV-2 during the coronavirus pandemic demonstrates the safety and efficacy of the lipid nanoparticle (LNP) delivery platform for mRNA vaccines and provides impetus to the rapid expansion of mRNA LNP-based cancer vaccines and cell therapy.^14^ These LNP vaccines achieve protection by eliciting high antibody titers, memory B cell, and T follicular helper cell responses;^15^ some evidence also suggests the involvement of IFN-γ^+^ or IL2^+^CD8^+^ T cells and CD4^+^ Th1 cells.^16^ The collective findings from Phase 1/2 clinical studies of several mRNA vaccines for high-risk melanoma,^17^ non-small-cell lung cancer,^18^ pancreatic cancer,^19^ and other cancers have demonstrated the feasibility of mRNA-based cancer vaccines.^20^ The convergence of state-of-the-art LNP technology with personalized medicine approaches amplifies the optimism surrounding this strategy.^21^ Nonetheless, realizing the full potential of mRNA LNP cancer vaccines requires further optimization of LNP immune activation profiles and delivery strategies to permit more efficient programming of immune responses for potent anticancer efficacy.^16^

Recent literature reports that not only the choice of lipid components but also the relative proportions of the lipid ingredients in the formulation greatly influence the *in vivo* transfection efficiency and delivery outcomes.^16,22,23^ To systematically optimize LNP formulations for cell-preferential transfection, we developed a step-wise screening method that combines *in vitro* and *in vivo* assessment steps using a cohort of 1,080 LNP formulations prepared by varying molar ratios of luciferase pDNA or mRNA, ionizable lipid (DLin-MC3-DMA), cholesterol, DMG-PEG2000, and more importantly, a helper lipid selected from those that were previously used in FDA-approved or experimental LNP formulations.^22^ The helper lipids were selected to represent various charge features, including cationic [1,2-dioleoyl-3-trimethylammonium-propane (DOTAP) and dimethyl dioctadecyl ammonium (DDAB)], zwitterionic [1,2-dioleoyl-sn-glycero-3-phosphoethanolamine (DOPE) and 1,2-distearoyl-sn-glycero-3-phosphocholine (DSPC)], and anionic [1,2-dimyristoyl-sn-glycero-3-phosphate (14PA) and 1-stearoyl-2-oleoyl-sn-glycero-3-phospho-(1’-rac-glycerol) (18PG)].^22^ The LNPs demonstrating the highest levels of *in vitro* transfection were initially identified through screening tests, followed by individual formulation assessments to pinpoint the most effective LNPs. This screening identified three top-performing LNPs (C10, D6, and F5) based on transfection efficiency in primary bone marrow-derived dendritic cells (BMDCs).^16^ Following three doses of *s.c.* injections, these three LNPs induced comparably potent antigen-specific Th1 responses but substantially different Th2 responses.^16^ All three formulations showed significant tumour suppression and markedly prolonged survival in a prophylactic model of OVA-expressing melanoma in C57BL/6 mice.^16^ Compared with Th1-biased F5 and D6 LNPs, a dual Th1/Th2 activating C10 LNPs induced the highest level of potency in suppressing tumour growth and highest survival when tested in therapeutic melanoma models.^16^ Interestingly, the different levels of Th2 responses generated by these 3 selected LNP formulations also correlated with their distinct transfection abilities in non-APCs such as myoblasts. These data revealed that intricate tuning of the LNP composition will enable us to alter transgene expression in APCs and non-APCs when delivering antigen-encoding mRNA *in vivo*. Further, by biasing cell-preferential gene expression, we altered the immune activation profile elicited by these LNPs.^16^

Building on these findings, we aim to leverage the immunostimulatory microenvironment generated by NHC^9–11^ and the potent transfection efficiency and immune-activation capability of LNPs^16,22^ by incorporating LNPs into the NHC composite. This mRNA <u>L</u>NP-incorporated <u>N</u>HC microgel matri<u>x</u>, termed LiNx, may create a conducive niche to enhance the transfection of APCs and non-APCs at the injection site, and, more importantly, facilitate the interactions and signaling of all relevant immune cell types in the local niche (**Fig. 1a**). In this study, we prepared three different LiNx formulations with the top three LNPs selected from our library and examined the effect of the microgel niche on cell recruitment activity and transfection efficiency among different cell types. We then investigated antigen presentation, immune cell activation in different LiNx groups, and antigen-specific immune responses. More importantly, we assessed the efficacy of a single immunization with LiNx in comparison to the conventional three-dose prime-and-boost regimen for free LNPs. Furthermore, we validated the synergistic effects of combining the optimized mRNA LiNx cancer vaccine with systemic immune checkpoint blockade in therapeutic cancer models.

**Figure 1.**
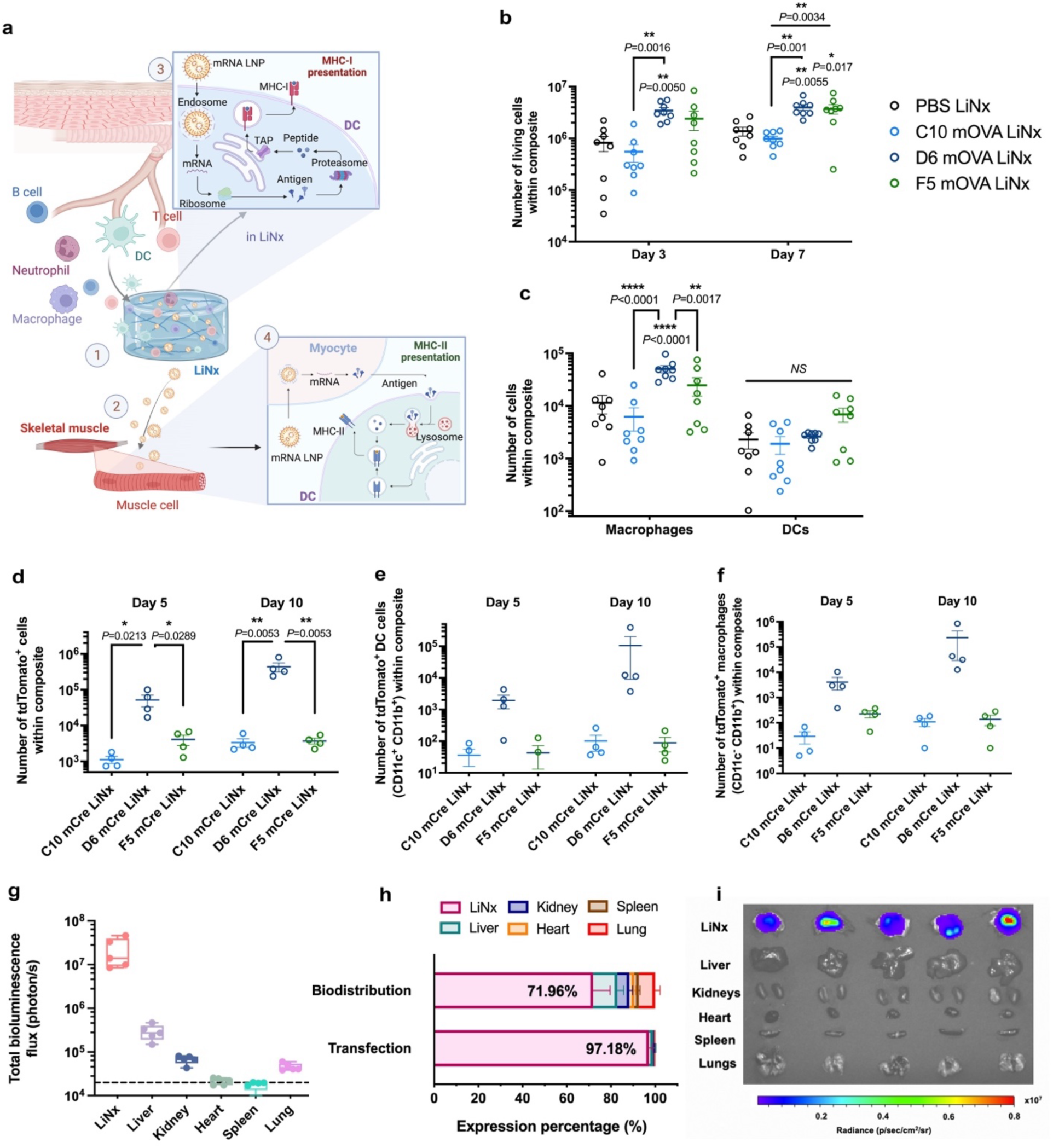
*In vivo* host cell recruitment and transfection efficiency of LiNx. **a,** Schematic of the LiNx vaccination platform, providing an immunostimulatory niche by effectively recruiting host immune cells to the microgel matrix (1). LiNx-delivered LNPs can preferentially transfect non-APCs (2) and recruited macrophage-like cells, followed by antigen expression, processing, and epitope presentation on MHC-I (3) or MHC-II (4) in different pathways. **b-c,** Kinetics of host cell recruitment following injection of three different LNP/mRNA LiNx in C57BL/6 mice (30 μg mOVA per mouse, *s.c.*). The Cellaca MX High-throughput Automated Cell Counter was used to count the number of living cells within composites on both day 3 and day 7 post-injection (b). Flow cytometry was employed to determine the count of macrophage-like cells (CD3^-^CD11b^+^Ly6g^-^CD11c^-^) and DC-like cells (CD3^-^ CD11b^+^ Ly6g^-^CD11c^+^) on day 3 (c). **d-f,** Ai9 mice were administered the three LiNx loaded with mCre via *s.c.* injections (30 μg mCre per mouse). Transfection of cells in the composite was analysed by flow cytometry. The number of cells positive for tdTomato **(d)**, as well as DC-like cells (CD11b^+^CD11c^+^) cells **(e)** and macrophage-like cells (CD11b^+^CD11c^-^) **(f)** positive for tdTomato on day 5 and day 10 post-injection were shown. **g-i**, C57BL/6 mice received *s.c.* injections of D6 LiNx loaded with either mLuc or Cy5-labeled mRNA (30 μg per mouse). **(g)** The transfection efficiency of D6 LiNx at the injection site and in major organs was assessed using IVIS imaging. The relative percentage of transfection and biodistribution for D6 LiNx is shown in **(h)**. IVIS images of luciferase transfection for D6 LiNx are presented in **(i)**. Data represent the mean ± s.e.m. (n = 8 for **(b-c)**, n = 4 for **(d-f)** and n = 5 for **(g-i)** biologically independent samples). Data were analysed using one-way ANOVA and Tukey’s multiple comparisons test. \**P* < 0.05, \*\**P* < 0.01, \*\*\*\**P* < 0.0001. NS, not significant.

## RESULTS

### Host cell recruitment and transfection profile of mRNA LiNx

The three top-performing mRNA LNP formulations (C10, D6, and F5 LNPs, **Supplementary Table 1**) identified from our previous study were selected to prepare the LiNx.^16^ All three mRNA LNPs led to high levels of expression of the OVA-derived SIINFEKL peptide on major histocompatibility complex class I (MHC-I) on BMDCs and enhanced maturation of BMDCs, marking them as promising candidates for exploration in the context of the LiNx. The NHC microgel with a shear storage modulus G′ of 250 Pa was chosen for its previously reported effectiveness in cell recruitment, migration, and retention (**Supplementary Fig. 1a**). We mixed the NHC microgel particles with mRNA LNPs using the syringe extrusion method immediately before injection (**Supplementary Fig. 1b**). Scanning electron microscopy images revealed that, after encapsulating LNPs within the NHC, the same fibrillar microarchitecture of the NHC is observed, with nanofibers entrapped within the HA hydrogel network (**Supplementary Fig. 2**). The accelerated stability test of NHC at 50°C showed that, at this temperature, NHC stiffness was reduced by approximately 40% after 7 days and around 70% after 14 days (**Supplementary Fig. 3**). Based on a scaling factor calculated empirically, the NHC scaffold used in this study can degrade over a period of 3 to 4 months at 37°C.

To investigate host cell recruitment within various LiNx formulations loaded with C10, D6, or F5 containing ovalbumin (OVA)-encoding mRNA, or PBS, respectively, we subcutaneously injected the LiNx into the right flank of C57BL/6 mice. The composites were harvested at 3 and 7 days post-injection, and the number of viable cells within the composites was quantified using Acridine Orange/Propidium Iodide (AO/PI) staining. As depicted in **Fig. 1b**, a substantial number of host cells were recruited inside the scaffold for all groups at both time points. Notably, the incorporation of D6 and F5 mRNA LNPs into the composites markedly increased the number of recruited cells. In comparison to PBS LiNx, there was a 4.2-fold increase for D6-mRNA LiNx and a 2.9-fold increase for F5-mRNA LiNx in the number of cells recruited inside the composites on day 3. By day 7, there was a 2.9-fold increase for D6-mRNA LiNx and a 2.7-fold increase for F5-mRNA LiNx in the number of cells recruited inside the composites relative to the PBS LiNx group. Furthermore, a notable presence of APCs, including macrophage-like cells (CD3^-^CD11b^+^Ly6g^-^ CD11c^-^ cells) and DC-like cells (DCs; CD3^-^CD11b^+^Ly6g^-^CD11c^+^ cells), was identified within the composites on day 3 by flow cytometry analysis. Notably, the D6-mRNA LiNx group exhibited the highest level of recruitment of macrophage-like cells, which were 4.4-fold, 8.13-fold, and 2.0-fold higher than those observed in the PBS, C10, and F5-mRNA LiNx groups, respectively (**Fig. 1c, Supplementary Fig. 4**).

For an efficient vaccine, the expression profile of the antigen, along with adequate recruitment of host cells, plays a crucial role in eliciting a robust immune response.^24^ To assess the *in vivo* delivery efficacy of C10-, D6-, and F5-mRNA LiNx, we conducted *s.c.* injections of LiNx loaded with Cre-recombinase mRNA (mCre) in genetically engineered tdTomato (tdTom) reporter mice (Ai9 mice).^17^ These mice contain a LoxP-flanked stop cassette, preventing the expression of the tdTom protein until it is removed by Cre recombinase, allowing for the expression of tdTom. Our findings revealed that all three LiNx formulations resulted in notable levels of tdTom^+^ cells, including DC-like cells, macrophage-like cells, and CD11c^-^CD11b^-^ cells in LiNx on days 5 and 10 post-injection (**Fig. 1d–f, Supplementary Figs. 5–7**). In comparison to C10-and F5-mRNA LiNx, D6-mRNA LiNx exhibited a substantially higher level of transfected cells in the composite. On day 10, the number of transfected cells in D6 LiNx was approximately 129-fold and 117-fold higher than those for the C10 LiNx and F5 LiNx groups, respectively. The D6-mRNA LiNx group also showed approximately a 1,000-fold higher number of transfected DC-like cells compared to the other two groups. Additionally, the number of transfected macrophage-like cells was 2,125-fold and 1,691-fold higher in D6 LiNx compared to C10 LiNx and F5 LiNx, respectively. The biodistribution of LiNx formulations was further examined using Cy5-labeled mRNA, as shown in **Supplementary Fig. 8a**. All three mRNA LNPs were gradually released from the LiNx injection site over two weeks post-injection. Among the formulations, D6 exhibited the most delayed release profile, with approximately 25% of mRNA LNPs still retained within the LiNx platform. Additionally, at 24 hours post-vaccination, D6 LiNx showed that most LNPs remained at the injection site, with no detectable signal in other major organs (**Supplementary Fig. 8b-c**). The decrease in signal at the injection site observed initially was likely due to the uptake and degradation of Cy5-mRNA by local or recruited cells. In terms of transfection efficiency of D6 LiNx was further validated using mLuc mRNA. The data revealed that D6 LiNx transfection was largely confined to the injection site at 24 hours post-injection, with approximately 97% of the transfection signal detected there (**Fig. 1g-i**) and minimal transfection observed in other organs. Taken together, these results underscore the promise of the LiNx platform, especially D6 LiNx, in facilitating *in vivo* host immune cell recruitment and delivering antigen-encoding mRNA.

### mRNA LiNx effectively forms a local immunostimulatory niche

The cellular profiles of infiltrated cells within the composites were assessed at a later time point, specifically on day 14 post *s.c.* injection using mOVA LiNx. As depicted in **Fig. 2a**, even after 14 days, a substantially higher number of infiltrating host cells persisted in both D6 and F5 LiNx groups. In comparison to PBS LiNx (PBS+NHC) control and C10 LiNx, D6 LiNx and F5 LiNx showed approximately a 9-fold increase in terms of the number of infiltrated host cells within the composites. Moreover, a substantial increase in the presence of CD3^+^CD4^+^ T cells (**Fig. 2b, Supplementary Fig. 9**), CD3^+^CD8^+^ T cells (**Fig. 2c, Supplementary Fig. 10**), and CD3^-^CD11b^-^ Npk46^-^CD19^+^ B cells (**Fig. 2d, Supplementary Fig. 11**) were detected in the D6 LiNx compared to other groups, showing a 73-fold higher CD4^+^ T cell count, a 200-fold higher CD8^+^ T cell count, and a 305-fold higher B cell count for the D6 LiNx, compared to the PBS LiNx control. The proportion of specific immune cell types on day 14 was assessed for various treatment groups, as illustrated in **Fig. 2e (Supplementary Fig. 12)**. The predominant immunocytes within the composites for PBS LiNx control, C10 LiNx, and F5 LiNx were neutrophils, with approximately 60% of immunocytes within the composites were neutrophils for both PBS LiNx and C10 LiNx groups, and around 80% for the F5 LiNx. In contrast, the D6 LiNx contained substantial amounts of T cells and B cells: approximately 50% of the immunocytes within the LiNx were T cells, ∼20% CD8^+^ T cells, and ∼30% CD4^+^ T cells, while B cells constituted about 20% of the immunocytes. The immunostimulatory niche generated in D6 LiNx, characterized by a substantial influx of T cells and B cells, is highly advantageous for facilitating crosstalk among various immune cell types.

**Fig. 2.**
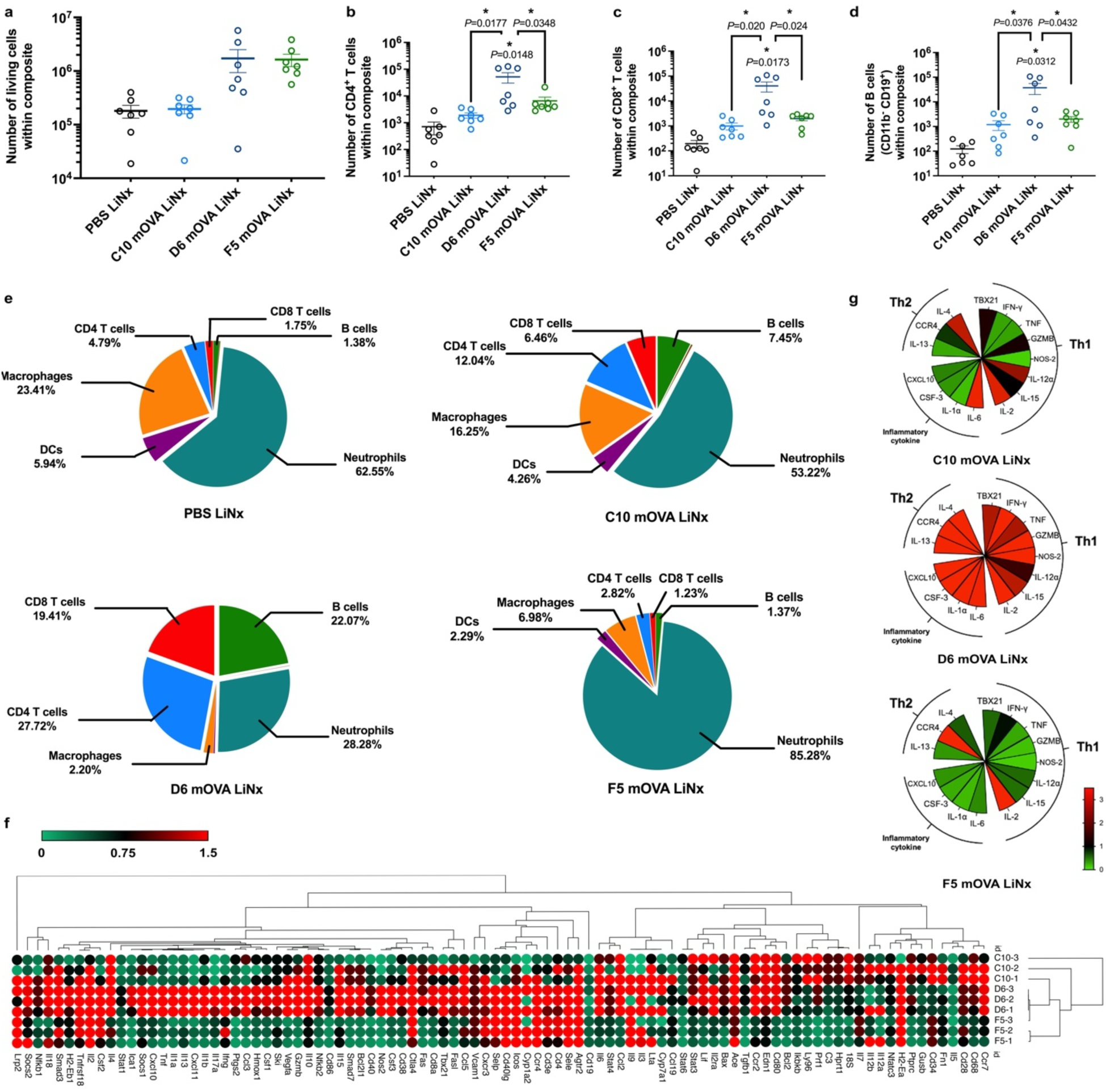
I*n vivo* assessments of local microenvironment generated by LiNx. **a,** Recruitment of host cells was assessed two weeks after *s.c.* administration of three different LNP/mRNA LiNx formulations in C57BL/6 mice (30 μg mOVA per mouse). The Cellaca MX High-throughput Automated Cell Counter was employed to count the living cells within the composites. **b–d,** On day 14, flow cytometry was utilized to determine the counts of CD3^+^CD4^+^ T cells **(b)**, CD3^+^CD8^+^ T cells **(c)**, and CD11b^-^CD19^+^ B cells **(d)** within the composite. **e,** Composition of host immune cells recruited within the composite was evaluated at two weeks after *s.c.* administering three different LiNx formulations in C57BL/6 mice (30 μg mOVA per mouse). This analysis included CD4 T cells, CD8 T cells, B cells, DC-like cells (CD11b^+^ CD11c^+^), macrophage-like cells (CD11b^+^CD11c^-^), neutrophils (CD11b^+^Ly6G^+^), and NK cells (CD11b^-^ NKp46^+^). **f–g,** RT-PCR array analysis was conducted using RNA isolated from the local microenvironment generated by C10, D6, and F5 LiNx formulations. C57BL/6 mice received the three different LiNx formulations loaded with mOVA through *s.c.* injection (30 μg mOVA per injection). The presented heatmap shows a panel of genes relevant to immune responses **(f)**. Subsequent RT-PCR analysis of selected genes from the panel confirmed that D6 LiNx promoted critical proinflammatory cytokine transcripts involved in the establishment of a local immunostimulatory niche. These included genes related to inflammatory cytokines and chemokines (IL-6, IL-1α, CSF-3, and CXCL10), Th1 response (TBX21, TNF, IFN-γ, GZMB, NOS-2, IL-12α, IL-15, and IL-2), and genes associated with Th2 response (IL-4, CCR-4, and IL-13). Data represent mean ± s.e.m. (n = 7 biologically independent samples for **a–e** and n = 3 biologically independent samples for **f–g**). Data were analysed using one-way ANOVA and Tukey’s multiple comparisons test for **a–d**. \**P* < 0.05.

The local microenvironments influenced by various LiNx formulations were further characterized using real-time polymerase chain reaction (PCR) arrays, analyzing RNA extracted from the local injection sites on day 14 post-injection. The mRNA levels related to immune responses were detected, quantified, and analysed (**Fig. 2f–g**). The findings suggest that, in contrast to other groups, the local injection site for D6 mOVA LiNx displayed increased expression of genes associated with inflammatory cytokines and chemokines, including IL-6, IL-1α, CSF-3, and CXCL10. Moreover, there were elevated levels of genes linked to Th1 immune responses, such as TBX21, TNF, IFN-γ, GZMB, NOS-2, IL-12α, IL-15, and IL-2, as well as genes associated with Th2 immune responses, namely IL-4, CCR-4, and IL-13 (**Supplementary Figs. 13–15**).

The above results reveal that, at 14 days post-injection, substantial host immune cell recruitment events were observed in the LiNx groups, particularly the D6 LiNx. The majority of these cells were associated with adaptive immune responses, specifically T cells and B cells, rather than neutrophils. The close proximity of these immune cells may enable crosstalk among them, potentially advantageous for the induction of a robust immune response. Moreover, the D6 LiNx group exhibited the formation of an immunostimulatory microenvironment characterized by elevated levels of immunostimulatory cytokines, transcripts related to both Th1 and Th2 responses, as well as an engaged innate inflammatory response.

### LiNx induces a potent antigen-specific immune response

The vaccination outcomes of the three LiNx formulations were assessed in C57BL/6 mice following *s.c.* injections with a single dosage consisting of 30 μg OVA mRNA, with PBS LiNx serving as the control. The LiNx-induced antigen-specific CD8^+^ T cell response was initially examined by collecting and analyzing cells from both LiNx and draining lymph nodes (dLNs) of the treated mice on day 14. As depicted in **Fig. 3a–b** and **Supplementary Figs. 16-17**, the D6 LiNx demonstrated a substantially higher level of antigen-specific CD8^+^ T cells compared to the C10 and F5 LiNx groups, with ∼64-fold and ∼11-fold increases detected inside the niches, respectively. Moreover, within the dLNs, the D6 LiNx exhibited a 3.7-fold and 2.9-fold higher number of antigen-specific CD8^+^ T cells, compared to the C10 and F5 LiNx groups, respectively. In contrast, the LiNx loaded with empty D6 LNPs did not generate a detectable immune response (**Supplementary Fig. 18**).

**Fig. 3.**
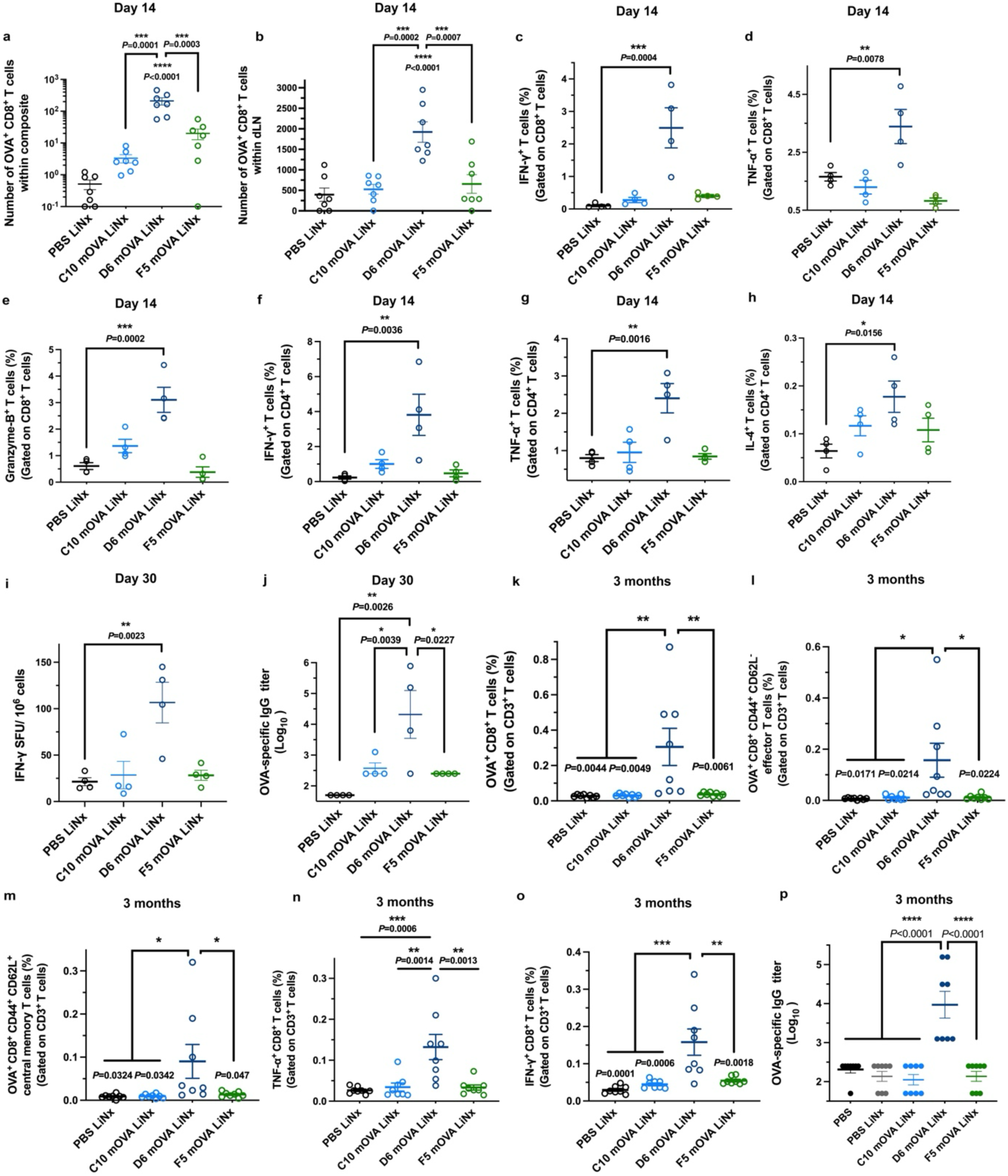
I*n vivo* assessments of antigen-specific immune activation by three different mRNA LiNx formulations. **a–b,** C57BL/6 mice were administered three different LNP/mRNA LiNx formulations loaded with mOVA via *s.c.* injection (30 μg per mouse). The number of OVA-specific CD8 T cells within the LiNx **(a)** and dLNs **(b)** was analysed by flow cytometry at two weeks following a single dosage of LiNx immunization. **c–h,** C57BL/6 mice were administered three different LNP/mRNA LiNx formulations loaded with mOVA via *s.c.* injection (30 μg per mouse). Mice were sacrificed two weeks after the injection, and their splenocytes were isolated and restimulated *in vitro* with OVA and SIINFEKL peptides (100 μg mL^-1^ OVA and 2 μg mL^−1^ SIINFEKL) for 12 h and assessed via flow cytometry and intracellular cytokine staining to determine the percentages of CD8^+^IFN-γ^+^ cells **(c)**, CD8^+^TNFα^+^ cells **(d)**, CD8^+^ Granzyme B^+^ cells **(e)**, CD4^+^IFN-γ^+^ cells **(f)**, CD4^+^TNFα^+^ cells **(g)**, and CD4^+^IL-4^+^ cells **(h)**. **i**, Frequency of IFN-γ-producing cells among restimulated splenocytes isolated from vaccinated mice on day 30 post-vaccination, assessed via ELISPOT. **j**, Titer of OVA-specific IgG antibodies in blood serum on day 30, determined by ELISA. **k-o,** C57BL/6 mice were administered with three different LNP/mRNA LiNx formulations loaded with mOVA via *s.c.* injection (30 μg per mouse). Mice were sacrificed three months after the injection, and their splenocytes were isolated for analysis. The percentages of OVA-specific CD8 T cells (CD3^+^ CD8^+^ OVA^+^ cells) **(k)**, OVA-specific effector CD8 T cells (CD3^+^CD8^+^OVA^+^CD44^+^CD62L^-^ cells) **(l),** and OVA-specific central memory CD8 T cells (CD3^+^CD8^+^OVA^+^CD44^+^CD62L^+^ cells) **(m)** were determined. Splenocytes were restimulated *in vitro* with OVA and SIINFEKL peptide (100 μg mL^-1^ OVA and 2 μg mL^−1^ SIINFEKL) for 12 h and assessed via flow cytometry and intracellular cytokine staining to determine the percentages of CD3^+^CD8^+^TNFα^+^ cells **(n)** and CD3^+^CD8^+^IFN-γ^+^ cells **(o)**. **(p)** Titers of OVA-specific IgG antibodies in blood serum on day 90, determined by ELISA. Data represent the mean ± s.e.m. (n = 7 **(a–b)** biologically independent samples, n = 4 **(c–j)** biologically independent samples, n = 8 **(k–p)** biologically independent samples). Data were analysed using one-way ANOVA and Tukey’s multiple comparisons test for **a–o**. \**P <* 0.05, \*\**P <* 0.01, \*\*\**P <* 0.001, \*\*\*\**P <* 0.0001.

The spleens from vaccinated mice were harvested on day 14, dissociated into a cell suspension, and subjected to *ex vivo* antigen restimulation. The D6 LiNx yielded increased frequencies of CD8^+^IFN-γ^+^, CD8^+^TNFα^+^, and CD8^+^Granzyme B^+^ cell populations (**Fig. 3c–e, Supplementary Fig. 19**). Specifically, compared to the C10 and F5 LiNx-treated groups, the D6 LiNx exhibited approximately 9.0-and 6.2-fold increases in CD8^+^IFN-γ^+^ cell frequencies, respectively. Alongside the robust CD8^+^ T cell response, the D6 LiNx also showed higher numbers of Th1 cells (CD4^+^IFN-γ^+^ and CD4^+^TNFα^+^), as depicted in **Fig. 3f–g**. On day 14 post-vaccination, there was a 16.4-fold higher level of antigen-specific CD4^+^IFN-γ^+^ Th1 cells and a 3.0-fold higher level of antigen-specific CD4^+^TNFα^+^ Th1 cells for the D6 LiNx group. In contrast, no significant increases were observed in the C10 and F5 LiNx-treated groups. Furthermore, the frequency of antigen-specific CD4+IL-4+ Τh2 cells was examined for all groups (**Fig. 3h**). In comparison to the PBS LiNx control, the D6 LiNx group exhibited a higher number of antigen-specific Th2 cells (2.8-fold increase, *P* = 0.0156), in contrast to the C10 and F5 LiNx groups (1.8- and 1.7-fold higher, respectively; *P* > 0.5).

The antigen-specific response after vaccination was further evaluated on day 30 post-vaccination. As depicted in **Fig. 3i**, the splenocytes from vaccinated mice collected on day 30 were subjected to *in vitro* antigen restimulation. The D6 LiNx induced a substantially higher level of antigen-specific T-cell response (**Supplementary Fig. 20**). Approximately 3.7-fold and 3.8-fold higher levels of IFN-γ-secreting cells were detected within the D6 LiNx group, compared with C10 LiNx and F5 LiNx, respectively. In addition, as shown in **Fig. 3j**, D6 LiNx induced a higher OVA-specific IgG titer including both IgG1 and IgG2c subclass titers, indicating a potent humoral response (**Supplementary Fig. 21**) as well. The antibody responses generated by C10 and F5 LiNx groups were limited.

The memory immune responses of LiNx were evaluated at three months post-vaccination. Substantial levels of antigen-specific CD8^+^ T cells were still present in the spleen of mice in the D6 LiNx group (**Fig. 3k, Supplementary Figs. 22 and 23**). More importantly, within these T cells, there were both CD44^+^CD62L^-^ effector T cells and higher levels of CD44^+^CD62L^+^ central memory T cells. Specifically, the D6 LiNx group exhibited approximately a 23.8-fold higher level of antigen-specific effector T cells and a 10.1-fold higher level of central memory T cells, underscoring the establishment of long-term memory responses (**Fig. 3l–m**). Splenocytes harvested from vaccinated mice on day 90 were also subjected to *ex vivo* antigen restimulation. As depicted in **Fig. 3n–o**, the D6 LiNx group exhibited 5.0-fold higher CD8^+^TNFα^+^ cells than the PBS LiNx control (*P* = 0.0006) and 5.3-fold higher CD8^+^IFN-γ^+^ cells than the PBS LiNx control (*P* = 0.0001) (**Supplementary Figs. 24–26**). In addition to T cell responses, a substantially higher OVA-specific IgG titer, including both IgG1 and IgG2c subclass titers, was observed in the D6 LiNx-treated group at 90 days post-vaccination, indicating a potent long-lasting humoral response (**Fig. 3p**, **Supplementary Fig. 27**). Furthermore, an examination of the biosafety profiles for C10, F5, and D6 LiNx showed no significant differences in body weight across all LiNx-treated groups throughout the vaccination schedule, and no significant changes in serum cytokine levels (IFN-γ, TNF-α, IL-4, IL-10, and IL-2) post-vaccination in the D6 LiNx treated mice (**Supplementary Fig. 28 and 29**).

The above results indicate that, following a single-dose vaccination, D6 LiNx effectively elicited more potent antigen-specific Th1 and Th2 responses than C10 and F5 LiNx. Moreover, D6 LiNx vaccination resulted in a strong and persistent memory immune response. In contrast, no memory immune responses were observed three months after vaccination in the C10 and F5 LiNx groups.

### Anti-tumour effects induced by a single-dose vaccination of D6 mRNA LiNx

Given the potent antigen-specific immune responses induced by the D6 LiNx, we investigated its efficacy as a cancer vaccine in therapeutic and prophylactic tumour models. In the first study, C57BL/6 mice were *s.c.* inoculated with 3 × 10^5^ OVA-expressing MC38 colorectal cancer cells on the right posterior side on day 0. On day 4, the mice received vaccinations with C10, D6, or F5 mRNA LiNx, each containing 30 μg of mOVA (**Fig. 4a**). Control groups included the NHC and OVA protein (30 μg of OVA per mouse) mixed with Alhydrogel^®^ or OVA in NHC. Additionally, C57BL/6 mice were immunized with 3 doses of D6 LNPs containing 10 μg of mOVA on days 4, 11, and 18, as a control. As illustrated in **Fig. 4b–d**, the single dose D6 LiNx exhibited effective tumour suppression in this treatment model, resulting in a median survival time of 75 days compared to 31 days for the negative control group (**Supplementary Fig. 30**). In the D6 LiNx group, 50% of the mice remained tumour-free beyond 100 days. Mice treated with the NHC, OVA in NHC, OVA in Alhydrogel^®^, C10 LiNx, and F5 LiNx exhibited limited levels of therapeutic efficacy, leading to low extension of median survival time. In comparison to the group treated with the standard three doses D6 mRNA LNP immunization, which resulted in a median survival time of 37.5 days, a single dose of D6 LiNx produced a markedly improved antitumour effect, demonstrating the capability of LiNx in enhancing the anti-tumour effect of mRNA LNPs.

**Fig. 4.**
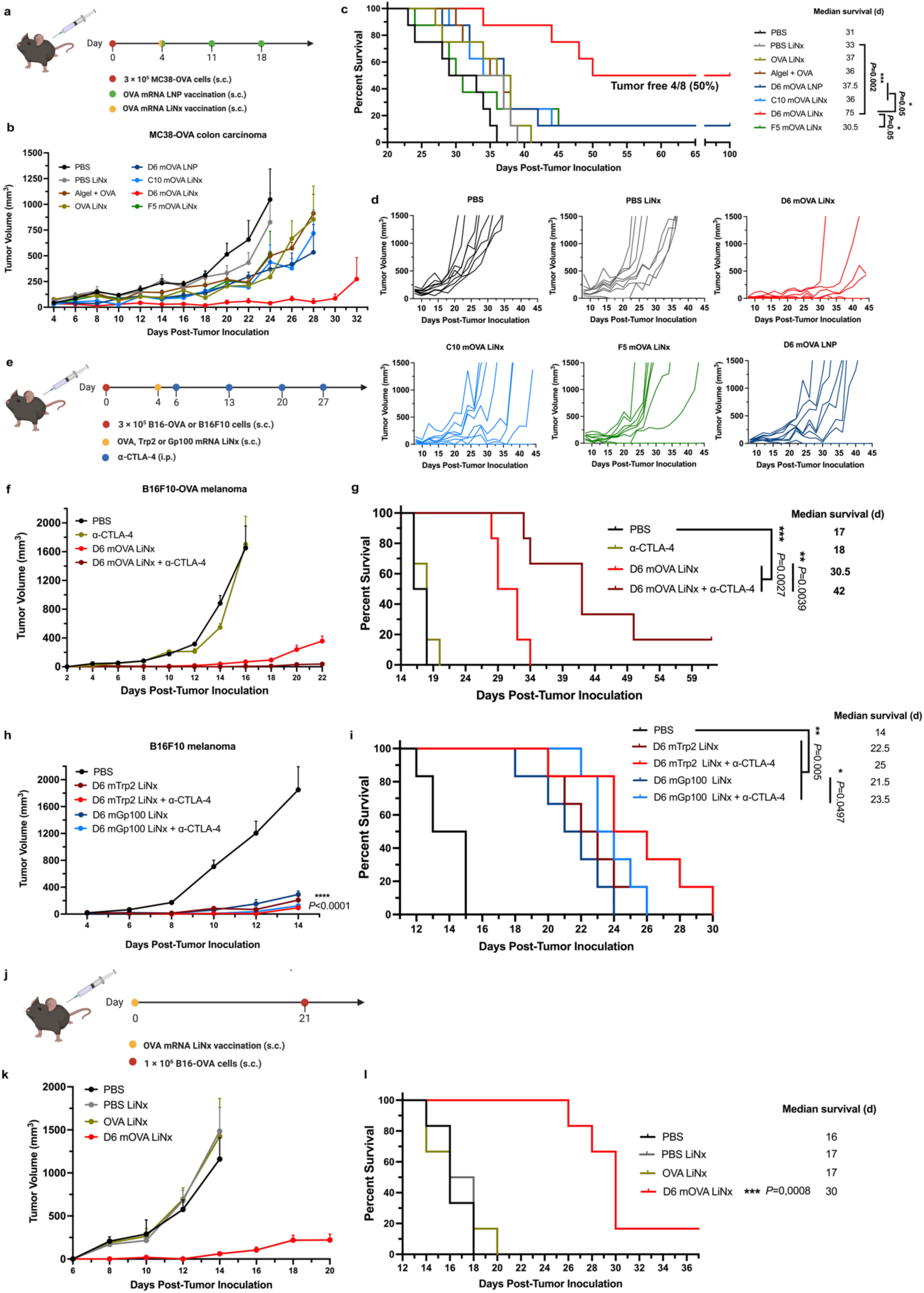
Anti-tumour efficacy of the top LiNx formulations as therapeutic and prophylactic vaccines. **a–d,** Schematic and results of a syngeneic therapeutic vaccination model for MC38-OVA in C57BL/6 mice. Mice were inoculated *s.c.* with MC38-OVA and then given three *s.c.* injections, one week apart, of mOVA-loaded D6 LNPs (10 μg mOVA per injection) or PBS. For LiNx treatment group, mice were administered three different LiNx formulations loaded with mOVA LNPs (30 μg per mouse) or OVA protein (10 μg protein per mouse) via a single *s.c.* injection. OVA protein mixed with Alhydrogel® (1:1) (10 μg protein per mouse) served as a control group. Average tumour volume **(b)**, survival curves **(c)**, and individual tumour volume **(d)** are shown. **e–i,** Schematic and results of a therapeutic vaccine against another syngeneic model, B16F10 melanoma in C57BL/6 mice, using OVA model antigen **(f–g)** and melanoma-associated antigens **(h-i)**. Mice were inoculated *s.c.* with B16-OVA **(f–g)** or B16F10 cells **(h–i)** and then *s.c.* administered with D6 LiNx loaded with mOVA, mTrp2 or mGp100 (30 μg per mouse). Three groups received the anti-CTLA-4 mAb (100 μg per *i.p.* injection) treatment in combination with LiNx treatment. Average tumour volume **(f, h),** and survival curves **(g, i)** are shown. **j–l,** Schematic and results of a prophylactic vaccine against melanoma in C57BL/6 mice using OVA model antigen. Mice were *s.c.* administered with a single dose of D6 LiNx loaded with mOVA (30 μg per mouse) and then inoculated *s.c.* with B16-OVA cells on day 21. Average tumour volume **(k),** and survival curves **(l)** are shown. Data represent mean ± s.e.m. with n = 8 (**c**–**d**), n = 6 (**f–i**, **k**–**l**) biologically independent samples. Survival curves were compared using log-rank Mantel–Cox test, and the stack of *P* values were corrected by Holm-Šídák method for multiple comparisons with α set to 0.05. \**P <* 0.05, \*\**P <* 0.01, \*\*\**P <* 0.001; NS, not significant; *i.p.* intraperitoneal; αCTLA-4, anti-CTLA-4 mAb.

We further evaluated the therapeutic efficacy of D6 LiNx in the B16-OVA melanoma tumour model, utilizing the model OVA antigen. C57BL/6 mice were *s.c.* inoculated on the right posterior side with 3 × 10^5^ B16-OVA cells on day 0. On day 4, with an average tumour size of ∼25 mm³, mice were vaccinated with D6 LiNx containing 30 μg mOVA (**Fig. 4e**). As depicted in **Fig. 4f–g**, D6 LiNx demonstrated a stronger tumour suppression effect in this treatment model, resulting in a median survival time of 30.5 days compared to 17 days for the negative control (**Supplementary Fig. 31**). As shown in **Fig. 2f**, the local microenvironment at the injection site in D6 LiNx-treated group exhibited a highly elevated CTLA-4 marker, which prompted us to combine anti-CTLA-4 therapy with the LiNx vaccination to evaluate their combined effect. When D6 LiNx was administered in conjunction with an immune checkpoint inhibitor (100 μg anti-CTLA-4 monoclonal antibody, given *i.p.* on days 6, 13, 20, and 27), a synergistic effect was observed, extending the median survival time to 42 days. In contrast, no significant tumour suppression effect was observed in the group treated solely with anti-CTLA-4 antibody, as compared to the PBS LiNx control.

Subsequently, the D6 LiNx was evaluated in the syngeneic melanoma mouse model using two clinically relevant tumour antigens, tyrosinase-related protein 2 (Trp2) and glycoprotein 100 (Gp100) (**Fig. 4e**). C57BL/6 mice were *s.c.* inoculated on the right posterior side with 3 × 10^5^ B16F10 cells on day 0. On day 4, with an average tumour size of ∼25 mm³, mice were vaccinated with D6 LiNx containing 30 μg of mRNA encoding either Trp2 or Gp100. The potent anti-tumour effect was observed with these two antigens, resulting in prolonged median survival times of 22.5 and 21.5 days for D6 mTrp2 LiNx and D6 mGp100 LiNx, respectively (**Fig. 4h–i, Supplementary Fig. 32**). However, no significant improvement was observed when combining these two D6 LiNx formulations with anti-CTLA-4 antibody treatment.

Beyond the therapeutic cancer model, D6 LiNx was also assessed in a prophylactic B16-OVA melanoma model. C57BL/6 mice were immunized on day 0 with D6 LiNx containing 30 μg of mOVA. On day 21, animals were *s.c.* inoculated on the right posterior side with 1 × 10^6^ B16-OVA cells (**Fig. 4j**). A single injection of D6 LiNx yielded a stronger tumour inhibition efficacy with prolonged overall survival times compared to the PBS, PBS-NHC, and OVA-NHC groups. The median survival time was 30 days for the D6 LiNx group, compared to 16–17 days for the three control groups (**Fig. 4k–l, Supplementary Fig. 33**). In addition, LiNx loaded with empty D6 LNPs did not generate a significant anti-tumour effect (**Supplementary Fig. 34**).

To further confirm the efficacy of the D6 mOVA LiNx and assess its long-term protective effect, we conducted a tumour rechallenge study with an increased sample size (**Supplementary Fig. 35**). C57BL/6 mice were immunized as previously described and, on day 21, were subcutaneously inoculated with 3 × 10⁵ B16-OVA cells on the right posterior side. Survival was monitored over 100 days, and 17 of 18 mice (94.4%) remained tumour-free following the single D6 LiNx vaccination (**Supplementary Fig. 35b**). These mice were subsequently rechallenged with 3 × 10^5^ B16-OVA cells on the right posterior side. As shown in **Supplementary Fig. 35c**, 11 of 17 (64%) mice remained tumour-free post-rechallenge, demonstrating the long-term protective effect of D6 LiNx. These findings further confirm the antitumour efficacy of the D6 LiNx platform and highlight its potential for sustained protection.

### Modulation of the tumour microenvironment by LiNx

To assess the effect of LiNx treatment on the tumour microenvironment, we analysed the infiltrating immune cells in the tumour mass. As shown in **Fig. 5a**, C57BL/6 mice were inoculated with 3 × 10^5^ B16-OVA cells on day 0. Starting on day 4, mice received three subcutaneous (*s.c.*) injections of mOVA-loaded D6 LNPs (10 μg mOVA per injection) or PBS at one-week intervals (days 4, 11, and 18). For the LiNx treatment group, mice were administered the D6 LiNx formulations (30 μg per mouse) via a single *s.c.* injection on day 4. On day 21, the mice were euthanized, and tumour tissues were collected, homogenized, and analysed by flow cytometry. As shown in **Fig. 5b**, treatment with the D6 LiNx resulted in a substantial increase in CD8^+^ T cells within the tumour microenvironment. Approximately 4.5% of the total cell population consisted of CD8^+^ T cells, representing a three-fold increase compared to treatment with 3 doses of the D6 LNPs, which only induced about 1.5% CD8^+^ T cells in the tumour. In addition, a 3.9-fold higher (1.3%) OVA-specific CD8^+^ T cells among the total cell population in the tumour was observed in the D6 LiNx group, compared to the 3 doses of D6 LNPs (**Fig. 5c**). Similarly, higher levels of CD4^+^ T cells, NK cells, macrophages, and DCs were also observed in the D6 LiNx group (**Fig. 5d–g**). This indicates that the D6 LiNx induced a robust antigen-specific immune response and modulated the tumour microenvironment.

**Fig. 5.**
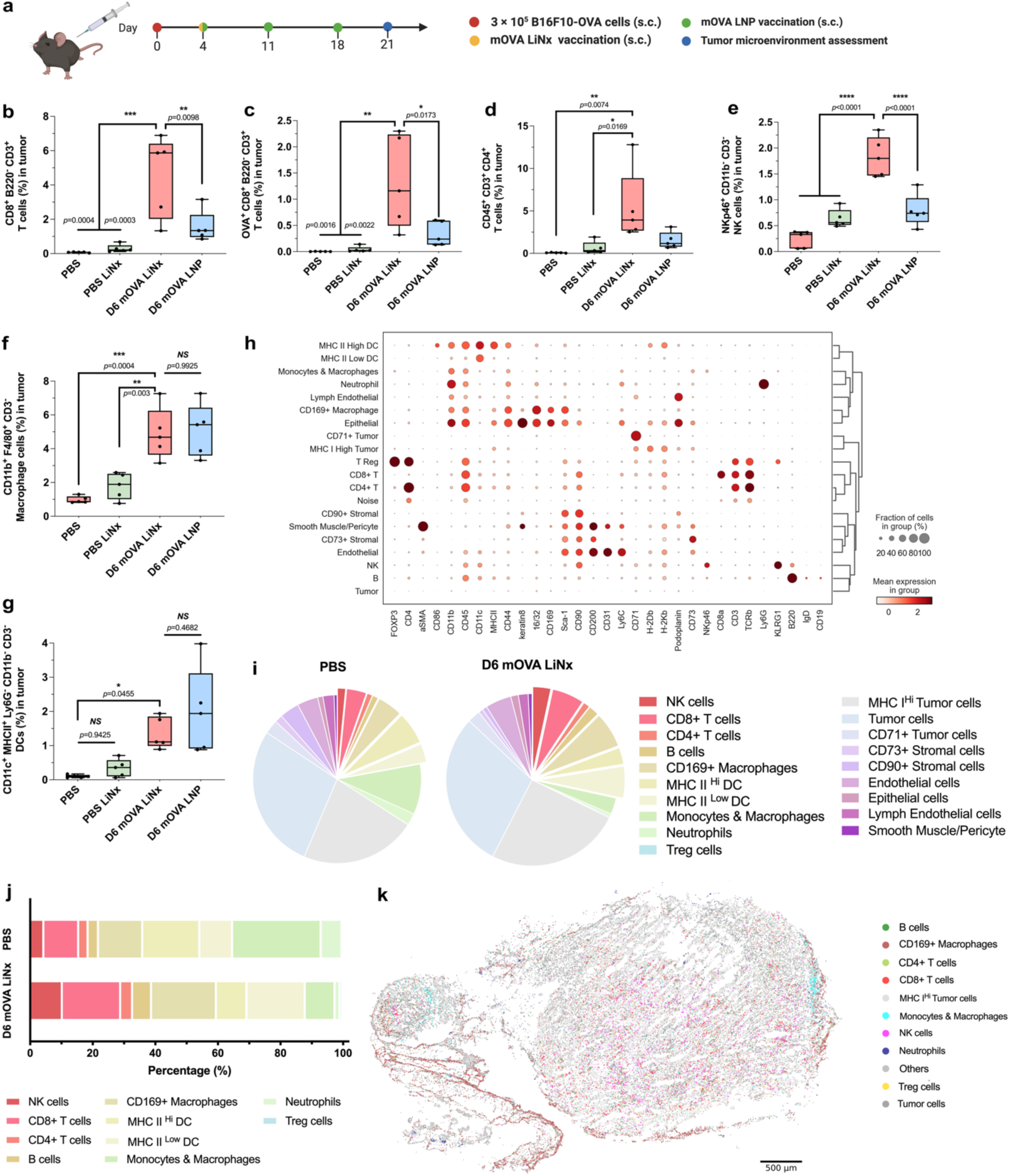
Modulation of tumour microenvironment of the D6 LiNx formulation. **a–g,** Schematic and analysis of the tumour microenvironment in a therapeutic vaccination model for B16-OVA in C57BL/6 mice. Mice were inoculated subcutaneously (*s.c.*) with B16-OVA and subsequently received three weekly *s.c.* injections of mOVA-loaded D6 LNPs (10 μg mOVA per injection) or PBS. For the LiNx treatment group, mice were administered with the D6 LiNx loaded with mOVA (30 μg per mouse) via a single *s.c.* injection on day 4. On day 21, tumours were collected and analysed by flow cytometry to determine the percentages of (**b**) CD3^+^CD8^+^B220^-^ T cells, (**c**) OVA-specific CD3^+^CD8^+^B220^-^ T cells, (**d**) CD45^+^CD3^+^CD4^+^ T cells, (**e**) NKp46^+^CD11b^-^CD3^-^ NK cells, (**f**) F4/80^+^CD11b^+^CD3^-^ macrophages, and (**g**) MHCII^+^CD11c^+^CD11b^-^Ly6G^-^CD3^-^ DCs. **h-k,** CODEX multiplex imaging analysis of tumour tissue collected on day 14 post B16-OVA inoculation. For the LiNx treatment group, mice were administered with the D6 LiNx formulation (30 μg mOVA per mouse) via a single *s.c.* injection on day 4. **(h)** Dotplot of cell types by marker expression derived from CODEX multiplexed imaging for the pooled D6 LiNx (n = 2) and PBS-treated tumours (n = 3) using unsupervised clustering with the percentage of cells with a marker Z-score above 0.7. **(i)** Percentage distribution of immune and tumour cell types within the D6 LiNx (n = 2) and PBS-treated tumours (n = 3) as quantified by CODEX multiplexed imaging. **(j)** Comparison of the normalized percentage of immune cell subsets within the D6 LiNx-treated tumours (n = 2) and PBS-treated tumours (n = 3). **(k)** Cell type map of the representative D6 LiNx tumour derived from CODEX multiplexed imaging with colored data points representing selected immune and tumour cell types with other cell types lumped together in the same gray color (scale bar = 500 μm).

To further validate the effect of the LiNx platform on the tumour microenvironment, we employed CODEX multiplexed fluorescence microscopy for spatial proteomics that uses iterative imaging and DNA-barcoded antibodies to enable the simultaneous imaging of multiple markers.^25–28^ We designed a panel to identify major adaptive and innate immune cell types (**Fig. 5h, Supplementary Tables 2–4**). C57BL/6 mice were inoculated with 3 × 10^5^ B16-OVA cells on day 0 and were treated with the D6 LiNx (30 μg mOVA per mouse) via a single *s.c.* injection on day 4. On day 14 post-tumour inoculation, tumour tissues were collected, sectioned, and analysed. As shown in **Figs. 5i–k** and **Supplementary Figs. 36–37**, the LiNx group recruited substantially higher levels of adaptive immune cells, including CD8^+^ and CD4^+^ T cells, as well as B cells, infiltrating the tumour region compared to the PBS group. Additionally, tumours collected from mice treated with the D6 LiNx showed a substantial increase in NK cells and CD169^+^ macrophages. In contrast, neutrophils and non-CD169^+^ macrophages & monocytes were among the majority of immune cells in the tumour region isolated from the PBS group. The observed elevation of CD169^+^ macrophages indicates a shift in the tumour microenvironment induced by the D6 LiNx immunization. Notably, recent studies have shown that a high density of CD169^+^ macrophages is associated with prolonged survival and favourable clinical outcomes in patients with tumours.^29–35^

Taken together, our results reveal that treatment with the D6 LiNx altered the tumour microenvironment by increasing the infiltration of T and B cells, which are involved in the adaptive immune response, and a notable elevation in NK cells and CD169^+^ macrophages, which play a critical role in the antitumour process. These results highlight the potency of the D6 LiNx platform as a newly designed immunostimulatory vaccine platform.

### LiNx amplifies the immune response of mRNA LNPs

To further differentiate the immune responses triggered by D6 mRNA LNPs and LiNx, we analysed the expression of 547 genes in splenocytes from vaccinated mice using the nCounter Analysis System (NanoString Technology). In detail, the D6 LiNx comprising 30 μg mOVA was *s.c.* injected on day 0, compared with three doses of D6 LNPs containing 10 μg mOVA each on days 0, 7, and 14. On day 21, spleens from vaccinated mice were collected, dissociated into single-cell suspension, and then subjected to *in vitro* antigen restimulation. After 24 h, splenocytes were collected, and the RNA was extracted and examined using NanoString analysis. As depicted in **Fig. 6a–b**, the comparison between untreated mice and the D6 LiNx group revealed 376 distinctive genes, including 25 downregulated and 351 upregulated genes. When comparing the single-dose D6 LiNx with the three-dosage D6 LNPs treatment, 344 genes exhibited significant differences, consisting of 35 downregulated and 309 upregulated. Upon closer examination of various subsets of related genes, including Th1, Th2, Th17, and Treg-related genes,^36^ as well as B cell receptor signaling genes, the D6 LiNx-treated group exhibited upregulation in 6 out of 15 Th1-related genes, 9 out of 16 Th2-related genes, 13 out of 19 Th17-related genes, and 12 out of 30 B cell signaling-related genes, in comparison with a 3-dose free-D6 LNP control group (**Fig. 6c–e**). Among Treg cell-related genes, there were no significant differences, except for the downregulation of 1 out of 7 genes in the D6 LiNx group. By employing Database for Annotation, Visualization, and Integrated Discovery (DAVID) functional annotations for Kyoto Encyclopedia of Genes and Genomes (KEGG) pathways to establish connections between the information and higher-order functional aspects, we discovered that the D6 LiNx platform augmented immune responses through multiple pathways (**Fig. 6f**). In addition to its enhancement effect on Th1 and Th2 differentiation, antigen presentation, and B cell/T cell receptor signaling pathways, a noteworthy revelation emerged: Th17 differentiation and the IL-17 signaling pathway exhibited marked enhancement following D6 LiNx treatment. This unexpected finding suggests that the increased effectiveness of D6 LiNx in anticancer efficacy, as compared to free D6 LNPs, may be attributed to the substantial enhancement of Th17 cell-mediated anti-tumour immunity, which has been recognized for its considerable therapeutic potential (**Fig. 6g**).^37–44^ To validate the functionality of the Th17 immune response induced by the D6 LiNx, we performed IL-17 depletion experiments using a prophylactic vaccination model for OVA-expressing melanoma in C57BL/6 mice. The mice received either a single *s.c.* injection of the D6 LiNx (30 μg mOVA per injection) or three *s.c.* injections of mOVA-loaded D6 LNPs (10 μg mOVA per injection) administered one week apart, followed by *s.c.* inoculation with B16-OVA cells. IL-17-depleting antibodies were administered *i.p.* every three days (200 μg per mouse). The results demonstrated that IL-17 depletion led to approximately a 40% reduction in survival rate in the D6 LiNx treatment group (**Fig. 6h**). In contrast, IL-17 depletion had no significant effect on the survival rate of the traditional 3-dose D6 mRNA LNP treatment group, underscoring the critical role of the Th17 immune response in the observed anti-tumour effect of the D6 LiNx formulation. Moreover, no significant changes in lymphocyte composition were detected in the blood of α-IL-17-treated C57BL/6 control mice, indicating that antibody treatment does not substantially alter the immune system (**Supplementary Fig. 38**).

**Fig. 6.**
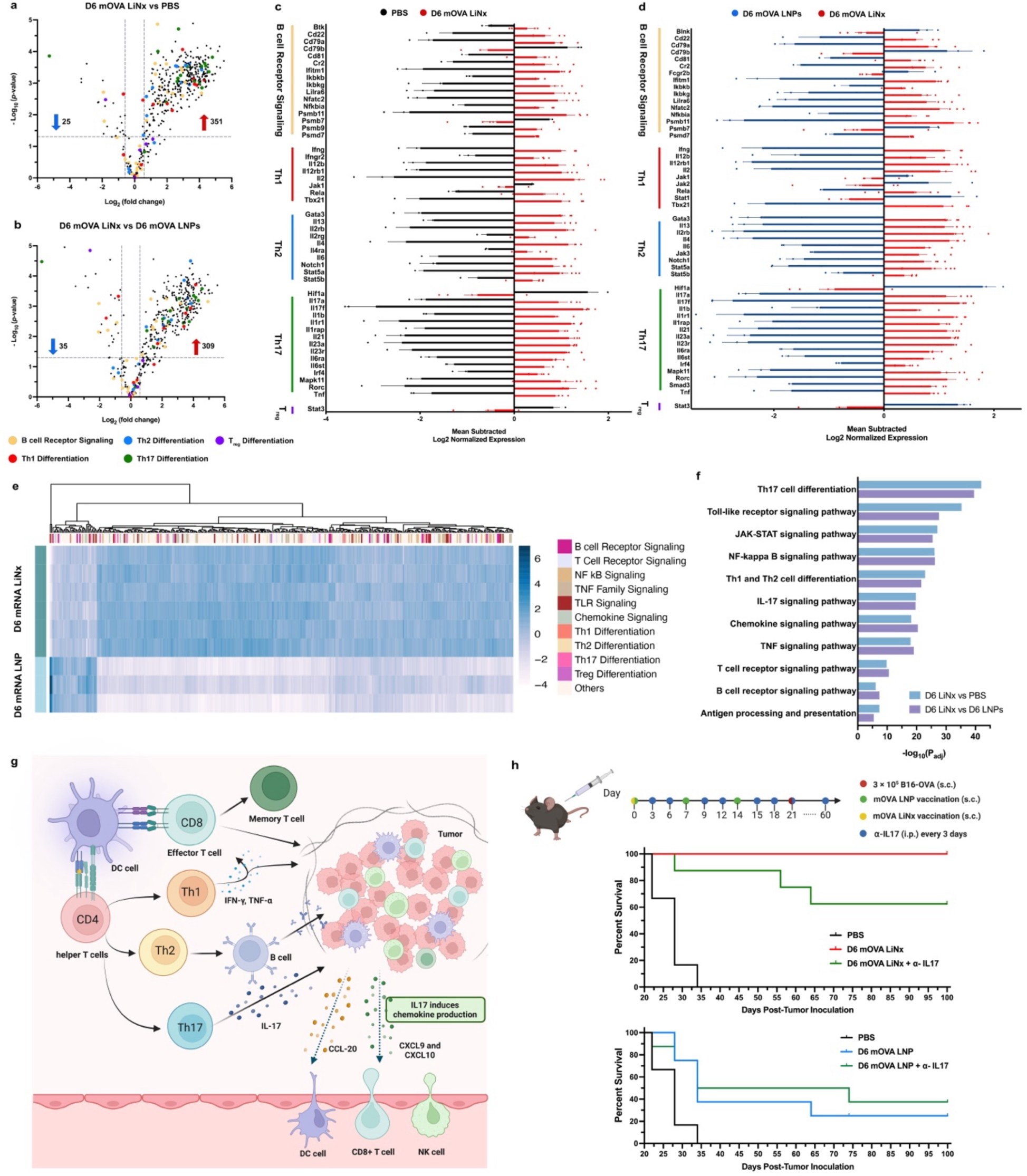
Characterization of immune activation profile generated by the D6 LiNx. C57BL/6 mice were given three *s.c.* injections, one week apart, of mOVA-loaded D6 (10 μg mOVA per injection) or PBS. For LiNx treatment group, mice were administered D6 LiNx loaded with mOVA (30 μg per mouse) on day 0. On day 21 post-vaccination, the splenocytes were restimulated *in vitro* with OVA and SIINFEKL peptide (100 μg mL^-1^ OVA and 2 μg mL^−1^ SIINFEKL) for 12 h and the RNA was extracted and assessed using a NanoString nCounter Analysis System. **a–b,** Volcano plot illustrates the differentially expressed genes between the D6 LiNx and PBS groups **(a)**, as well as between the D6 LiNx and D6 LNP groups **(b)**. *P* < 0.05; two-sided unpaired limma-moderated t-test; absolute fold change ≥ 1.5. **c–d,** Selected genes related to B cell signaling, Th1, Th2, Th17, and Treg differentiation are shown. **e,** Heatmap presenting a panel of genes relevant to immune responses for D6 LNP group and D6 LiNx group. **f,** Bar plots showing differential pathways enriched after vaccination of a single-dose D6 LiNx compared with PBS or three doses of D6 LNPs using Database for Annotation, Visualization and Integrated Discovery (DAVID) functional annotations for Kyoto Encyclopedia of Genes and Genomes (KEGG) pathways. Data represent mean ± s.e.m. **(d–e)**. Experiments were conducted with n = 6 biologically independent samples for the D6 LiNx group and n = 3 biologically independent samples for PBS LiNx and D6 LNP groups. **g**, Schematic of the immunostimulatory niche generated by LiNx through effective recruitment of host immune cells to the microgel matrix, leading to antigen presentation to CD8^+^ and CD4^+^ T cells (including Th1, Th2, and Th17 cells). Th17 cells facilitate the migration and persistence of effector T cells and NK cells within the tumour microenvironment by promoting the secretion of chemokines CXCL9 and CXCL10 by primary tumour cells. Moreover, Th17 cells stimulate the release of CCL20 from tumour cells, thereby attracting CCR6^+^DCs. **h**, Schematic and results of IL-17 depletion experiments in the prophylactic vaccination model for OVA-expressing melanoma in C57BL/6 mice. Mice were given one *s.c.* injection of D6 LiNx (30 μg mOVA per injection) or three *s.c.* injections, 1 week apart, of mOVA-loaded D6 LNPs (10 μg mOVA per injection) before *s.c.* inoculation of B16-OVA cells (3*10^5^ cells), and antibody for IL-17 depletion were injected every 3 d (*i.p.*, 200 μg per mouse). Survival curves over time are shown (n = 6 biologically independent mice for the PBS group and 8 biologically independent mice for other groups).

## DISCUSSION

Previous reports on LNP-mediated gene delivery and mRNA vaccination revealed that both the choice of lipids^45^ and their molar ratios significantly impact various aspects,^22^ including the encapsulation efficiency^46^ of nucleic acid payloads^47^, transfection efficiency,^22^ targeting profiles within cells and tissues,^48^ as well as the immune response profile.^16^ Despite enormous potential, the spatial and temporal control over the immune response still persists in LNP-based engineering.^49^ In this study, we tested a new concept of LiNx, an mRNA LNP-loaded microgel matrix, which promotes endogenous immune cell recruitment and retention, and creates a programmable immunostimulating microenvironment, therefore maximizing antigen expression and presentation locally. This configuration yields a more robust immune response with a single-dose vaccination in mouse colon carcinoma and melanoma models.

Here, we employed a previously developed NHC due to its ability to persistently recruit and retain host immune cells following *s.c.* or *i.m.* injection. This is crucial for facilitating effective crosstalk between relevant immune cell types. We tested three different LNPs with potent transfection capabilities of BMDCs in the LiNx context. Notably, the D6 LiNx, among the three, establishes the most effective local niche at week 2, characterized by a high abundance of both T cells and B cells, in contrast to the predominant neutrophil or macrophage composition observed in the C10 and F5 LiNx groups. The formation of a lymphoid tissue-like niche provides an immunostimulatory environment favourable for the induction of adaptive immune response. Interestingly, substantial differences were observed regarding host cell recruitment, transfection efficiency, local microenvironment, and anti-cancer immune responses among the three LiNx formulations. These variations may be attributed to differences in LNP stability, transfection profiles, and/or immunostimulatory cues provided by the LNPs themselves. In future applications of the LiNx platform, it is imperative to screen combinations of different LNPs and NHC. Additionally, physical properties such as the stiffness and biodegradability of NHC may also influence the transfection and immune response outcomes. Therefore, optimizing NHC for enhanced immune responses should involve a comprehensive examination of these physical attributes.

When we analysed the tumour microenvironment following treatment with the D6 LiNx platform using flow cytometry and CODEX multiplex imaging, we observed a marked increase in adaptive immune cells, particularly antigen-specific T cells, infiltrating the tumour. Notably, the D6 LiNx treatment group also exhibited a substantial presence of NK cells and CD169^+^ macrophages. The elevated NK cell levels are particularly striking, given their critical role in the innate immune response and the ability to directly target and eliminate tumour cells. Similarly, CD169+ macrophages, known for their role in antigen presentation, may facilitate T cell activation within the tumour microenvironment.^29–35^ These findings highlight the enhanced potency of the D6 LiNx platform, which strongly correlates with robust activation of both adaptive and innate immune responses. This dual activation is likely to contribute to a more sustained and effective immune response against tumours. Collectively, our results underscore the potential of the D6 LiNx platform as a highly effective immunostimulatory vaccine platform.

When comparing a single dose of D6 LiNx to three doses of conventional D6 LNPs, we observed enhanced immune responses and therapeutic efficacy with D6 LiNx. This phenomenon may be attributed to the initial successful transfection of host cells by D6 LNPs in the LiNx, leading to continuous host cell recruitment and effective crosstalk among recruited cells in LiNx, creating a local environment conducive to the potentiation of the immune response. More interestingly, our analysis revealed that the LiNx elicited a more robust Th17 response, in addition to Th1 and Th2 responses, suggesting the critical role of Th17 cells in generating potent antitumour efficacy in this context (**Fig. 5g**).

In summary, we present an effective approach to potentiate mRNA LNP cancer vaccine by creating an immune cell recruiting and retention microgel matrix as a local immunostimulating niche. The LiNx matrix does not rely on lymph node trafficking of LNPs, but rather leverages its capability to locally enhance antigen expression and presentation, as well as immune cell crosstalk, resulting in potent antigen-specific responses. Among the three tested LiNx formulations, D6 LiNx exhibited continuous host cell recruitment and differentiation, as well as Th1, Th2, and Th17 responses that correlate with the most potent antitumour efficacy among the three LiNx formulations and when compared to the traditional three-dose regimen for D6 LNP vaccination. The results showcase a potent strategy to augment immune responses elicited by mRNA LNPs through a biomaterials-enabled lymphoid niche. This presents a versatile vaccine platform applicable to a variety of disease treatment and prevention strategies, thereby expanding the utility of mRNA LNP-based immunotherapies.

## MATERIALS AND METHODS

### Materials

DLin-MC3-DMA was purchased from MedKoo Biosciences. DOPE, DSPC, 18PG, and DMG-PEG-2000 were obtained from Avanti Polar Lipids. Cholesterol was from Sigma-Aldrich. B16F10 (CRL-6475) were purchased from ATCC (American Type Culture Collection, USA). B16-OVA cells were kindly provided by the lab of Prof. Jonathan Schneck. MC38 OVA (KC-2370) was purchased from KYINNO Biotechnology. All mRNA (Cre mRNA, OVA mRNA, Trp2 mRNA, or Gp100 mRNA) constructs were purchased from TriLink BioTechnologies. The anti-CTLA-4 (α-CTLA-4) monoclonal antibody (mctla4-mab10-10) was purchased from InvivoGen. CODEX antibody information was compiled in **Supplementary Table 4.**

Sodium hyaluronate with a molecular weight of 1.5 × 10^6^ Da (HA, research grade) was purchased from LifeCore Biomedical Inc. (Chaska, MN, USA). Glycidyl acrylate was obtained from TCI America Inc. (Portland, OR, USA). Poly(ethylene glycol) dithiol with an average molecular weight of 5 kDa (PEG-SH, MW 5 kDa) was from JenKem Technology (Plano, TX, USA). All other chemical reagents were purchased from Sigma Aldrich (St. Louis, MO, USA) unless otherwise noted.

### LNP synthesis and characterization

LNPs were synthesized by directly adding an organic phase containing the lipids to an aqueous phase containing mRNA in 1.5-mL microcentrifuge tubes. To prepare the organic phase, a mixture of DLin-MC3 DMA, cholesterol, DMG-PEG2000, and a helper lipid selected from a group consisting of DOPE, DSPC, and 18PG were dissolved in ethanol. The mRNA (Cre mRNA, OVA mRNA, Trp2 mRNA, or Gp100 mRNA) was dissolved in 25 mM magnesium acetate buffer (pH 4.0). For larger scale LNP production, the aqueous and ethanol phases prepared were mixed at a 3:1 ratio in a flash complexation (FNC) device using syringe pumps and purified by dialysis against DI water using a 100-kDa MWCO cassette at 4°C for 24 h and were stored at 4°C before injection. The size, polydispersity index, and zeta potentials of LNPs were measured using dynamic light scattering (ZetaPALS, Brookhaven Instruments). Diameters are reported as the intensity average.

### Animals and primary cells

All animal procedures were performed under an animal protocol approved by the Johns Hopkins Institutional Animal Care and Use Committee (protocol #MO22E117). Male and female C57BL/6 mice, 6–8 weeks of age, were purchased from the Jackson Laboratory. Male Ai9 mice, 6–8 weeks of age, were bred in Johns Hopkins Animal Facilities and randomly grouped. The mice were supplied with free access to pelleted feed and water. The pelleted feed generally contained 5% fiber, 20% protein, and 5–10% fat. The mice usually ate 4–5 g of pelleted feed (120 g per kg body weight) and drank 3–5 mL of water (150 mL per kg body weight) per day. The temperature of the mouse rooms was maintained at 18–26°C (64–79°F) at 30–70% relative humidity with a minimum of 10 room air changes per hour. Standard shoebox cages with corncob as bedding were used to house the mice. The LNPs or LiNx were given through *s.c.* (right flank) injection at a predetermined dose per mouse.

### Antibodies, cell isolation, and staining for flow cytometry

Antibodies used in this study are FITC, APC, Brilliant Violet 750 anti-mouse CD11c (BioLegend #117306, 117310, 117357); Brilliant Violet 421 anti-mouse CD86 (BioLegend #105032); PE anti-mouse SIINFEKL H-2KB (ThermoFisher # 12574382); FITC, Brilliant Violet 605, Brilliant Violet 421 anti-mouse CD45 (BioLegend # 103108, 103140, 103134); APC anti-mouse CD3 (BioLegend # 100236); FITC, APC, Brilliant Violet 750 anti-mouse CD8 (BioLegend # 100706, 100712, BD Biosciences # 747502); PerCP-Cyanine 5.5 anti-mouse CD4 (BioLegend # 100540); PE anti-mouse IFN-γ (BioLegend # 505808); Brilliant Violet 421 anti-mouse IL-4 (BioLegend # 504120); PE-Cyanine 7 anti-mouse TNFα (BioLegend # 506324); and APC anti-mouse Granzyme B (BioLegend # 396408). All antibodies were diluted at a ratio of 1:100 before use.

For isolation, re-stimulation, and staining of splenocytes, the spleen was removed and minced using a sterile blade and dissociated in 250 μL of digestion medium (45 units µL^-1^ collagenase I, 25 units µL^-1^ DNase I and 30 units µL^-1^ hyaluronidase). The suspension was transferred into a 15- mL tube containing 5–10 mL of digestion medium and then filtered through a 70-μm filter and washed once with PBS. Cells were pelleted at 300 ×g for 5 min at 4 °C resuspended in 5 mL of red blood cell lysis buffer (BioLegend), and then incubated on ice for 5 min. Cells were then pelleted at 300 ×g for 5 min at 4 °C and washed twice with PBS. Isolated splenocytes were counted using the Cellometer cell counter and ViaStain AOPI staining solution (CS2-0106-25 mL, Nexcelom) and diluted in PBS to be used for restimulation. Splenocytes were re-stimulated *in vitro* with OVA (InvivoGen Cat. vac-pova) and SIINFEKL peptide (InvivoGen Cat. vac-sin) (10 μg mL^-1^ OVA and 2 μg mL^-1^ SIINFEKL) for 12 h. After re-stimulation, cells were collected and centrifuged at 300×g for 5 min. The cell pellet was washed with staining buffer 3 times and stained with antibodies against surface markers (total volume 100 μL) for 30 min in the dark at 4°C. The stained cells were washed twice with 1 mL of PBS and then fixed and permeabilized using the fixation/permeabilization solution kit (BD Cat# 555028). Then, cells were stained with anti-IFN- γ or other antibodies against intracellular cytokines. Flow data were acquired on Attune (ThermoFisher) and analysed using FlowJo software.

For isolation and staining of cells from lymph nodes or LiNx, isolated lymph nodes or LiNx were mechanically digested through 70 μm nylon cell strainers to prepare single-cell suspensions. The cell suspension was washed once with PBS via centrifugation (300×g) for 5 min. Then, the cells were resuspended in 100 µL of staining buffer and stained with antibodies (total volume 100 μL) for 20 min in the dark at 4°C. The stained cells were washed twice with 1 mL of PBS and resuspended in 300 µL of staining buffer for flow cytometry analysis. Flow data were acquired on Attune (ThermoFisher) and analysed using FlowJo software.

### ELISpot assay

Multiscreen filter plates (Millipore-Sigma #S2EM004M99) were coated with antibodies specific for IFN-γ (BD Biosciences #551881) and blocked following the manufacturer’s protocols. Then, 1 × 10^5^ isolated splenocytes were plated per well and stimulated with SIINFEKL peptide (2 μg mL^−1^ SIINFEKL) for 24 h. All tests were performed in duplicate or triplicate and included assay-positive controls as well as cells from a reference donor with known reactivity. Spots were visualized with mouse IFN-γ detection antibody (BD Biosciences #551881) followed by incubation with Streptavidin-HRP (BD Biosciences #557630) and AEC Substrate (BD Biosciences #551951). Plates were then sent to SKCCC Immune Monitoring Core for analysis.

### Enzyme-linked immunosorbent assay (ELISA)

For antibody detection, groups of C57BL/6 mice were immunized with different vaccines on days 0, 7, and 14. On day 21, 100 μL of blood sample was drawn from the tail vein, and levels of antigen-specific IgG in the serum were measured by ELISA. Flat-bottomed 96-well plates (Nunc) were precoated with OVA protein at a concentration of 2 μg protein per well in 100 mM carbonate buffer (pH 9.6) at 4°C overnight, which were then blocked with 10% fetal bovine serum (FBS) in PBS-Tween (PBS-T). Sera obtained from immunized animals were diluted 100 times in PBS-T (PBS-0.05% Tween), pH 7.4, and then in 4-fold serial dilution. The undiluted and diluted serum was added to the wells and incubated at 37°C for 2 h. Horseradish peroxidase-conjugated goat anti-mouse IgG (Southern Biotech Associates, #1013-05) was used at a dilution of 1:5,000 in PBS-T-10% FBS for labeling. After adding the horseradish peroxidase substrates, optical densities were determined at a wavelength of 450 nm in an ELISA plate reader (Bio-Rad). A sample is considered positive if its absorbance is twice as much as or higher than the absorbance of the negative control.

### Immunization and tumour therapy experiments

Mice aged 6–8 weeks were injected subcutaneously with B16-OVA, MC38-OVA cells (1 × 10^6^ in the prophylactic model and 3 × 10^5^ in the therapeutic model) or 3 × 10^5^ B16F10 melanoma cells into the right flank. In therapeutic studies, vaccinations began when tumour sizes were less than 50 mm^3^ (on day 4 after tumour inoculation). Animals were immunized by subcutaneous injection of different LNP or LiNx formulations containing OVA mRNA, mTrp2, or m Gp100 as described in the main text. One dosage of LiNx was given, and a total of three doses were given for the LNP group. For combinatorial immunotherapy, at days 6, 13, and 20 and an additional at day 27 for OVA- expressing melanoma after inoculation, some groups were intraperitoneally injected with 100 μg checkpoint inhibitor (α-CTLA-4 mAb). Tumour growth was measured three times a week using a digital caliper and calculated as 0.5 × length × width × width. Mice were euthanized when the tumour volumes reached 2,000 mm^3^.

### IL-17 depletion study

Depletions of IL-17 were done using α-IL-17A (clone 17F3, BioXCell) at 200 μg *i.p.* every 3 d. The dosing was initiated at 3 d after the first vaccination and continued every 3 d until day 60. On day 21 post-vaccination, 3 × 10^5^ B16-OVA melanoma cells were injected into the right flank. Tumour growth was measured three times a week using a digital caliper and calculated as 0.5 × length × width × width. Mice were euthanized when the tumour volumes reached 2,000 mm^3^.

### Subsequent RT-PCR analysis

Real-time polymerase chain reaction (RT-PCR) was carried out using TaqMan™ Array Mouse Immune Response (Applied Biosystems, Cat#4414079). The results were analysed according to the 2−ΔΔCT method and normalized to the housekeeping gene GAPDH.

### nCounter Analysis System (NanoString Technology)

Total RNA was extracted from stimulated spleen cells using an RNA extraction kit (Zymo Research, Cat# R2062). Quality and concentration were evaluated using NanoDrop. Predesigned NanoString nCounter CodeSets targeting mouse immune responses-related genes were employed (NanoString, Cat#115000052). Hybridization of RNA samples to these CodeSets was performed following the manufacturer’s protocol. Post-hybridization, samples were processed on the nCounter Analysis System, and data normalization was conducted using nSolver Analysis Software with reference to housekeeping genes.

### CODEX multiplexed imaging

Tumour tissues were collected, embedded in optimal cutting temperature (OCT) compound for CODEX, and immediately frozen. After freezing, explants were sectioned to a thickness of 7 µm using an Epredia^TM^ HM525 NX cryostat and arranged on slides. The tissue arrays were stained with the validated panels of CODEX antibodies and imaged in accordance with a previously established protocol.^50^ Briefly, this entailed cyclic stripping, annealing, and imaging of fluorescently labeled oligonucleotides complementary to the oligonucleotide conjugated to the antibody. Each array underwent CODEX multiplexed imaging; metadata from each CODEX run can be found in **Supplementary Table 2**. Raw imaging data were processed using the Akoya PhenoCylcer Fusion 2.2.0 software for image stitching, drift compensation, deconvolution, and cycle concatenation. After the raw imaging data were processed, they were evaluated for specific signals. Any markers that produced an untenable pattern or a low signal-to-noise ratio were excluded from the ensuing analysis. Uploaded images were visualized in ImageJ (https://imagej.net/software/fiji/).

### CODEX single-cell segmentation

To obtain quantitative single-cell information, individual cells and extracted single-cell protein expression were segmented. Processed data were segmented using the SPACEc package, which can also be downloaded here (https://github.com/yuqiyuqitan/SPACEc/tree/master).^51^ SPACEc incorporates Mesmer and Cellpose, which are both deep learning-based segmentation methods.^52,53^ Mesmer was used for our segmentation, with DAPI (nuclei) along with CD45 and CD90 (surface membrane) as reference channels.

### Cell-type analysis

Across the LiNx and PBS-treated tumours, 208,486 cells were identified and classified into 19 cell types and states (197652 cells) with noise excluded based on marker expression. Cell type identification was done following the strategies that were developed.^54,55^ Briefly, segmented cells of appropriate sizes were selected by gating DAPI-positive cells, followed by Z-normalization of protein markers used for clustering (some phenotypic markers were not used in the unsupervised clustering). The data were overclustered with Leiden-based clustering with the scanpy Python package. Clusters were assigned a cell type based on average cluster protein expression and location within the image. Impure clusters were split or reclustered with K-means clustering with the sklearn Python package following mapping back to the original fluorescent images.

### Statistical analysis

A two-tailed Student’s t-test or a one-way analysis of variance (ANOVA) was performed when comparing two groups or more than two groups, respectively. Survival curves were compared using the log-rank Mantel-Cox test, and the stack of *P* values was corrected by the Holm-Šídák method for multiple comparisons with alpha set to 0.05. Statistical analysis was performed using Microsoft Excel and Prism 8.0 (GraphPad). A difference is considered significant if *P<* 0.05 (\**P* < 0.05, \*\**P* < 0.01, \*\*\**P* < 0.001, \*\*\*\**P* < 0.0001).

## Author contributions

Y.Z. and H.-Q.M. conceived of and designed this study. H.-Q.M. and J.P.S. secured the funding for this study. Y.Z., R.S., J.M., Z.-C.Y., S. L., I.V., J.L., X.L., K.D., X.L., D.Y., C.W., J.L.S., B.Y.X.N., Y.M., K.V.B., J.W.H., and J.K. performed the experiments. Y.Z., R.S., Y.D., J.M., I.V., N.K.L., S.L., G.P.H., S.K.R., J.C., J.P.S., J.C.D., B.Y.X.N., Y.M., J.W.H., and H.-Q.M. participated in data analysis and interpretation. The manuscript was written by Y.Z. and H.-Q.M., with revisions by J.W.H., B.Y.X.N., C.W., J.M., J.P.S., J.C.D., S.K.R., L.C., and inputs from all the other authors.

## Data and materials availability

All data needed to evaluate the conclusions in the paper are present in the paper and/or the Supplementary Materials.

## Code availability statement

There is no code used in the paper.

