## Supporting information for "mRNA lipid nanoparticle-incorporated nanofiber-hydrogel composite generates a local immunostimulatory niche for cancer immunotherapy"

### Supplementary Figures


**Figure 1.** Rheological assessment and injectability of LiNx formulation (D6 LiNx).

**Figure 2.** Scanning electron microscopy (SEM) images of D6 LiNx.

**Figure 3.** Accelerated degradation profile at 50 °C for the NHC composite used in this study.

**Figure 4.** Gating strategy for flow cytometry plots for in vivo assessments of locally recruited cells by the top LiNx formulations on days 3 and 7.

**Figure 5.** Gating strategy for flow cytometry plots for in vivo assessments of local cell transfection by the top LiNx formulations on days 5 and 10.

**Figure 6.** Representative flow cytometry plots for TdTomato$^+$ cells in LiNx on days 5 and 10 post-administration.

**Figure 7.** *In vivo* transfection of CD11c$^-$CD11b$^-$ cells by C10, D6, and F5 mCre LiNx formulations.

**Figure 8.** Local retention and biodistribution profile of the labelled mRNA LNPs formulated in the D6 LiNx post-vaccination.

**Figure 9.** Gating strategy for flow cytometry plots for in vivo assessments of locally recruited CD4$^+$ T cells by the LiNx formulations on day 14.

**Figure 10.** Gating strategy for flow cytometry plots for in vivo assessments of locally recruited CD8$^+$ T cells by the LiNx formulations on day 14.

**Figure 11.** Gating strategy for flow cytometry plots for in vivo assessments of locally recruited B cells by the LiNx formulations on day 14.

**Figure 12.** Gating strategy for flow cytometry plots for in vivo assessments of locally recruited NK cells, macrophages, DC cells, and neutrophils by LiNx formulations on day 14.

**Figure 13.** RT-PCR analysis of selected genes related to inflammatory cytokines and chemokines expression in the local microenvironment generated within the LiNx.

**Figure 14.** RT-PCR analysis of selected genes related to Th1 immune responses in the local microenvironment generated by the LiNx.

**Figure 15.** RT-PCR analysis of selected genes related to Th2 immune responses in the local microenvironment generated by the LiNx.

**Figure 16.** Gating strategy for flow cytometry assessment of OVA-specific T cells in the LiNx or draining lymph nodes on day 14 post-administration.

**Figure 17.** Representative flow cytometry plots for OVA-specific T cells in the LiNx or draining lymph nodes on day 14 post-administration.

**Figure 18.** *In vivo* assessments of antigen-specific immune activation by LiNx loaded with mOVA D6 LNPs or empty D6 LNPs.

**Figure 19.** Gating strategy for flow cytometry assessments of antigen-specific immune responses generated by the LiNx formulations.

**Figure 20.** Representative images of IFN-γ secreting cells from the enzyme-linked immunospot assay.

**Figure 21.** Titers of OVA-specific IgG subclass antibodies in serum samples collected on day 30 following immunization with the LiNx formulations.

**Figure 22.** Gating strategy for flow cytometry analysis for *in vivo* assessment of OVA-specific $CD8^+$ T cells by the LiNx formulations in the spleen on day 90.

**Figure 23.** Representative flow cytometry plots for *in vivo* assessment of OVA-specific T cells in the spleen on day 90 post-administration.

**Figure 24.** Gating strategy for flow cytometry assessment of cytotoxic T cell response on day 90 post-vaccination.

**Figure 25.** Representative flow cytometry plots for assessment of $CD3^+CD8^+IFN\text{-}\gamma^+$ cells on day 90 post-vaccination.

**Figure 26.** Representative flow cytometry plots for assessment of $CD3^+CD8^+TNF\text{-}\alpha^+$ cells on day 90 post-vaccination.

**Figure 27.** Titers of OVA-specific IgG subclass antibodies in serum samples collected on day 90 post-vaccination.

**Figure 28.** Body weights of mice vaccinated with the mOVA LiNx formulations.

**Figure 29.** Serum cytokine levels of mice vaccinated with the D6 mOVA LiNx formulation at 24 h post-vaccination.

**Figure 30.** Anti-tumour efficacy of the LiNx formulations as therapeutic vaccines for MC38-OVA tumour model.

**Figure 31.** Anti-tumour efficacy of the D6 LiNx as a therapeutic vaccine for B16-OVA tumour model.

**Figure 32.** Anti-tumour efficacy of the D6 LiNx as a therapeutic vaccine for B16F10 tumour model.

**Figure 33.** Anti-tumour efficacy of the D6 LiNx as a prophylactic vaccine for B16-OVA tumour model.

**Figure 34.** Anti-tumour efficacy of the LiNx loaded with either mOVA D6 LNPs or empty D6 LNPs in a prophylactic vaccine model for B16-OVA tumour.

**Figure 35.** Prophylactic efficacy of D6 LiNx in B16-OVA tumour model and resistance to rechallenge.

**Figure 36.** Representative images from the CODEX fluorescence imaging analysis on tumour samples collected.

**Figure 37.** Cell type map of a representative PBS-treated tumour sample from CODEX multiplexed imaging experiment.

**Figure 38.** Immunocytes profile of the blood samples extracted from mice treated with or without α-IL-17 antibody.

### Supplementary Table

**Table 1.** Formulation details and particle sizes for the three selected LNPs CODEX staining conditions and cycle information.

**Table 2.** CODEX staining conditions and cycle information.

**Table 3.** CODEX marker staining evaluation.

**Table 4.** CODEX antibody information.

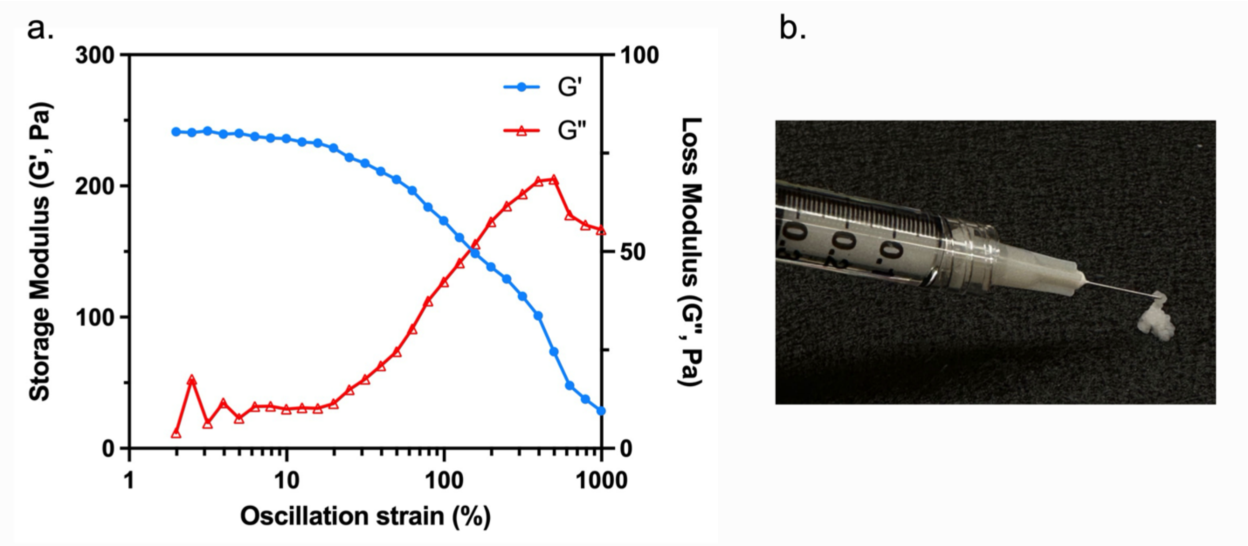


**Supplementary Figure 1. Rheological assessment and injectability of the LiNx formulation (D6 LiNx). (a)** Rheological profile of D6 LiNx, showing storage modulus (G’) and loss modulus (G'') across an oscillatory strain (δ) range of 0–1000% in a strain sweep test. **(b)** Image showing the injectability of the composite through a 30-gauge needle.

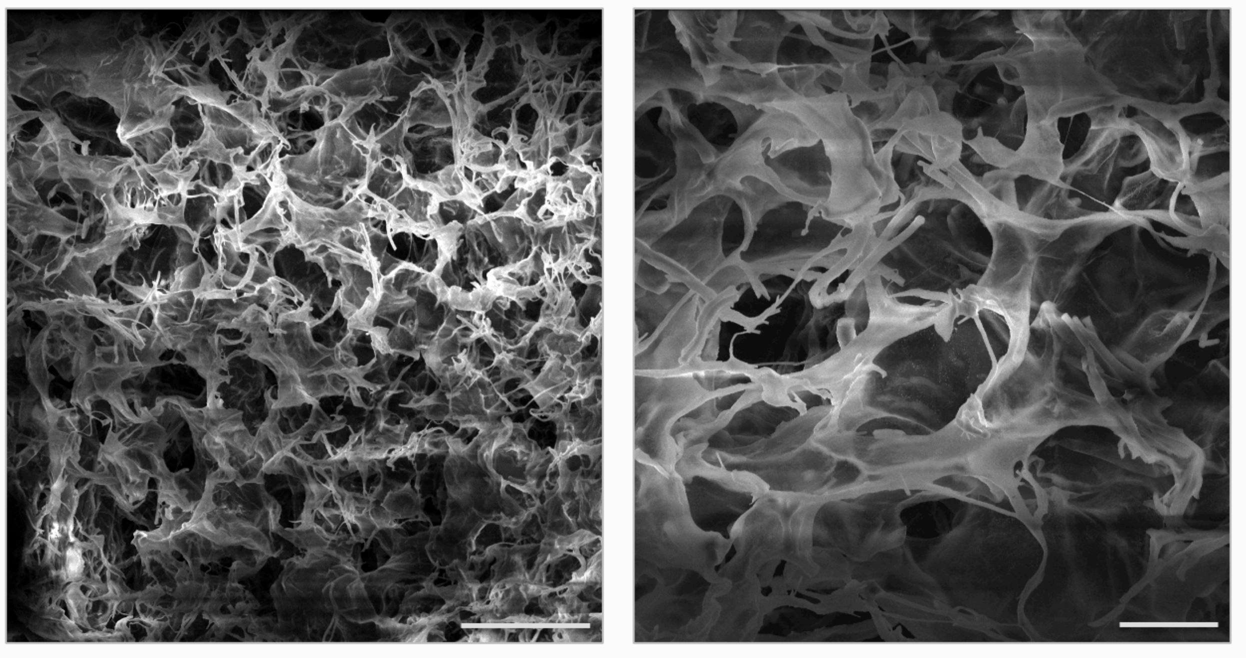

**Supplementary Figure 2. Scanning electron microscopy (SEM) images of D6 LiNx.** The SEM imaging was taken following D6 mRNA LNPs incorporation into the NHC composite. Scanning electron microscopy images showed that, after encapsulating LNPs within the NHC, the fibrillar microarchitecture of the NHC is preserved, with nanofibers connected to the HA hydrogel network. Scale bars: 100 µm (left) and 25 µm (right).

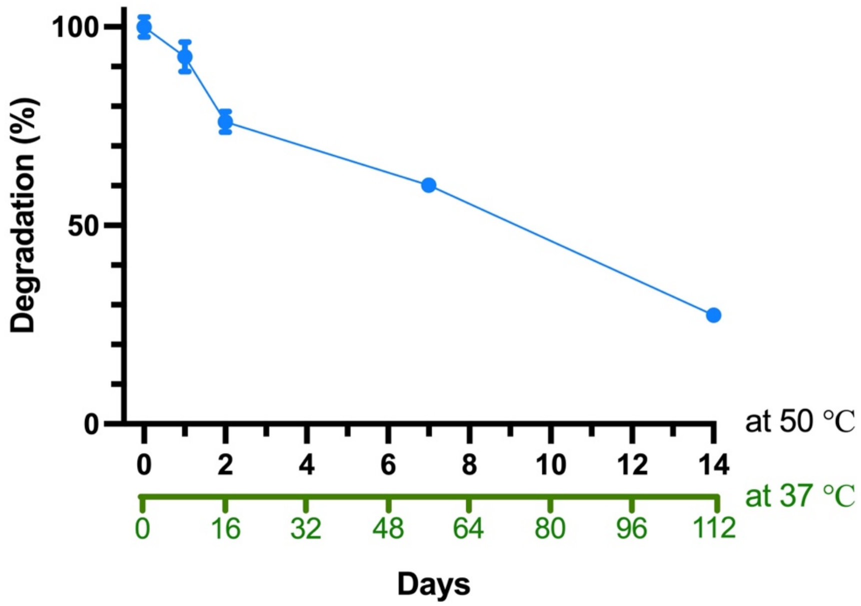


**Supplementary Figure 3. Accelerated degradation profile at 50 °C for the NHC composite used in this study.** Rheological analysis of NHC degradation was conducted at 50 °C. Samples were incubated at 50 °C in syringes and periodically tested using a rheometer to measure the storage modulus (G'). A decrease in G' over time indicates reduced stiffness, reflecting the degradation profile of the NHC scaffold. The equivalent degradation time at 37 °C was estimated based on a scaling factor. Data represent the mean ± s.d. (n = 3).

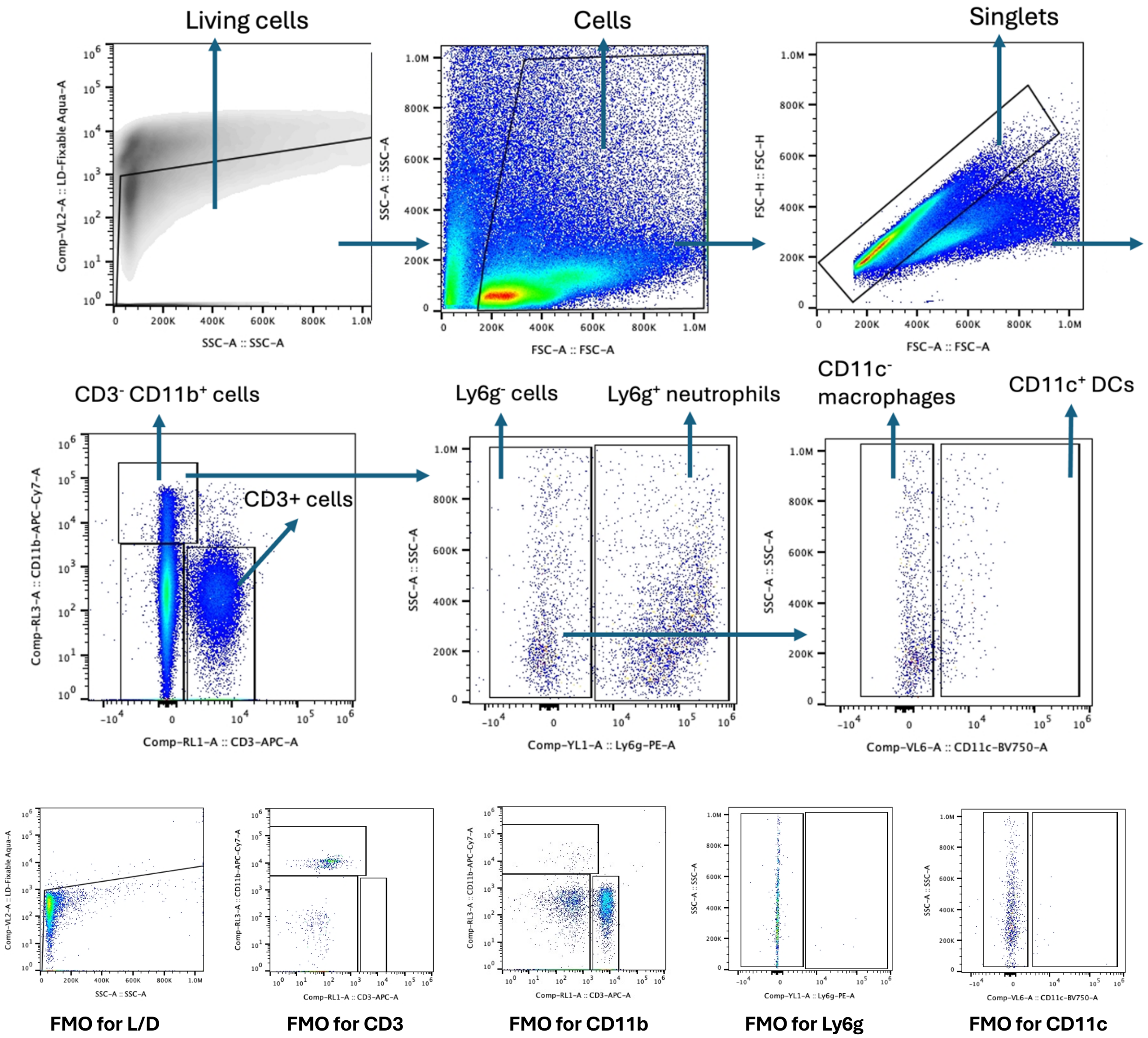


**Supplementary Figure 4. Gating strategy for flow cytometry assessment of locally recruited cells by the top LiNx formulations on days 3 and 7.** Initially, viable cells were identified and gated out based on the L/D aqua-A–SSC-A plot. Lymphocytes were gated out using SSC-A and FSC-A parameters. Subsequently, singlet cells were gated out using the FSC-H–FSC-A plot. Next, $CD3^-$ $CD11b^+$ cells were gated, and $Ly6g^+$ cells were further characterized as neutrophils. Additionally, $Ly6g^-CD11c^+$ cells were identified as dendritic cells (DCs), while $Ly6g^-CD11c^-$ cells were designated as macrophages.

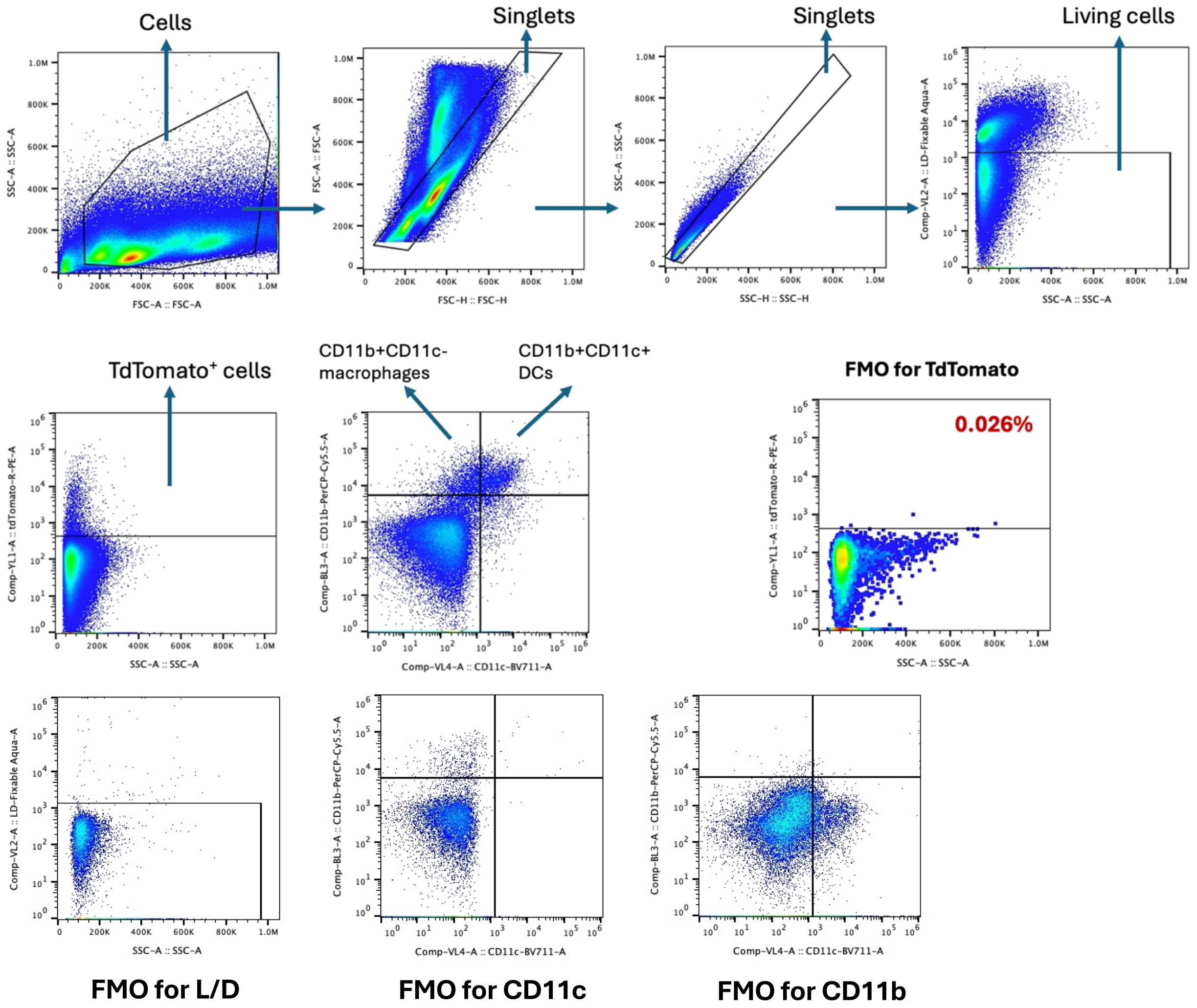


**Supplementary Figure 5. Gating strategy for flow cytometry assessment of local cell transfection by the top LiNx formulations**. Initially, lymphocytes were gated out using SSC-A and FSC-A parameters. Subsequently, singlet cells were gated out using the FSC-H–FSC-A plot and SSC-A–SSC-H plot. Viable cells were identified and gated out based on the L/D aqua-A–SSC-A plot. The analysis focused on tdTomato$^+$ CD11b$^+$CD11c$^-$ macrophages and CD11b$^+$CD11c$^+$ DCs. The fluorescence minus one (FMO) control for tdTomato$^+$ cells was provided.

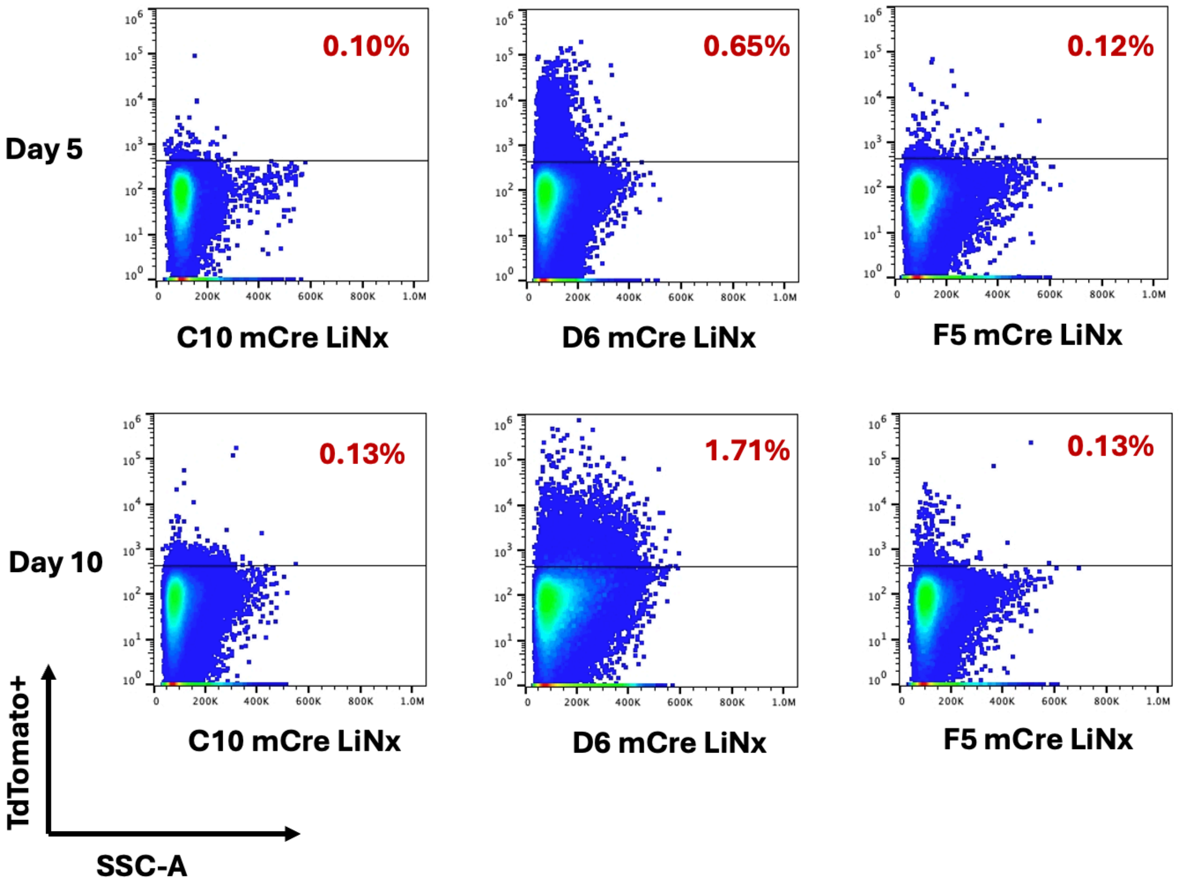


**Supplementary Figure 6. Representative flow cytometry plots for tdTomato⁺ cells in LiNx on days 5 and 10 post-administration.** Ai9 mice were administered the top three LiNx formulations loaded with mCre via *s.c.* injections (n = 4, 30 µg mCre per mouse). Transfection of cells in LiNx was analysed by flow cytometry. Percentages of cells positive for tdTomato are shown.

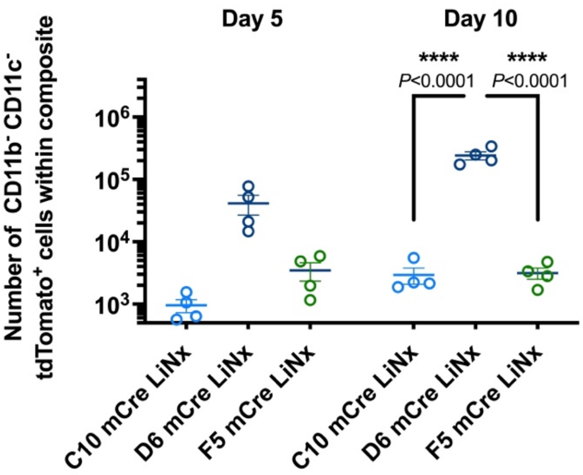


**Supplementary Figure 7. In vivo transfection of CD11c⁻CD11b⁻ cells by C10, D6, and F5 mCre LiNx formulations**. Ai9 mice were administered with LiNx loaded with C10, D6, or F5 mCre LNPs via s.c. injections (30 µg mCre per mouse). Transfection of CD11c⁻CD11b⁻ cells in the LiNx was analysed by flow cytometry. The number of CD11c⁻CD11b⁻ cells positive for tdTomato on days 5 and 10 post-injection were shown. Data represent mean ± s.e.m. (n = 4 biologically independent samples). Data were analysed using one-way ANOVA and Tukey's multiple comparisons test. *P < 0.05, **P < 0.01, ****P < 0.0001. NS, not significant.

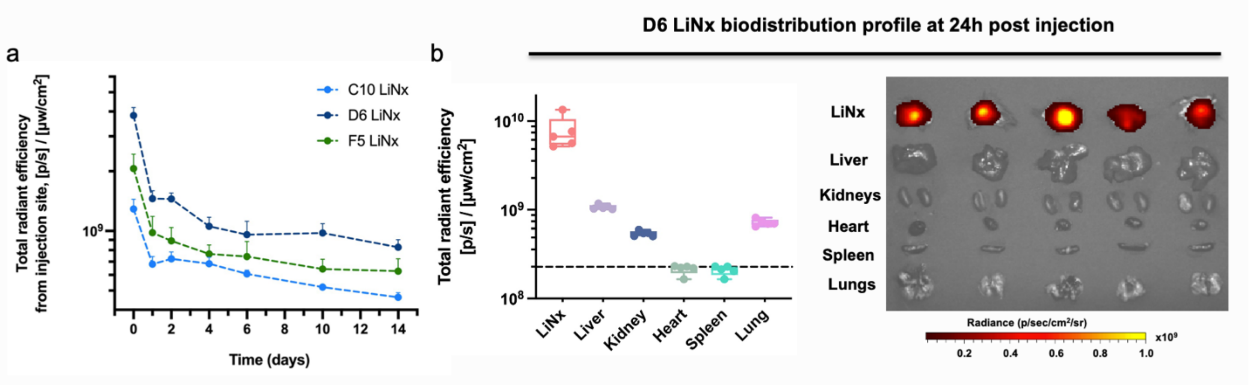


**Supplementary Figure 8. Local retention and biodistribution profile of the labelled mRNA LNPs formulated in the D6 LiNx post-vaccination.** C57BL/6 mice received *s.c.* injections of D6 LiNx loaded with Cy5-labeled mRNA (30 µg per mouse). The Cy5 radiant efficiency from the D6 LiNx injection (a) was monitored over 2 weeks using IVIS imaging. (b) Cy5 radiant efficiency in major organs and at the injection site was assessed at 24 hours post-injection. IVIS biodistribution images of D6 LiNx at 24 hours post-injection were shown in (c). Data represent the mean ± s.e.m. (n = 5 biologically independent samples).

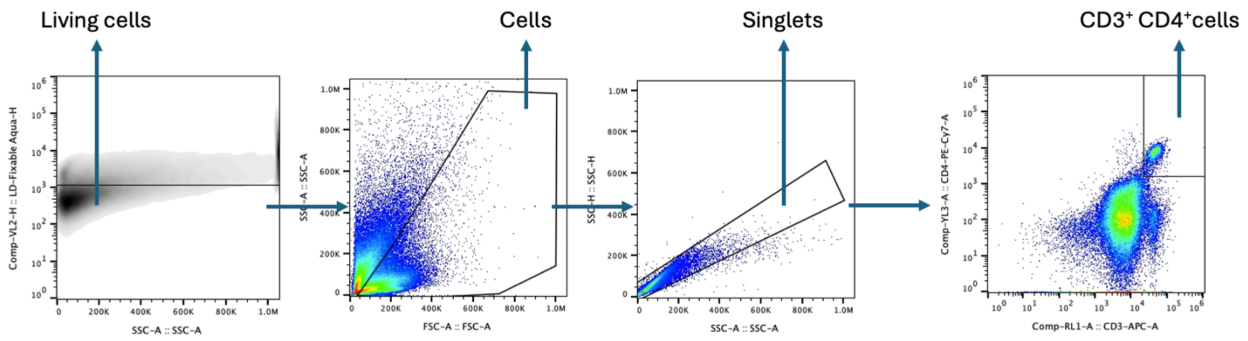


**Supplementary Figure 9. Gating strategy for flow cytometry assessment of locally recruited CD4⁺ T cells by the LiNx formulations on day 14.** Initially, viable cells were identified and gated out based on the L/D aqua-A–SSC-A plot. Lymphocytes were gated out using SSC-A and FSC-A parameters. Subsequently, singlet cells were gated out using the SSC-H–SSC-A plot. The CD3⁺CD4⁺ cells were characterized as CD4⁺ T cells.

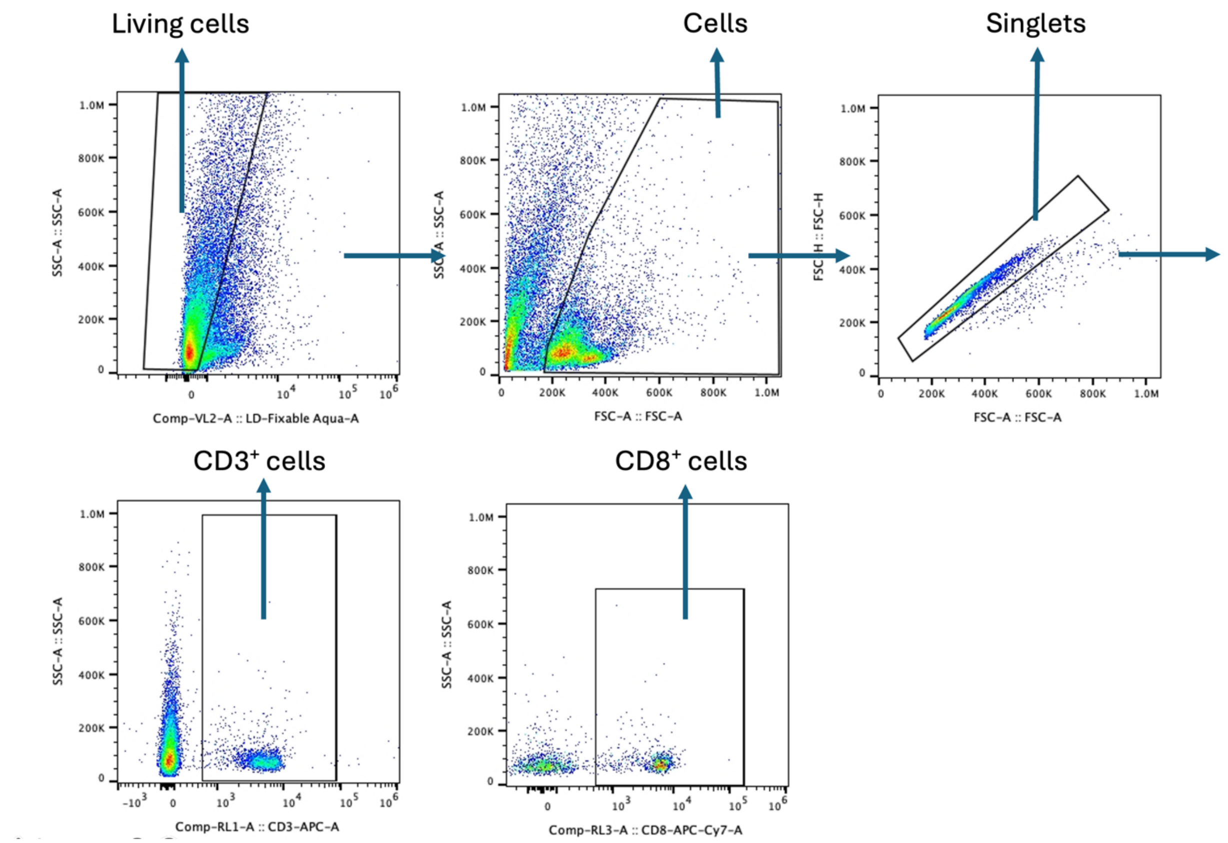


**Supplementary Figure 10. Gating strategy for flow cytometry assessment of locally recruited CD8⁺ T cells by the top LiNx formulations on day 14.** Initially, viable cells were identified and gated out based on the L/D aqua-A–SSC-A plot. Lymphocytes were gated out using SSC-A and FSC-A parameters. Subsequently, singlet cells were gated out using the FSC-H–FSC-A plot. Next, CD3⁺ cells were gated, and CD3⁺ CD8⁺ cells were characterized as CD8⁺ T cells.

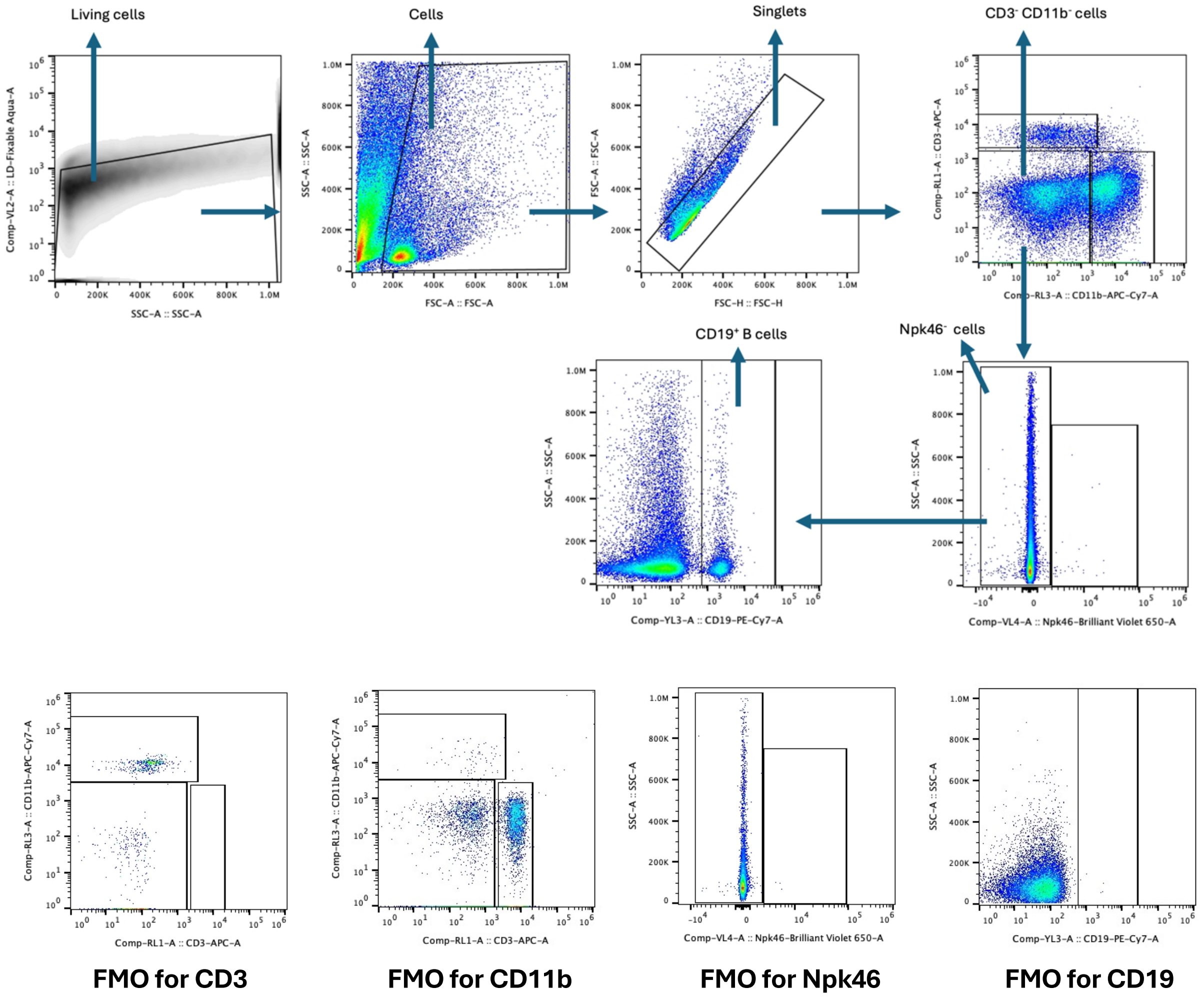


**Supplementary Figure 11. Gating strategy for flow cytometry assessment of locally recruited B cells by the LiNx formulations on day 14.** Initially, viable cells were identified and gated out based on the L/D aqua-A–SSC-A plot. Lymphocytes were gated out using SSC-A and FSC-A parameters. Subsequently, singlet cells were gated out using the FSC-H–FSC-A plot. Next, $CD3^-$ $CD11b^-$ cells were gated, and $Npk46^-$ $CD19^+$ cells were characterized as B cells.

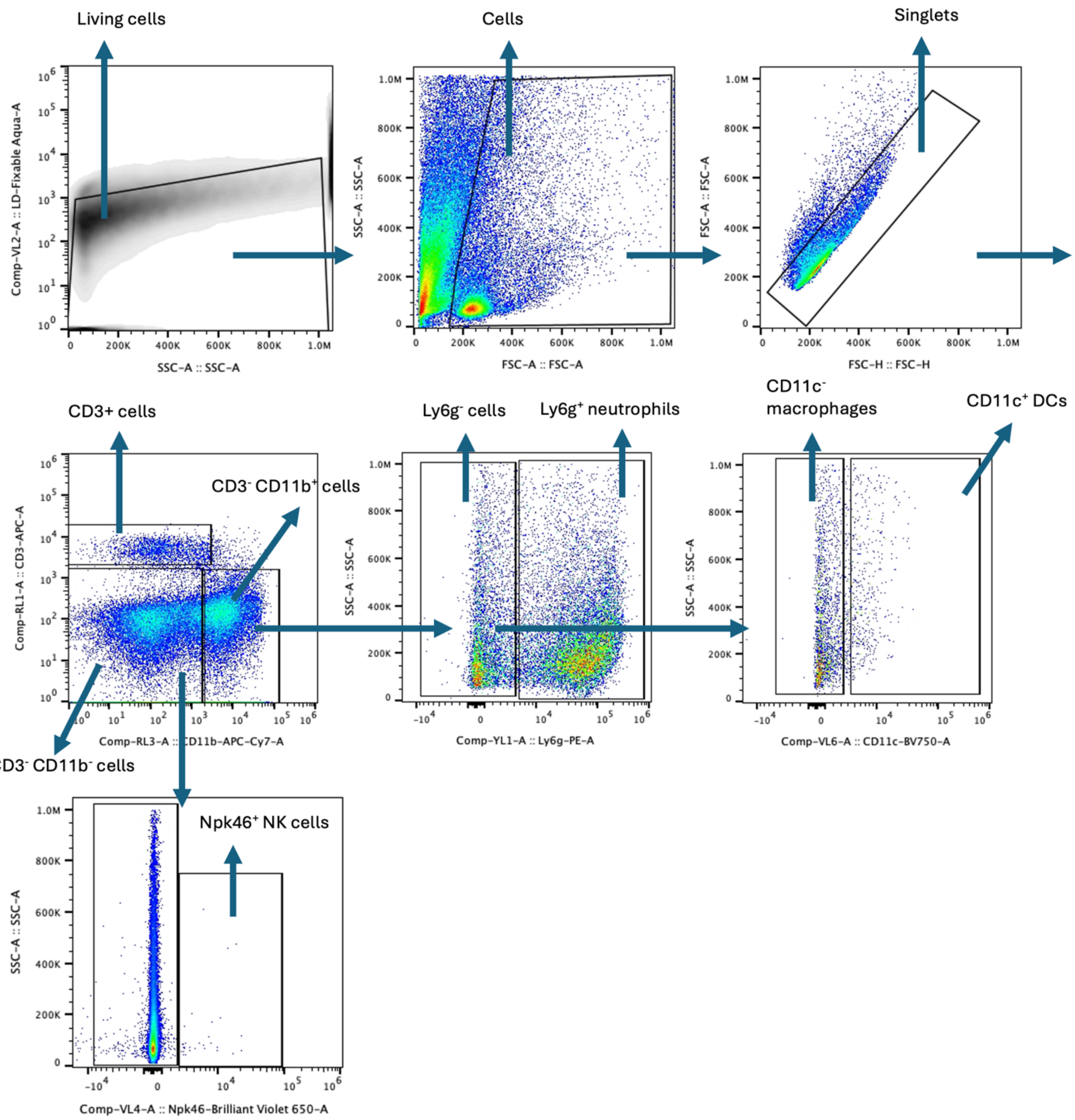


**Supplementary Figure 12. Gating strategy for flow cytometry assessment of locally recruited NK cells, macrophages, DC cells, and neutrophils by the LiNx formulations on day 14.** Initially, viable cells were identified and gated out based on the L/D aqua-A–SSC-A plot. Lymphocytes were gated out using SSC-A and FSC-A parameters. Subsequently, singlet cells were gated out using the FSC-H–FSC-A plot. First, $CD3^{-}CD11b^{+}$ cells were gated, and within this population, $Ly6g^{+}$ cells were identified and characterized as neutrophils. Subsequently, $Ly6g^{-}$ cells were gated. Among these, $CD11c^{+}$ cells were identified as dendritic cells (DCs), while $CD11c^{-}$ cells were identified as macrophages. Finally, within the $CD3^{-}CD11b^{-}$ cell subset, $Npk46^{+}$ cells were identified and characterized as NK cells.

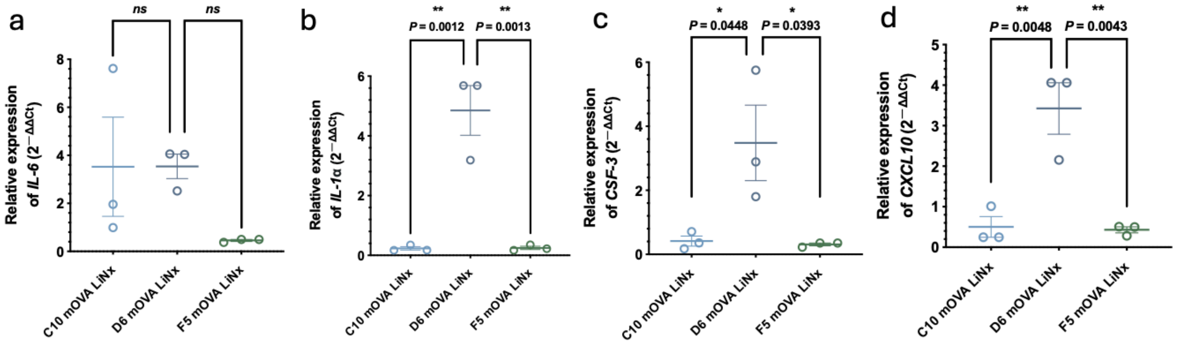


**Supplementary Figure 13. RT-PCR analysis of selected genes related to inflammatory cytokines and chemokines expression in the local microenvironment generated within the LiNx.** An RT-PCR array was conducted using RNA isolated from the local microenvironment generated by three distinct LiNx formulations. C57BL/6 mice received the three different LiNx formulations loaded with mOVA through *s.c.* injection (30 μg mOVA per injection). The selected genes related to inflammatory cytokines and chemokines expression in the local microenvironment generated by the LiNx, including IL-6 (a), IL-1α (b), CSF-3 (c), and CXCL10 (d), are shown. Data represent the mean ± s.e.m. (n = 3 biologically independent samples). Data were analysed using one-way ANOVA and Tukey's multiple comparisons. **P* < 0.05, ***P* < 0.01. NS, not significant.

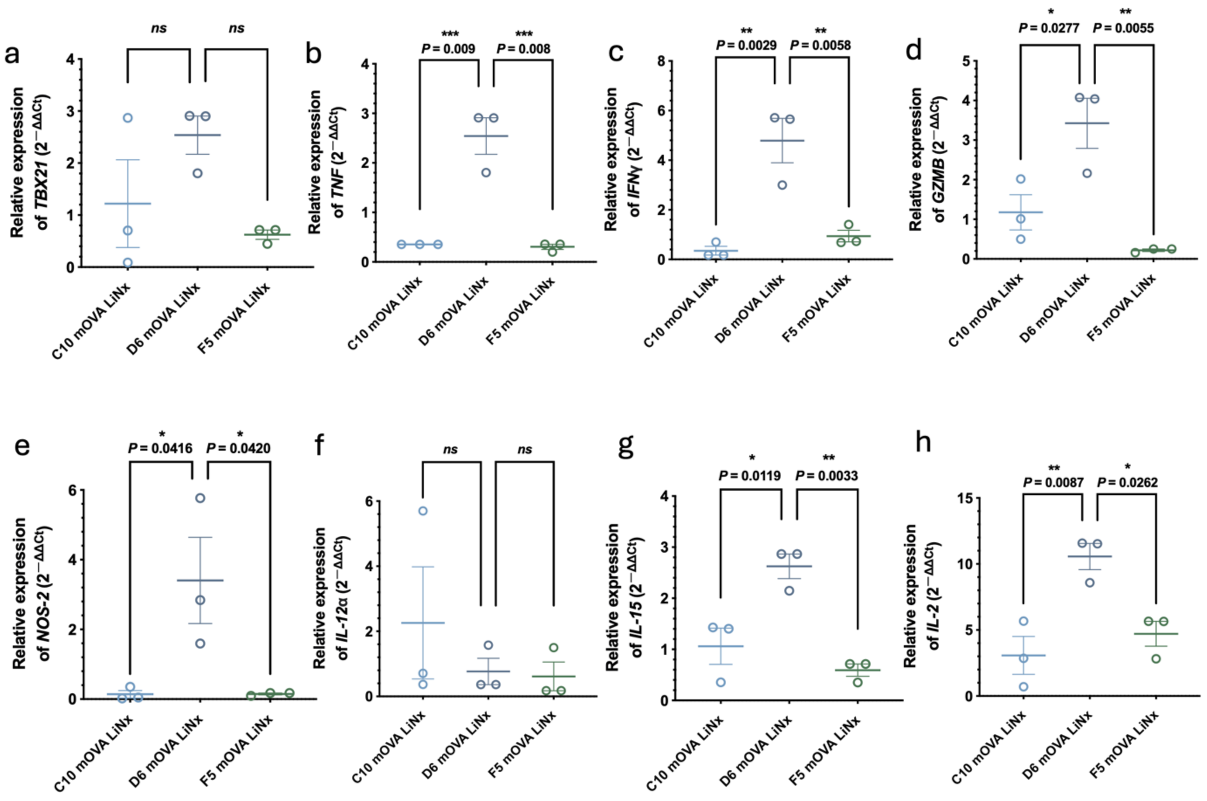


**Supplementary Figure 14. RT-PCR analysis of selected genes related to Th1 immune responses in local microenvironment generated by the LiNx.** An RT-PCR array was conducted using RNA isolated from the local microenvironment generated by three distinct LiNx formulations. C57BL/6 mice received three different LiNx formulations loaded with mOVA through *s.c.* injection (30 µg mOVA per injection). The selected genes related to Th1 immune responses in local microenvironment generated by the LiNx, including TBX21 (a), TNF (b), IFN-γ (c), GZMB (d), NOS-2 (e), IL-12α (f), IL-15 (g), and IL-2 (h), are shown. Data represent the mean ± s.e.m. (n = 3 biologically independent samples). Data were analysed using one-way ANOVA and Tukey's multiple comparisons test. **P* < 0.05, ***P* < 0.01, ****P* < 0.001. NS, not significant.

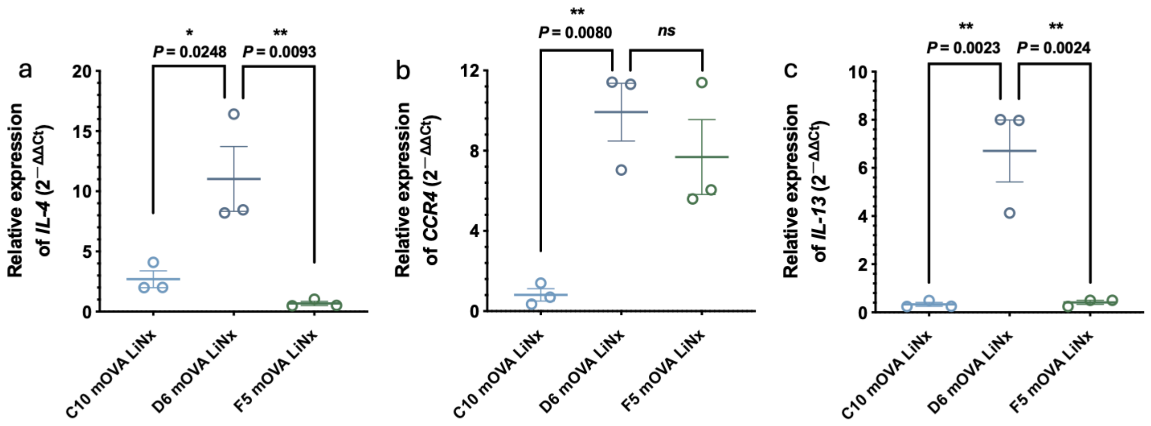


**Supplementary Figure 15. RT-PCR analysis of selected genes related to Th2 immune responses in the local microenvironment generated by the LiNx.** An RT-PCR array was conducted using RNA isolated from the local microenvironment generated by three distinct LNP/mRNA LiNx formulations. C57BL/6 mice received three different LiNx formulations loaded with C10, D6, and F5 mOVA LNPs through *s.c.* injection (30 µg mOVA per injection). The selected genes related to Th2 immune responses in the local microenvironment generated by LiNx, including IL-4 (a), CCR-4 (b), and IL-13 (c), are shown. Data represent the mean ± s.e.m. (n = 3 biologically independent samples). Data were analysed using one-way ANOVA and Tukey's multiple comparisons test. $*P < 0.05$, $**P < 0.01$. NS, not significant.

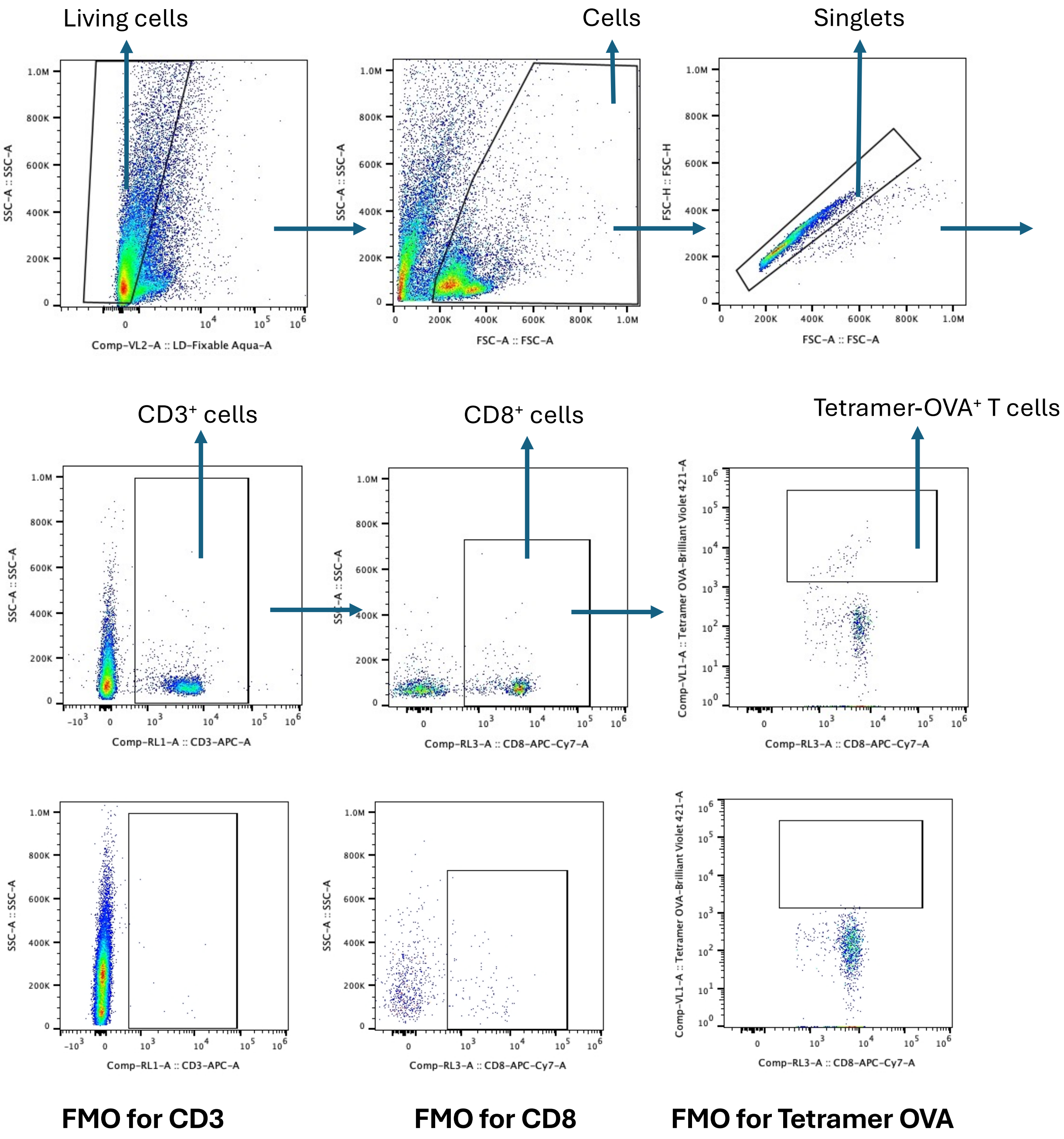


**Supplementary Figure 16. Gating strategy for flow cytometry assessment of OVA-specific T cells in the LiNx or draining lymph nodes on day 14 post-administration.** Initially, viable cells were identified and gated out based on the L/D aqua-A–SSC-A plot. Lymphocytes were gated out using SSC-A and FSC-A parameters. Subsequently, singlet cells were gated out using the FSC-H–FSC-A plot. Next, $CD3^+$ cells $CD8^+$ cells were gated; and $CD3^+$ $CD8^+$ OVA+ cells were characterized as OVA specific T cells.

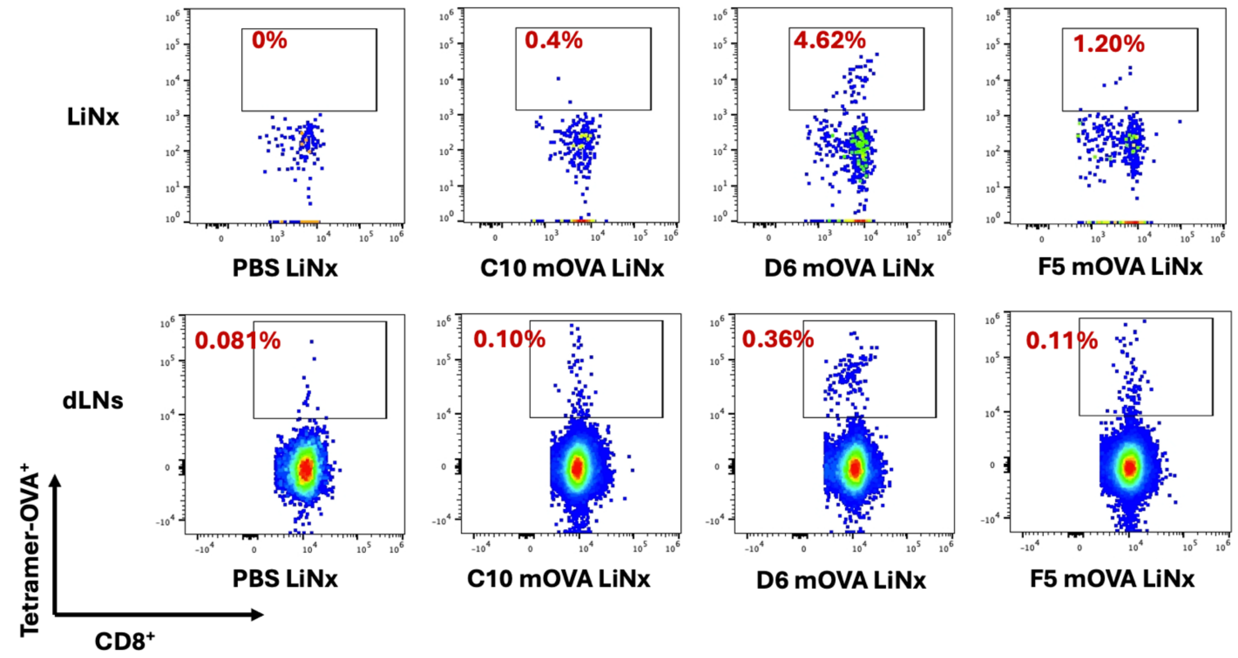


**Supplementary Figure 17. Representative flow cytometry plots for OVA-specific T cells in the LiNx or draining lymph nodes on day 14 post-administration.** C57BL/6 mice were administered with the three LiNx formulations loaded with C10, D6, and F5 mOVA LNPs via *s.c.* injections (n = 7, 30 µg mOVA per mouse). OVA-specific T cells in the LiNx and dLNs were analysed by flow cytometry. Percentages of cells positive for OVA tetramer are shown.

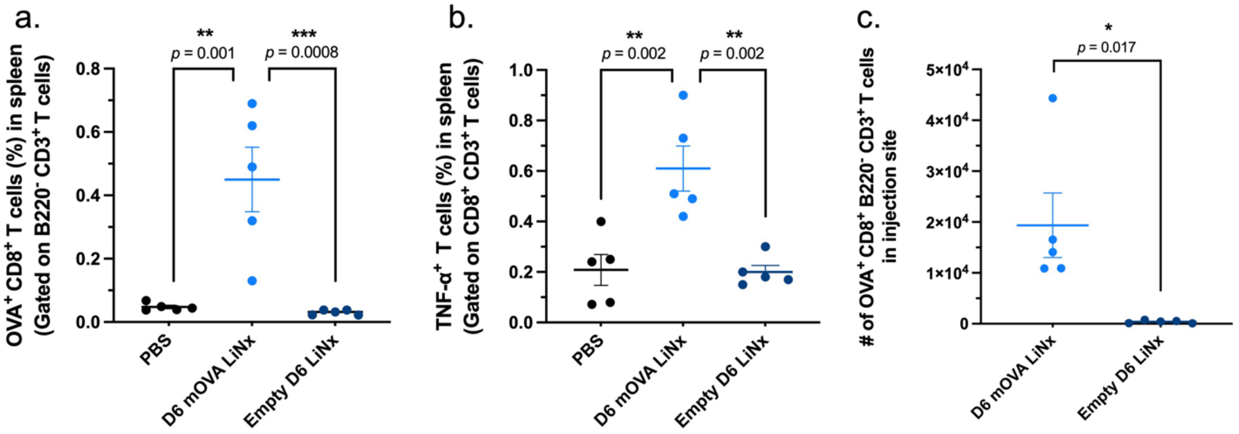


**Supplementary Figure 18. *In vivo* assessments of antigen-specific immune activation by the LiNx loaded with mOVA D6 LNPs or empty D6 LNPs. a–c,** C57BL/6 mice were administered with the D6 LiNx formulation loaded with D6 mOVA LNPs (30 µg mOVA per mouse) or empty D6 LNPs via *s.c.* injection (equivalent dose of LNPs). Mice were sacrificed two weeks after the injection, and their splenocytes were isolated and restimulated in vitro with OVA and SIINFEKL peptides (100 µg $mL^{-1}$ OVA and 2 µg $mL^{-1}$ SIINFEKL) for 12 hours and assessed via flow cytometry and intracellular cytokine staining to determine the percentages of OVA-specific CD8 T cells **(a)**, $CD8^{+}TNF\alpha^{+}$ cells **(b)**. The number of OVA-specific CD8 T cells within the LiNx **(c)** was analysed through flow cytometry at two weeks following a single dosage of D6 LiNx. Data represent the mean ± s.e.m. (n = 5 biologically independent samples). Data were analysed using one-way ANOVA and Tukey's multiple comparisons test. *$P < 0.05$, **$P < 0.01$, ***$P < 0.001$.

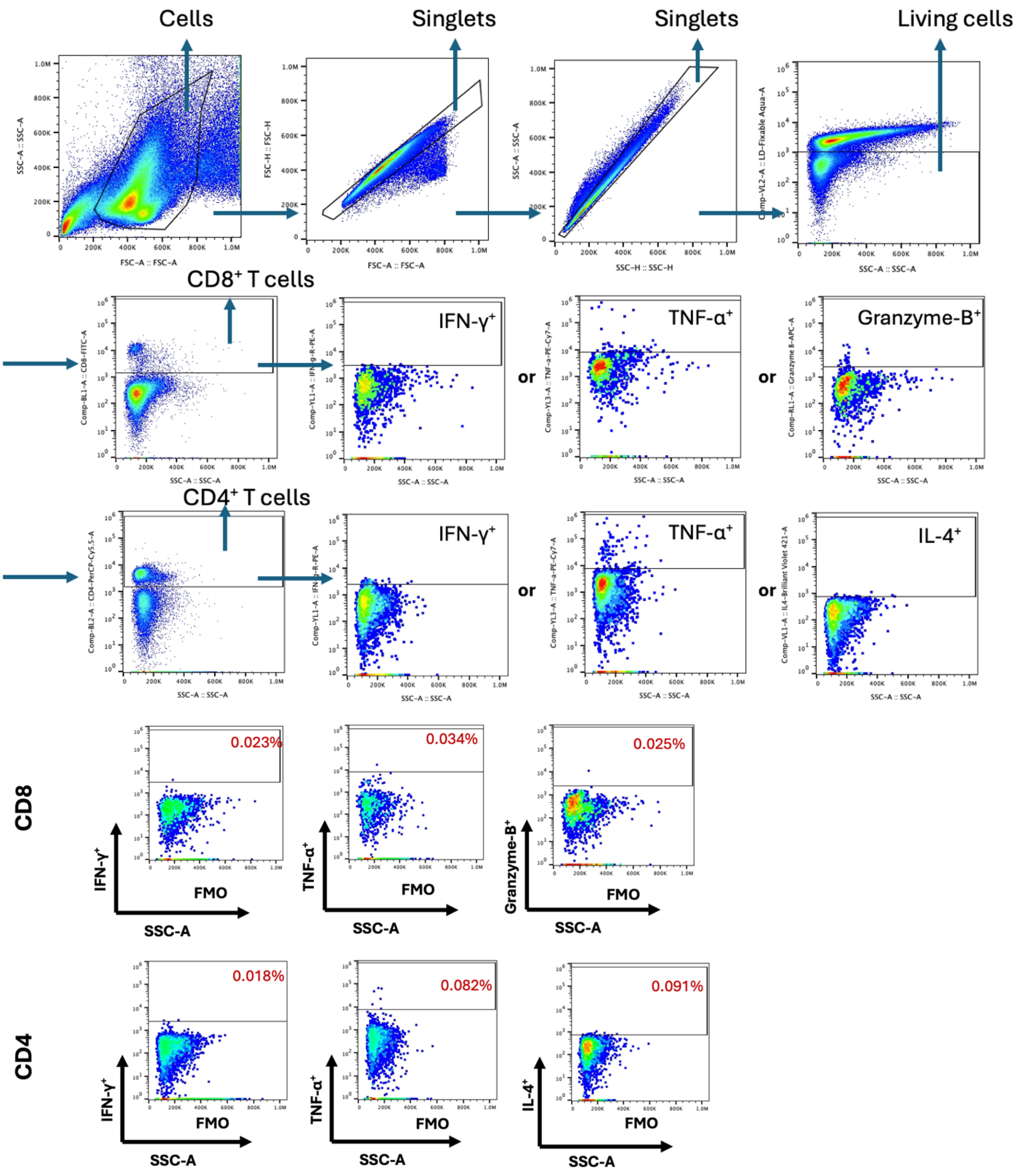


**Supplementary Figure 19. Gating strategy for flow cytometry assessments of antigen-specific immune responses generated by the LiNx formulations.** Initially, lymphocytes were gated out using SSC-A and FSC-A parameters. After gating out singlet cells using the FSC-H–FSC-A and SSC-A–SSC-H plots, viable cells were identified and gated out based on the L/D aqua-A–SSC-A plot. $CD8^+$ cells or $CD4^+$ cells were subsequently gated. Within the $CD8^+$ or $CD4^+$ T cell populations, those positive for IFN-γ, TNFα, Granzyme-B and IL-4 were identified.

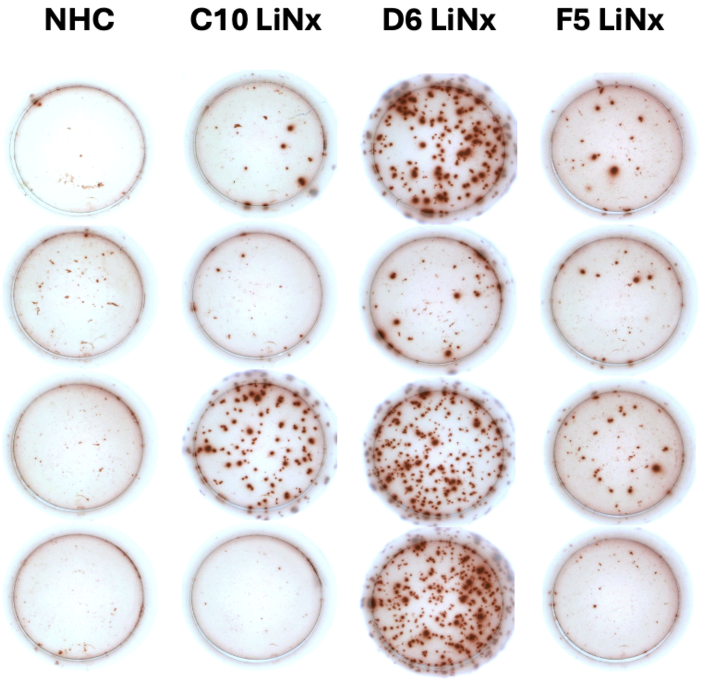


**Supplementary Figure 20. Representative images of IFN-γ secreting cells from the enzyme-linked immunospot assay.** Frequency of IFN-$\gamma$-producing cells among restimulated splenocytes, assessed via ELISPOT. Splenocytes were restimulated *in vitro* with SIINFEKL peptide (2 µg $mL^{-1}$ SIINFEKL) for 24 h.

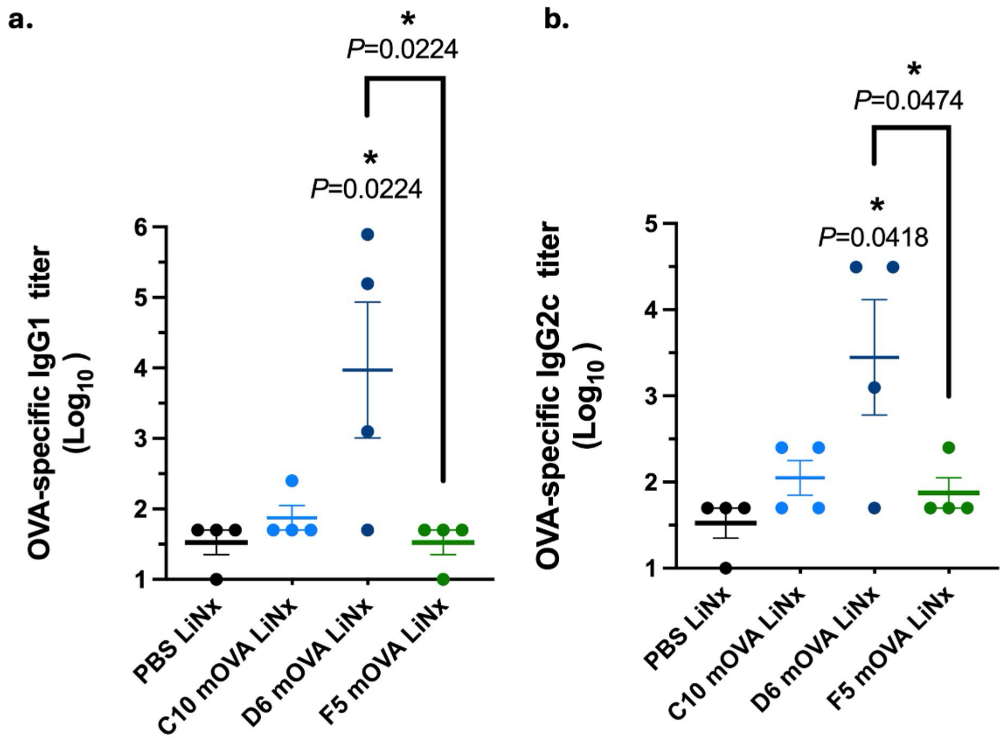


**Supplementary Figure 21. Titers of OVA-specific IgG subclass antibodies in serum samples collected on day 30 following immunization with the LiNx formulations**. IgG1 (**a**) and IgG2c (**b**) antibodies in serum on day 30 were determined by ELISA. Data represent the mean ± s.e.m. from a representative experiment (n = 4 biologically independent samples) of two independent experiments. Data were analysed using one-way ANOVA and Dunnett's multiple comparisons test. *$P$ < 0.05; ELISA, enzyme-linked immunoassay.

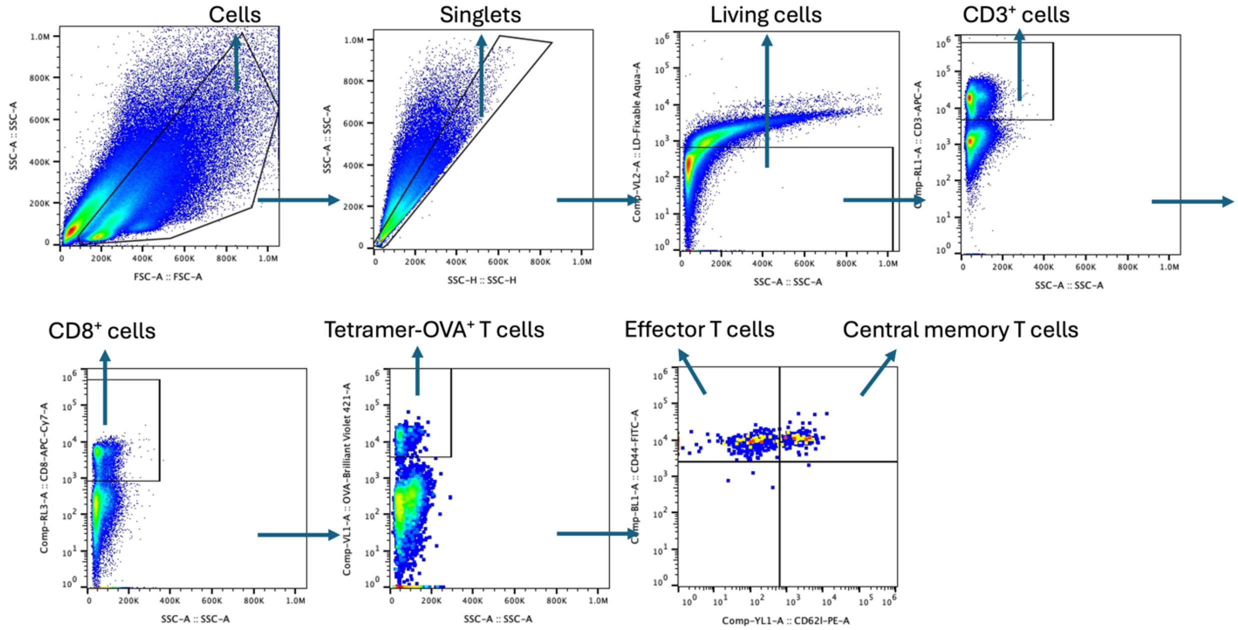


**Supplementary Figure 22. Gating strategy for flow cytometry analysis for *in vivo* assessment of OVA-specific CD8⁺ T cells by the LiNx formulations in the spleen on day 90.** Initially, lymphocytes were gated out using SSC-A and FSC-A parameters. After gating out singlet cells using the SSC-H–SSC-A plot, viable cells were further identified and gated out based on the L/D aqua-A–SSC-A plot. CD3⁺ cells were subsequently gated, and CD3⁺CD8⁺ cells were identified as CD8⁺ T cells. Within this CD8⁺ T cell population, cells positive for tetramer-OVA were defined as OVA-specific CD8⁺ T cells. Central memory T cells were defined as cells positive for both CD44 and CD62L, while effector T cells were defined as cells positive for CD44 and negative for CD62L.

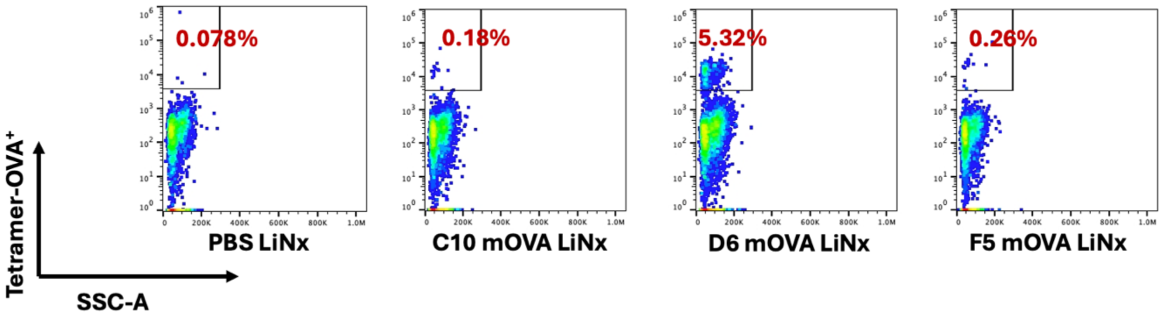


**Supplementary Figure 23. Representative flow cytometry plots for *in vivo* assessment of OVA-specific T cells in the spleen on day 90 post-administration.** C57BL/6 mice were administered with the three LiNx formulations loaded with C10, D6, and F5 mOVA LNPs via *s.c.* injections (n = 8, 30 µg mOVA per mouse). OVA-specific T cells in the spleen were analysed by flow cytometry. Percentages of cells positive for OVA tetramer are shown.

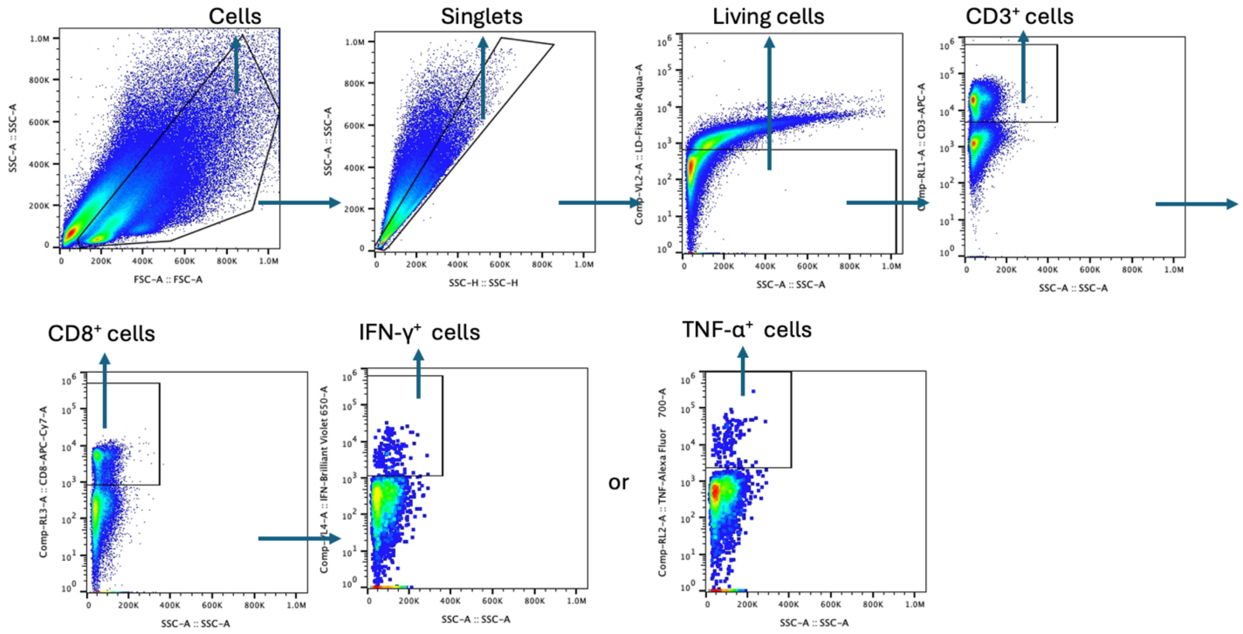


**Supplementary Figure 24. Gating strategy for flow cytometry assessment of cytotoxic T cell response on day 90 post-vaccination.** Initially, lymphocytes from the spleen were selected using SSC-A and FSC-A parameters and then singlet cells by the SSC-H–SSC-A plot. Viable cells were identified and selected based on the live/dead Fixable Aqua-A–SSC-A plot. Next, the $CD3^+$ and $CD8^+$ cell populations were selected with downstream analysis focused on IFN-γ and TNFα as illustrated in the representative figures.

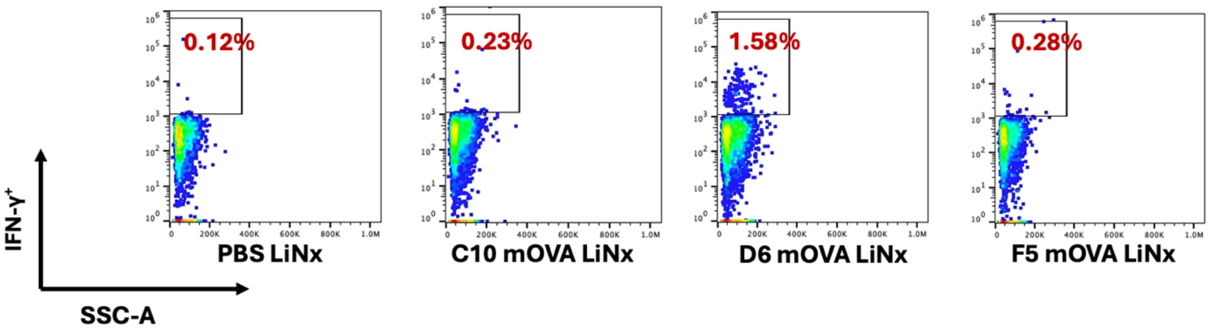


**Supplementary Figure 25. Representative flow cytometry plots for assessment of CD3⁺CD8⁺IFN-γ⁺ cells on day 90 post-vaccination.** Splenocytes from mice vaccinated with the displayed formulations were restimulated *in vitro* with OVA and SIINFEKL peptide (100 µg $mL^{-1}$ OVA and 2 µg $mL^{-1}$ SIINFEKL) for 6 h and assessed via FACS and intracellular cytokine staining to determine the percentages of $CD3^+CD8^+IFN\text{-}\gamma^+$ cells.

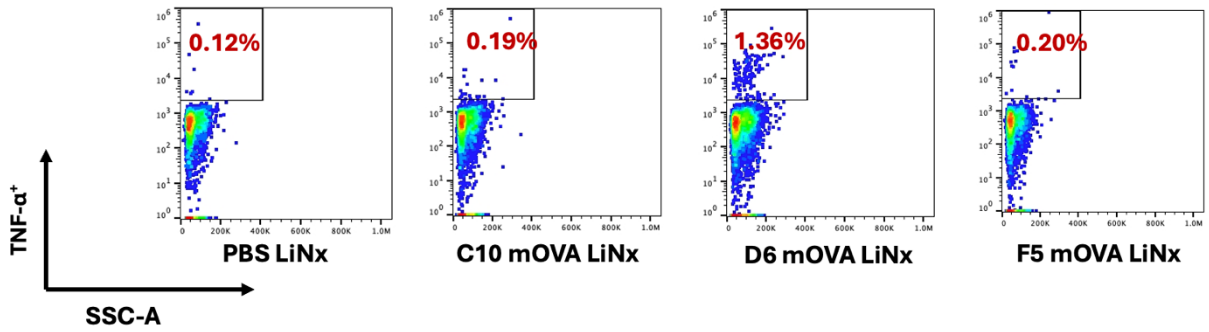


**Supplementary Figure 26. Representative flow cytometry plots for assessment of CD3$^+$CD8$^+$TNFα$^+$ cells on day 90 post-vaccination.** Splenocytes from mice vaccinated with the displayed formulations were restimulated *in vitro* with OVA and SIINFEKL peptide (100 μg $mL^{-1}$ OVA and 2 μg $mL^{-1}$ SIINFEKL) for 6 h and assessed via FACS and intracellular cytokine staining to determine the percentages of CD3$^+$CD8$^+$TNFα$^+$ cells.

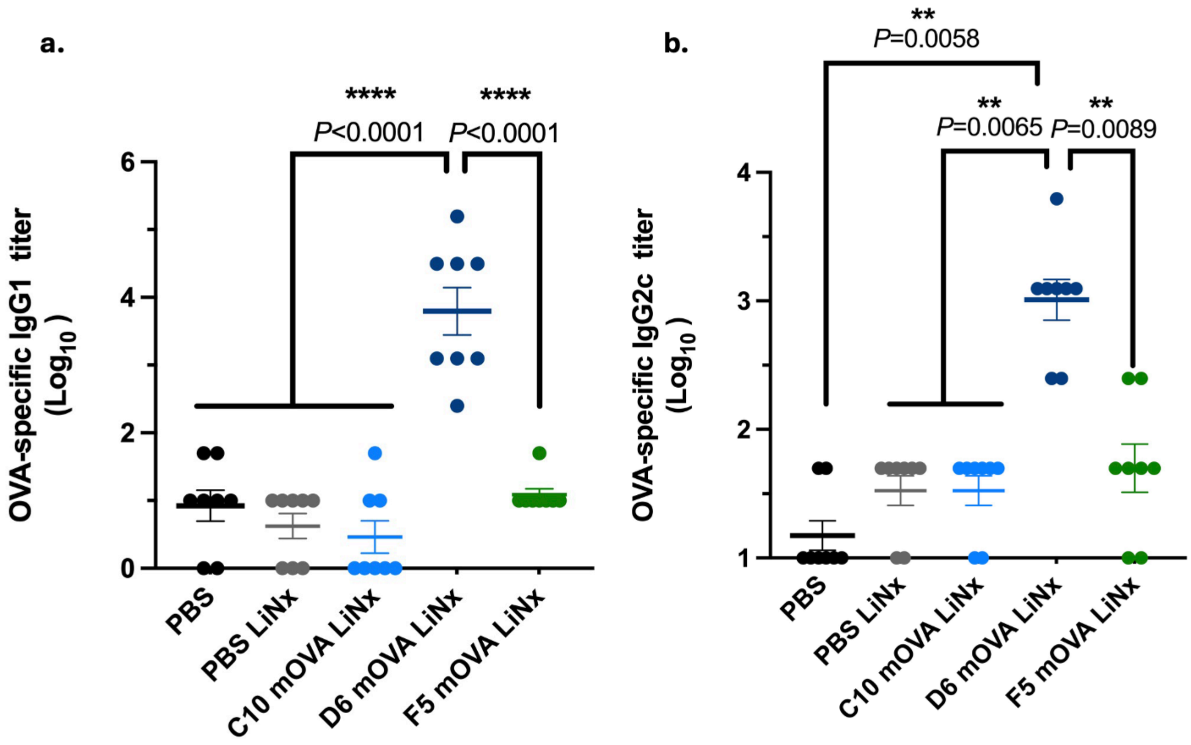


**Supplementary Figure 27. Titers of OVA-specific IgG subclass antibodies detected in serum samples collected on day 90 post-vaccination**. IgG1 (**a**) and IgG2c (**b**) antibodies in serum on day 90 were determined by ELISA. Data represent the mean ± s.e.m. with n = 8 biologically independent samples. Data were analysed using one-way ANOVA and Dunnett's multiple comparisons test. $^{**}P < 0.01$, $^{****}P < 0.0001$; ELISA, enzyme-linked immunoassay.

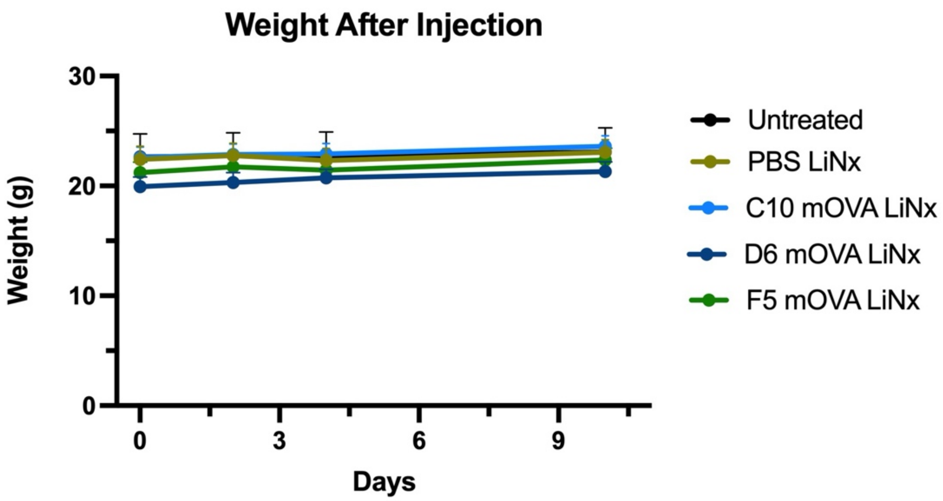


**Supplementary Figure 28. Body weights of mice vaccinated with the mOVA LiNx formulations**. The body weights of mice were monitored during vaccination schedule. Mice are given a single dosage of LiNx via *s.c.* injections on day 0 (30 μg mOVA per injection). Data represent the mean ± s.e.m. with n = 10 biologically independent samples.

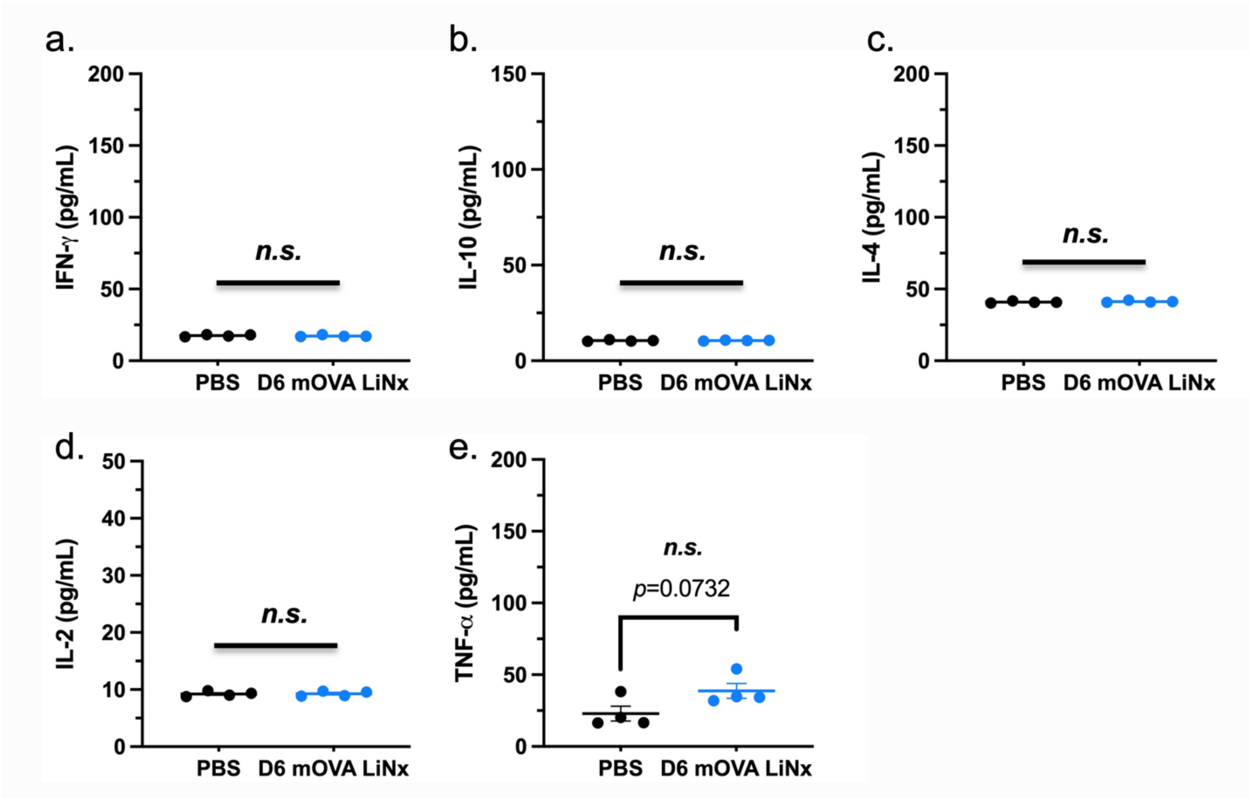


**Supplementary Figure 29. Serum cytokine levels of mice vaccinated with the D6 mOVA LiNx formulation at 24 h post-vaccination**. Serum cytokine levels, including IFN-γ **(a)**, IL-10 **(b)**, IL-4 **(c)**, IL-2 **(d)**, and TNF-α **(e)**, were measured in mice 24 hours post-vaccination with the D6 LiNx formulation. Mice received a single dose of LiNx via *s.c.* injection containing 30 μg mOVA per injection. Data represent the mean ± s.e.m. with n = 4 biologically independent samples.

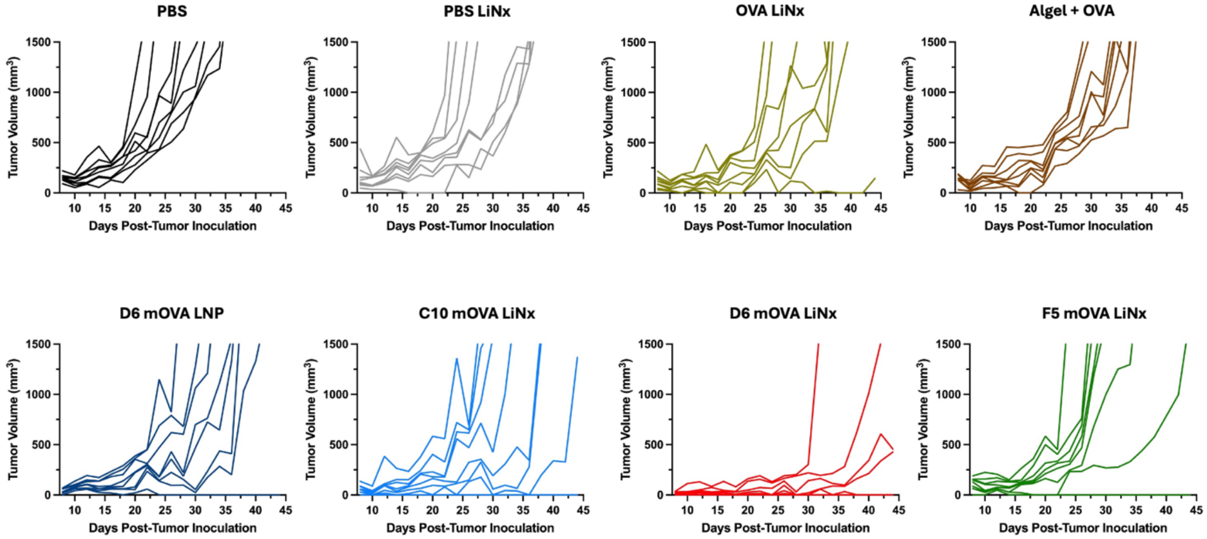


**Supplementary Figure 30. Anti-tumour efficacy of the LiNx formulations as therapeutic vaccines for MC38-OVA tumour model.** Mice were inoculated *s.c.* with MC38-OVA cells and then given three *s.c.* injections, one week apart, of mOVA-loaded D6 LNPs or LiNx (10 µg mOVA per injection) or PBS mixed with NHC. For the LiNx treatment groups, mice were administered with a single dose of LiNx loaded with mOVA (30 µg per mouse) or OVA protein (10 µg protein per mouse) via *s.c.* injection. OVA protein mixed with Alhydrogel® (1:1) (10 µg protein per mouse) served as a control group. Individual tumour volumes are shown over time.

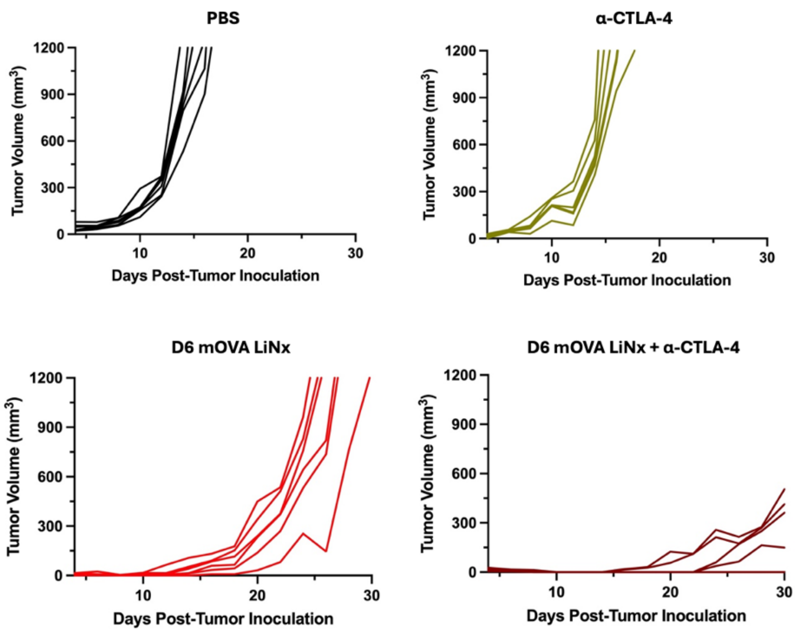


**Supplementary Figure 31. Anti-tumour efficacy of the D6 LiNx as a therapeutic vaccine for B16-OVA tumour model.** Mice were inoculated *s.c.* with B16-OVA cells and then given a single dose of D6 LiNx loaded with mOVA (30 µg per mouse) via *s.c.* injection. Two groups also received a repeated anti-CTLA-4 monoclonal antibody (mAb; 100 µg per *i.p.* injection) treatment alone or in combination with the LiNx. Individual tumour volumes are shown over time.

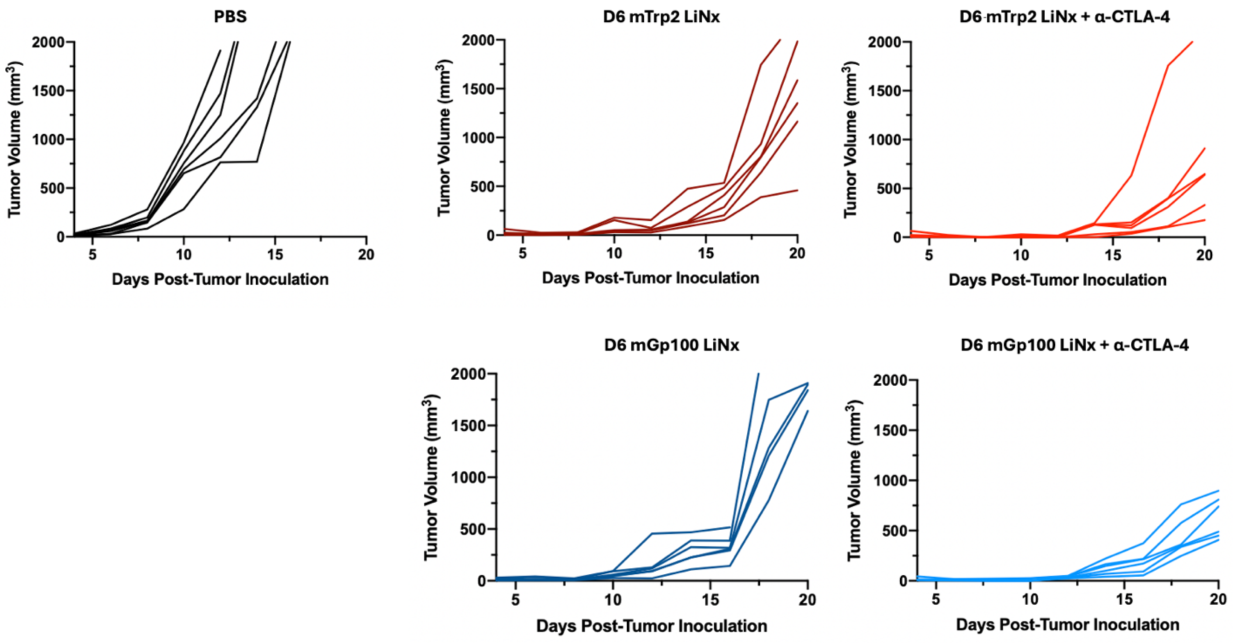


**Supplementary Figure 32. Anti-tumour efficacy of the D6 LiNx as a therapeutic vaccine for B16F10 tumour model.** Mice were inoculated *s.c.* with B16F10 cells and then given a single dose of D6 LiNx loaded with mTrp2 or mGp100 (30 µg per mouse) via *s.c.* injection. Two groups also received a repeated anti-CTLA-4 monoclonal antibody (mAb; 100 µg per *i.p.* injection) treatment in combination with the LiNx treatment. Individual tumour volumes are shown over time.

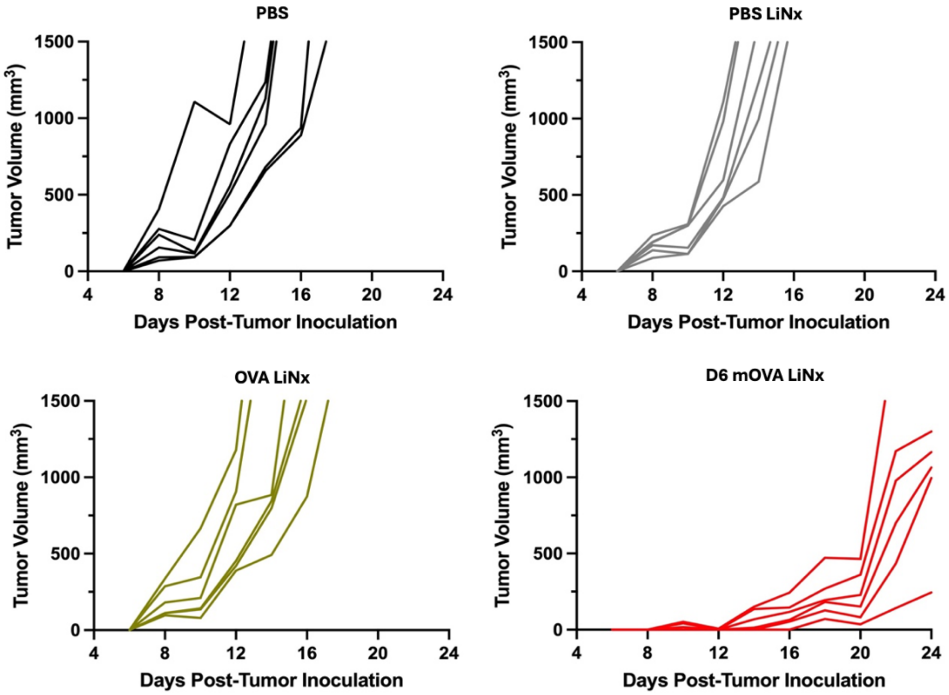


**Supplementary Figure 33. Anti-tumour efficacy of the D6 LiNx as a prophylactic vaccine for B16-OVA tumour model.** Mice were administered with a single dose of D6 LiNx loaded with mOVA (30 µg per mouse) via *s.c.* injection and then inoculated *s.c.* with B16F10-OVA cells on day 21. Individual tumour volumes are shown over time.

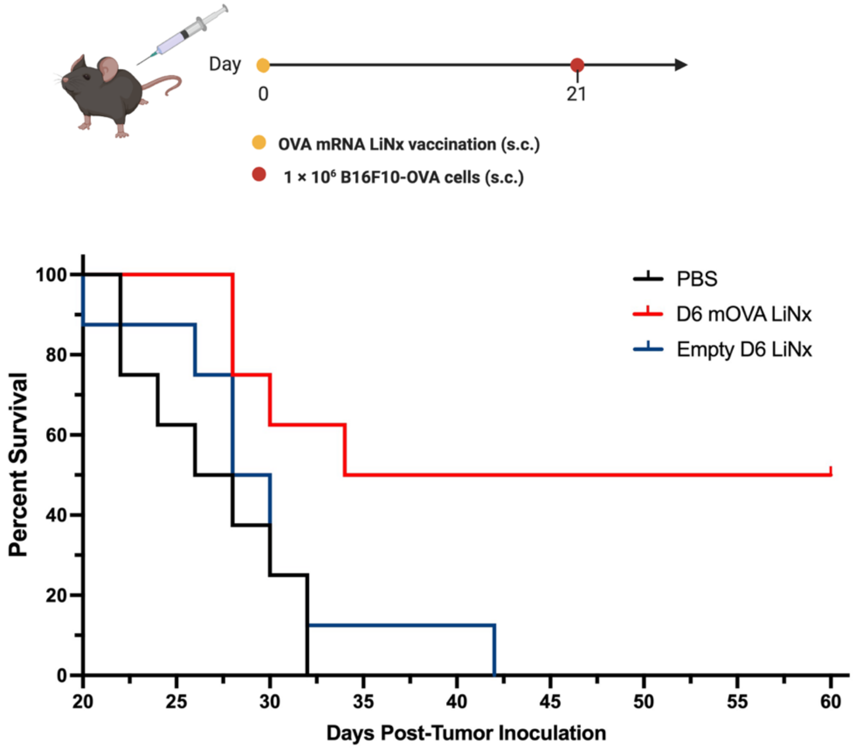


**Supplementary Figure 34. Anti-tumour efficacy of the LiNx loaded with either mOVA D6 LNPs or empty D6 LNPs in a prophylactic vaccine model for B16-OVA tumour.** Mice were administered with a single dose of D6 LiNx loaded with LNPs (30 µg mOVA per mouse) or empty D6 LNPs via *s.c.* injection and then inoculated *s.c.* with B16F10-OVA cells on day 21. Survival times are shown over time (n = 8 mice).

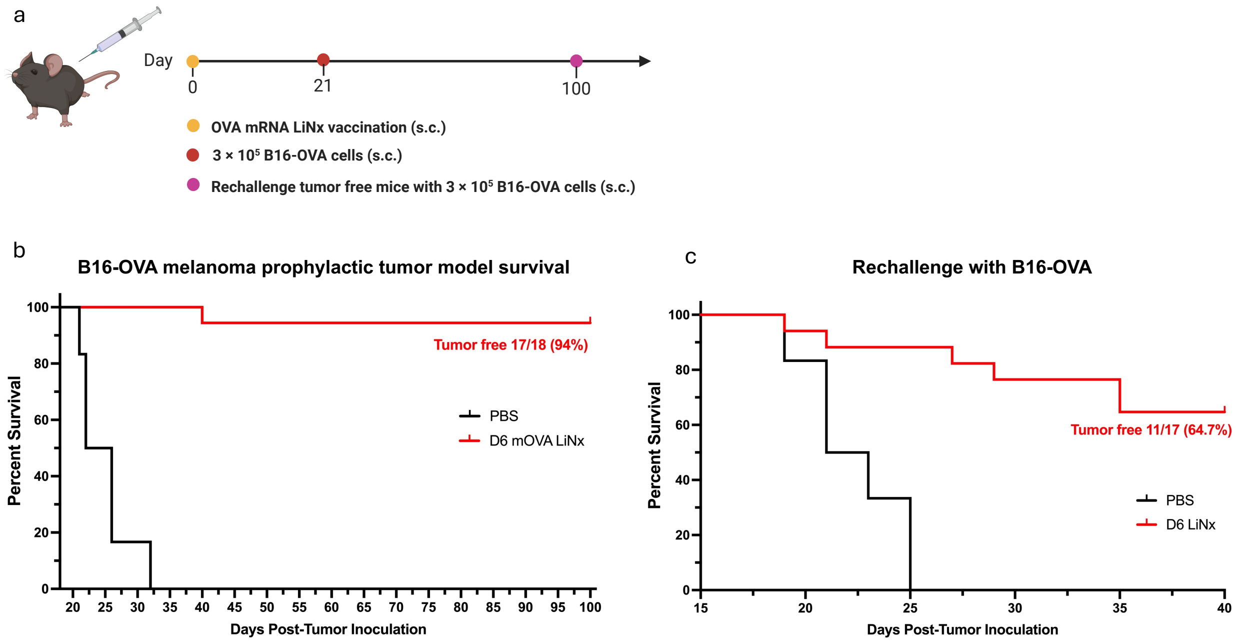


**Supplementary Figure 35. Prophylactic efficacy of the D6 LiNx in B16-OVA tumour model and resistance to rechallenge.** Schematic and results of a prophylactic vaccination and rechallenge model for B16-OVA in C57BL/6 mice. Mice were vaccinated with the D6 mOVA LiNx (30 µg per mouse) before s.c. inoculation of OVA-expressing melanoma (B16-OVA) cells. After 100 days, the tumour free mice were rechallenged with B16-OVA cells. Mice at a similar age were included as a control group injected with PBS. Survival curves (**b**, **c**) are shown. In **b**, n = 6 for PBS group and n = 18 biologically independent samples for the LiNx group. In **c**, n = 6 for PBS group, and n = 17 biologically independent samples for the LiNx group.

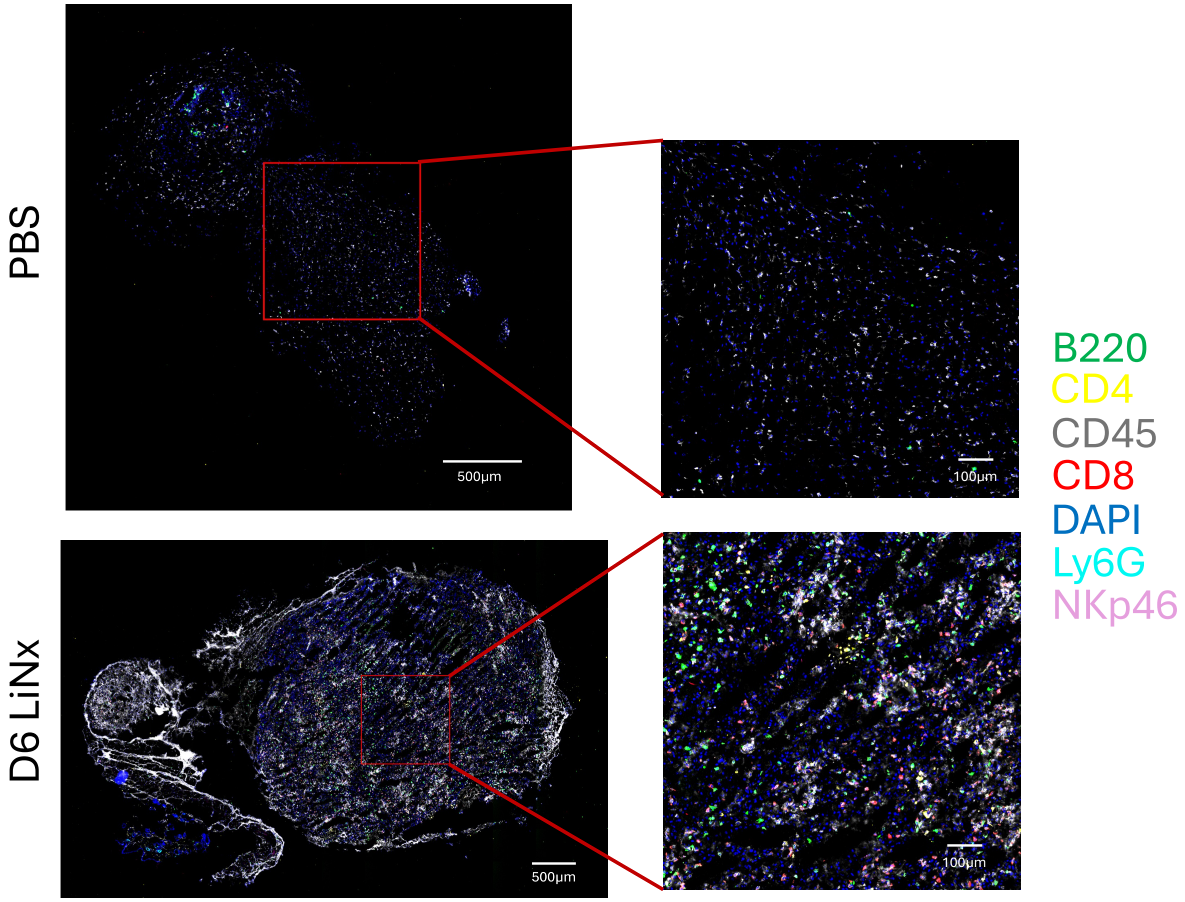


**Supplementary Figure 36. Representative images from the CODEX fluorescence imaging analysis on the collected tumour samples.** Statining details can be found in the Experimental section. Images shown here are from tumour samples stained with selected markers including B220 (green), CD4 (yellow), CD45 (white), CD8 (red), DAPI (blue), Ly6G (teal), and NKp46 (purple).

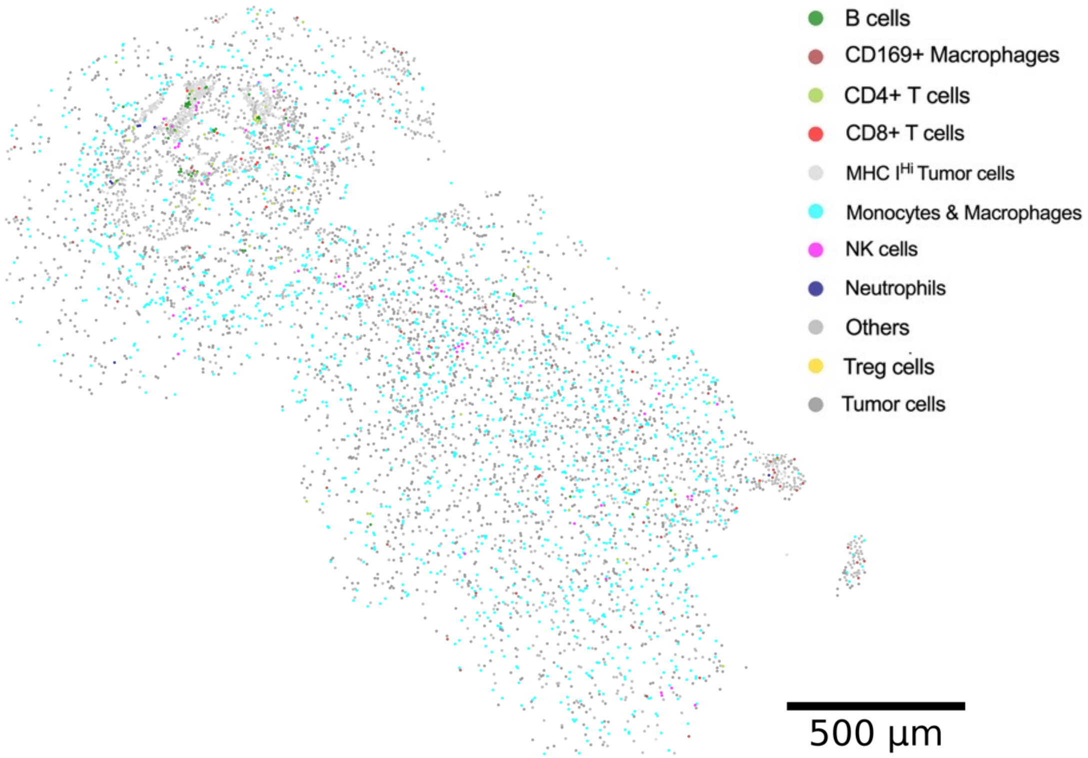


**Supplementary Figure 37. Cell type map of a representative PBS-treated tumour sample from CODEX multiplexed imaging experiment.** The coloured data points represent selected immune and tumour cell types with all other cell types lumped together in the same grey colour (scale bar = 500 µm).

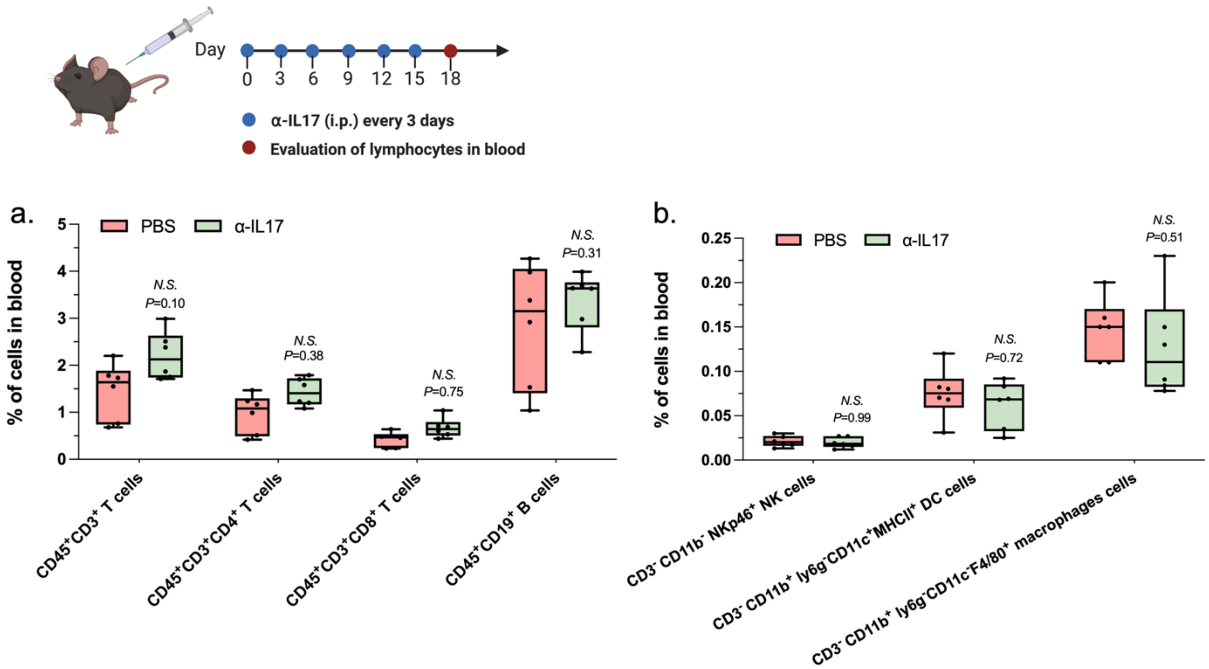


**Supplementary Figure 38. Immunocytes profile of the blood samples extracted from mice treated with or without α-IL-17 antibody.** Mice were administered with the α-IL-17 antibody (*i.p.*, 200 µg per injection every three days) over a 2-week period. Following this, blood samples were collected; and immune cell populations were analysed. (a) $CD45^+CD3^+$ T cells, $CD45^+CD3^+CD8^+$ T cells, $CD45^+CD3^+CD4^+$ T cells, and $CD45^+CD19^+$ B cells were examined. (b) $CD3^-CD11b^-NKp46^+$ NK cells, $CD3^-CD11b^+Ly6G^-CD11c^+MHCII^+$ DCs, and $CD3^-CD11b^+Ly6G^-CD11c^-F4/80^+$ macrophages were assessed. Data represent the mean ± s.e.m. (n = 4 biologically independent samples). Data were analysed using one-way ANOVA and Tukey's multiple comparisons. NS, not significant.

**Supplementary Table 1. Formulation details and particle sizes for the three selected LNPs**

| Code | Mol % | | | | N/P Ratio | Z-Average (nm) | PDI |
|---|---|---|---|---|---|---|---|
| | Helper lipid* | DLin-MC3 | Chol | DMG-PEG | | | |
| C10 | 40.00 | 40.00 | 19.96 | 0.04 | 4 | 208.9 ± 33.9 | 0.32 ± 0.06 |
| D6 | 3.64 | 36.36 | 59.41 | 0.59 | 12 | 133.0 ± 2.3 | 0.28 ± 0.02 |
| F5 | 5.45 | 54.55 | 39.92 | 0.08 | 8 | 148.6 ± 1.3 | 0.21 ± 0.04 |

* The helper lipid in C10 is 1,2-dioleoyl-*sn*-glycero-3-phosphoethanolamine (DOPE); the helper lipid in D6 is 1,2-distearoyl-*sn*-glycero-3-phosphocholine (DSPC); and the helper lipid in F5 is 1-stearoyl-2-oleoyl-sn-glycero-3-phospho-(1'-rac-glycerol) (18PG).

**Supplementary Table 2. CODEX staining conditions and cycle information**

| *Cycles* | **Ax488** | **Oligo** | **µL/ slide** | **Exp. time (ms)** | **Cy3** | **Oligo** | **µL/ slide** | **Exp. time (ms)** | **Cy5** | **Oligo** | **µL/ slide** | **Exp. time (ms)** |
|---|---|---|---|---|---|---|---|---|---|---|---|---|
| *1* | Ly6G | 55 | 0.5 | 150 | MHCII | 36 | 0.5 | 150 | FOXP3 | 72 | 0.5 | 150 |
| *2* | CD71 | 21 | 0.5 | 150 | PDPN | 74 | 2 | 150 | H2Db | 80 | 1 | 150 |
| *3* | B220 | 3 | 2 | 150 | CD8 | 6 | 0.5 | 150 | PD1 | 23 | 1 | 150 |
| *4* | SCA1 | 14 | 0.5 | 150 | keratin8 | 65 | 0.5 | 150 | NKp46 | 45 | 1 | 150 |
| *5* | Ly6C | 41 | 0.5 | 150 | CD31 | 60 | 0.5 | 150 | CD3 | 81 | 0.5 | 150 |
| *6* | CD44 | 44 | 0.5 | 150 | IgD | 79 | 0.5 | 150 | CD11c | 26 | 1 | 150 |
| *7* | 1632 | 11 | 0.5 | 150 | GRZB | 57 | 0.5 | 150 | CD4 | 7 | 1 | 150 |
| *8* | TCRb | 63 | 0.5 | 150 | F4/80 | 2 | 1 | 150 | H2Kb | 75 | 4 | 150 |
| *9* | Ki67 | 49 | 0.5 | 150 | Tim3 | 51 | 1 | 150 | CD169 | 59 | 1 | 150 |
| *10* | HDAC2 | 38 | 2 | 150 | CD19 | 24 | 1 | 150 | KLRG1 | 61 | 1 | 150 |
| *11* | aSMA | 69 | 2 | 150 | CD27 | 32 | 0.5 | 150 | CD62L | 43 | 1 | 150 |
| *12* | FAP | 15 | 2 | 150 | CD86 | 66 | 1 | 150 | CD103 | 29 | 1 | 150 |
| *13* | Blank | | | | CD90 | 17 | 0.5 | 150 | CD11b | 5 | 0.5 | 150 |
| *14* | Blank | | | | CD28 | 48 | 2 | 150 | CD45 | 20 | 1 | 150 |
| *15* | Blank | | | | CD152 | 58 | 2 | 150 | HIF-1a | 52 | 2 | 150 |
| *16* | Blank | | | | MPO | 8 | 2 | 150 | BRG1 | 42 | 2 | 150 |
| *17* | Blank | | | | CD106 | 76 | 1 | 150 | PD-L1 | 68 | 2 | 150 |
| *18* | Blank | | | | CD200 | 56 | 1 | 150 | TYRP1 | 46 | 2 | 150 |
| *19* | Blank | | | | CCR7 | 62 | 2 | 150 | CD73 | 67 | 2 | 150 |
| *20* | Blank | | | | FR4 | 71 | 2 | 75 | CD38 | 28 | 2 | 150 |
| *0* | Reference | | | 6 | Reference | | | 6 | Reference | | | 6 |

### Supplementary Table 3. CODEX marker staining evaluation

| *Cycles* | Ax488 | Evaluation | Cy3 | Evaluation | Cy5 | Evaluation |
|---|---|---|---|---|---|---|
| *1* | Ly6G | * | MHCII | * | FOXP3 | * |
| *2* | CD71 | * | PDPN | * | H2Db | * |
| *3* | B220 | * | CD8 | * | PD1 | *, NU |
| *4* | SCA1 | * | keratin8 | * | NKp46 | * |
| *5* | Ly6C | * | CD31 | * | CD3 | * |
| *6* | CD44 | * | IgD | * | CD11c | * |
| *7* | 1632 | * | GRZB | Bckg, NU | CD4 | * |
| *8* | TCRb | * | F4/80 | Neg, NU | H2Kb | * |
| *9* | Ki67 | *, NU | Tim3 | Weak, NU | CD169 | * |
| *10* | HDAC2 | *, NU | CD19 | * | KLRG1 | * |
| *11* | aSMA | * | CD27 | Weak, NU | CD62L | *, NU |
| *12* | FAP | Weak, NU | CD86 | * | CD103 | *, NU |
| *13* | Blank | | CD90 | * | CD11b | * |
| *14* | Blank | | CD28 | Neg, NU | CD45 | * |
| *15* | Blank | | CD152 | Neg, NU | HIF-1a | *, NU |
| *16* | Blank | | MPO | Neg, NU | BRG1 | *, NU |
| *17* | Blank | | CD106 | Neg, NU | PD-L1 | Neg, NU |
| *18* | Blank | | CD200 | * | TYRP1 | Neg, NU |
| *19* | Blank | | CCR7 | Neg, NU | CD73 | * |
| *20* | Blank | | FR4 | Neg, NU | CD38 | Neg, NU |
| *0* | Reference | | Reference | | Reference | |

Notes: *: Good; Neg: Not detected; Bckg: Strong levels of background signal; Weak: Weaker signal than expected; NU: Not used in clustering due to bad quality or irrelevancy.

### Supplementary Table 4. CODEX antibody information

| RRID | Company | Antibody | Clone | RRID | Company | Antibody | Clone |
|---|---|---|---|---|---|---|---|
| AB_2736987 | BioXcell | 16/32 | 2.4G2 | AB_394606 | BD | CD45 | 30-F11 |
| AB_2572996 | Invitrogen | aSMA | 1A4 | NA | SinoBiological | CD62L | 414 |
| AB_1107651 | BioXcell | B220 | Ra3-6B2 | AB_394742 | BD | CD71 | C2F2 |
| AB_3097587 | BioLegend | BRG1 | W22111B | AB_1089066 | BioLegend | CD73 | TY-11.8 |
| AB_389229 | BioLegend | CCR7 | 4B12 | AB_313144 | BioLegend | CD86 | GL-1 |
| AB_535944 | BioLegend | CD103 | 2E7 | AB_2275792 | BD | CD8a | 53-6.7 |
| AB_393741 | BD | CD106 | 429 (MVCAM.A) | AB_313168 | BioLegend | CD90 | G7 |
| AB_393577 | BD | CD11b | M1/70 | AB_2869866 | BD | F4/80 | T45-2342 |
| AB_313770 | BioLegend | CD11c | N418 | AB_2102369 | R&D | FAP | Polyclonal |
| AB_313251 | BioLegend | CD152 | UCT10-4B9 | AB_467576 | eBioscience | FOXP3 | FJK-16s |
| NA | BioRad | CD169 | MOMA-1 | AB_10828444 | Miltenyi Biotech | FR4 | TH6 |
| AB_395047 | BD | CD19 | 1D3 | NA | Novus | GRZB | AF1865 |
| AB_2687821 | BioXcell | CD200 | OX-90 | AB_397275 | BD | H-2Db | 28-14-8 |
| AB_1236456 | BioLegend | CD27 | LG.3A10 | AB_313734 | BioLegend | H-2Kb | AF6-88.5 |
| AB_1107624 | BioXcell | CD28 | 37.51 | AB_2566332 | BioLegend | HDAC2 | 13G8C67 |
| AB_395697 | BD | CD3 | 17A2 | AB_568569 | Invitrogen | HIF-1a | ESEE122 |
| AB_394815 | BD | CD31 | MEC 13.3 | AB_394858 | BD | IgD | 11-26c.2a |
| AB_2754555 | BioXcell | CD38 | NIMR5 | AB_531826 | Hybridoma | keratin8 | TROMA-1 |
| AB_393575 | BD | CD4 | RM4-5 | AB_10854564 | Invitrogen | Ki67 | SolA15 |
| AB_394645 | BD | CD44 | IM7 | AB_10949054 | BioXcell | KLRG1 | 2F1 |
| AB_1134214 | BioLegend | Ly6C | HK1.4 | AB_2291192 | BioLegend | PD-L1 | 10F.9G2 |
| AB_394206 | BD | Ly6G | 1A8 | AB_1877086 | BioLegend | PD1 | 29F.1A12 |
| AB_313316 | BioLegend | MHCII | M5/114.15.2 | AB_1089187 | BioXcell | Podoplanin | 8.1.1 |
| NA | BioXcell | MPO | 6G4 | AB_313348 | BioLegend | Sca-1 | D7 |
| AB_1727467 | BD | NKp46 | 29A1.4 | AB_10950158 | BioXcell | TCRb | H57-597 (HB218) |
| AB_10949464 | BioXcell | Tim3 | RMT3-23 | AB_10949462 | BioXcell | TYRP1 | Ta99 |
